# Human macroH2A1 drives nucleosome dephasing and genome instability in histone-humanized yeast

**DOI:** 10.1101/2023.05.06.538725

**Authors:** Max A. B. Haase, Luciana Lazar-Stefanita, Guðjón Ólafsson, Aleksandra Wudzinska, Michael J. Shen, David M. Truong, Jef D. Boeke

## Abstract

In addition to replicative histones, eukaryotic genomes encode a repertoire of non-replicative variant histones providing additional layers of structural and epigenetic regulation. Here, we systematically replaced individual replicative human histones with non-replicative human variant histones using a histone replacement system in yeast. Variants H2A.J, TsH2B, and H3.5 complemented for their respective replicative counterparts. However, macroH2A1 failed to complement and its expression was toxic in yeast, negatively interacting with native yeast histones and kinetochore genes. To isolate yeast with “macroH2A1 chromatin” we decoupled the effects of its macro and histone fold domains, which revealed that both domains sufficed to override native yeast nucleosome positioning. Furthermore, both modified constructs of macroH2A1 exhibited lower nucleosome occupancy that correlated with decreased short-range chromatin interactions (<20 Kb), disrupted centromeric clustering, and increased chromosome instability. While supporting viability, macroH2A1 dramatically alters chromatin organization in yeast, leading to genome instability and massive fitness defects.

## Introduction

The basic repeating unit of eukaryotic chromatin is the nucleosome core particle^1^. This is defined as approximately 146 bp of DNA wrap around a histone octamer that is comprised of a tetramer of histone H3 and H4 and two dimers of histones H2A and H2B^2^. Replicative histones package the bulk of DNA, are regulated in a cell-cycle specific manner, and are typically encoded in multicopy gene clusters^3^. The conserved role of replicative histones in DNA packaging and regulation is apparent by their high sequence identity in divergent species (for example when comparing yeast to human)^4, 5^. In contrast, non-replicative variant histones are typically encoded by distinct genes, separated from the replicative histone clusters, and as the name suggests, regulated independently of the cell cycle^6^. Moreover, variant histones typically have selective chromatin deposition/eviction mechanisms linked to specific chromatin remodelers and chaperones^7, 8^. Certain histone variants are considered ‘universal’ as they diverged prior to the diversification of eukaryotes (CenH3, H3.3, H2A.Z and H2A.X) and are broadly found in most species, reflecting their essential functions in ancient processes such as CenH3 in maintaining centromeric chromatin for chromosome segregation^9^. In contrast, some ancient histone variants have been differentially lost throughout evolution, such macroH2A in fungi, which evolved long ago in premetazoan protists prior to the divergence of metazoans and fungi and was lost in the latter^10^. Moreover, histone variants have continually emerged throughout evolution, through gene duplication as in the case of macroH2A2 in the basal roots of vertebrate evolution^11^ or via duplication and rapid diversification of short H2As in eutherian mammals^12^. Budding yeasts’ have a surprisingly small complement of variant histones, especially in contrast to a species such as humans (Figure 1A). The budding yeast *Saccharomyces cerevisiae* minimally encodes a centromeric-specific H3 (Cse4), which defines its point centromeres, a H2A.Z variant (Htz1) that localizes to either side of the nucleosome depleted region (NDR) near transcription start sites (TSS) and a histone H1 variant, Hho1, which plays specific roles in the compaction of chromatin during sporulation^13–15^.

**Figure 1.**
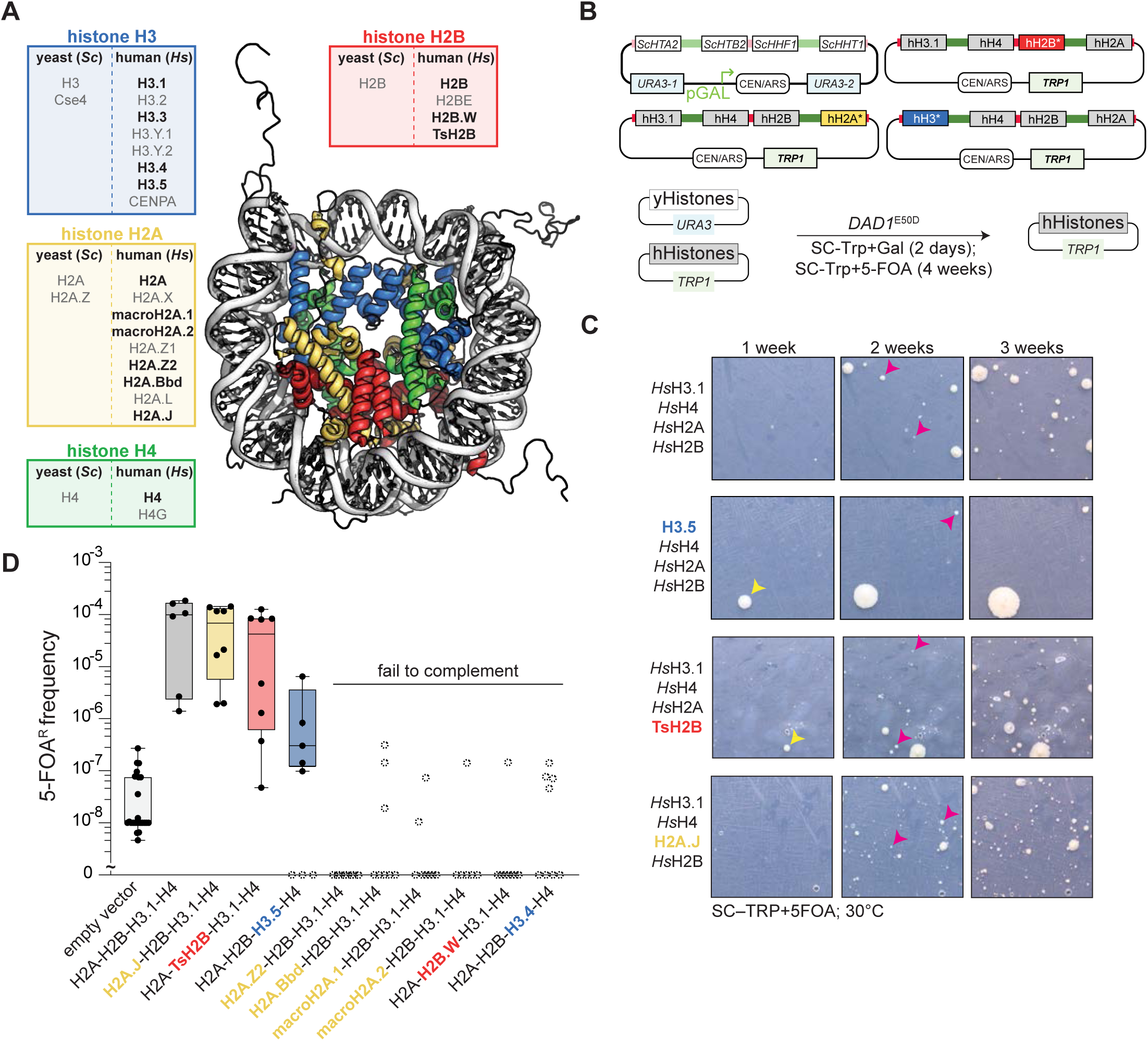
Complementation of human replicative histones with their variant histone counterparts in yeast. **(A)** Overview of human histone variants examined in this study (bolded) and remaining variants not studied. Color-coded nucleosome core particle is shown (1KX5). **(B)** Overview histone humanization assay (see methods for details). **(C)** Exemplar images of histone humanization assay at three time points (1 week, 2 weeks, and 3 weeks growth at 30°C). Yellow arrows denote large colonies that emerged early (within 1 week of growth) and pink arrows denote small colonies which emerged around 2 weeks of growth. **(D)** Humanization assay for single histone variant swaps. The background of the assay is determined by the empty vector swap, in which plasmid recombinants or spontaneous *ura3* mutants bypass 5-FOA selection at an average rate of ∼1 in 10 million cells. Histone variants are colored coded as in panel A. Open dashed-line circles indicate failure to isolate true humanized clones as assessed by PCR genotyping (Figure S1).

Histone variant incorporation into chromatin serves as an additional layer of regulation of chromatin structure and function^7^. For example, the variant macroH2A1 encodes a C-terminal macro domain approximately twice the size of its histone fold domain^16^. *In vitro* the macroH2A1 histone fold preferentially makes heterotypic nucleosomes with replicative H2A and resists chromatin remodeling by reducing the recruitment of the ATP-dependent chromatin remodeler, SWI/SNF^17, 18^. Moreover, macroH2A1 is enriched at transcriptionally silenced chromatin, directly inhibiting the recruitment of RNA polymerase II, chromatin remodelers and transcription factors^6, 19, 20^.

Nucleosomes are organized into phased arrays with a characteristic spacing between them, termed the nucleosome repeat length (NRL)^21, 22^. Nucleosome phasing is typically set against genomic barriers nearest to transcription start sites (TSS), defined by a nucleosome depleted region (NDR) and the precise positioning of the first downstream nucleosome (NDR +1 nucleosomes) which are critical in transcriptional regulation^23^. The complete nucleosome landscape is set by a multitude of interacting protein complexes and underlying DNA sequence/mechanics^24, 25^. In yeasts, the phased landscape near the TSS is largely determined by the action of ATP-dependent chromatin remodelers, which counteract nucleosome-disruptive processes such as transcription, DNA replication and repair^22, 24^. The combined action of RSC and INO80 remodelers precisely set the +1-nucleosome positioning in yeasts, establishing spacing near genomic barriers such as Reb1 binding sites^24–26^. Internucleosomal distance is independent of nucleosome density both *in vivo* and *in vitro*^5, 27–29^. Additionally, factors such as IWS1a, ISW1b, or Chd1 further refine nucleosome spacing to the characteristic NRL observed in wildtype (WT) cells.

*In vitro* chromatin reconstitution using replicative histones has proven to be a powerful probe of structural and functional effects from the bottom-up. However, these systems lack cellular processes such as transcription or DNA replication. On the other hand, due to their restricted deposition, the constant presence of replicative histones, and a potential multitude of interacting epigenetic states, direct study of a living genome chromatinized exclusively by variant histones has been limited. Remarkably, despite over ∼1 billion years of divergent evolution, the replicative histones of yeast could be entirely exchanged with human replicative histones^5, 30, 31^. These histone-humanized yeasts provide a powerful “*in vivo* reconstitution” system of human chromatin as we can “reset” the composition of the DNA packaging in yeast. Here, we adapted the histone-replacement system to directly test complementation of the majority of human variant histones for their corresponding replicative histones (e.g., does H2A.J substitute for replicative H2A?). We defined a set of human variant histones which can indeed fully complement their replicative counterparts in yeast (H2A.J, TsH2B, and H3.5). Moreover, we simultaneously replaced replicative H3, H2A, and H2B with human H3.5, H2A.J, and TsH2B, demonstrating that these three variant histones functional replace replicative histone in a living cell. We then focused on dissecting the incompatibility of the human variant histone macroH2A1 with yeast chromatin and systematically determined which residues are inviable in yeast. Doing so allowed us to decouple separate effects of the macro domain and histone fold domain, the latter being responsible for the fundamental incompatibility with yeast chromatin. Using both MNase-seq and HiC assays, we show that humanized yeast in which yeast-compatible versions of macroH2A1 replaces replicative H2A exhibit surprising structural and functional alterations to their chromatin alongside enhanced genome instability. Thus, while yeast may have never before packaged their genomes with these particular variant histones, it serves as a powerful system to study impact of chromatin from divergent species.

## Results

### Humanization of yeast chromatin with non-replicative human histone variants

Yeast can use either the human replicative histone *Hs*H3.1 or the variant *Hs*H3.3^5, 32^ with a preference for *Hs*H3.1^5^ (for clarity, replicative human histones are explicitly written with a preceding “*Hs*” and yeast histones with a preceding “S*c*”). However, it is not known whether life is sustainable when the genome is packaged entirely with other non-replicative histone variants beyond *Hs*H3.3. To address this, we adapted the dual-plasmid histone shuffling method for exchanging variant histones for replicative histones in yeast (Figure 1B)^5, 30, 31^.

We made plasmids in which a single replicative human histone gene is replaced by a variant type and used these with our histone shuffle strain to test for complementation (Figure 1B). We failed to isolate humanized clones for the majority of the histone variants (H2A.Z2, H2A.Bbd, macroH2A1, macroH2A2, H3.4, H2B.W), consistent with the idea these histone variants lack essential functions, typically executed by replicative histones, needed for packaging bulk DNA. For example, the variant macroH2A1 produced only 1 clone that appeared after two weeks of growth, however, genotyping of this clone revealed it contained yeast histone genes (Figure S1). In contrast, we readily isolated true histone humanized clones for variants *Hs*H2A.J (71% identical [amino acid sequence identity] to yeast H2A), *Hs*TsH2B (63% identical to yeast H2B), and *Hs*H3.5 (86% identical to yeast H3; Figure 1C–D; S1). Humanized clones were validated by genotyping colonies that appeared 2 weeks post plating to 5-FOA, which selects for cells that lost the yeast histone plasmid (Figure S1). As these variants could complement their replicative counterparts individually, we tested whether all three, together, could simultaneously replace replicative H3, H2A, and H2B. Remarkably, *Hs*H3.5, *Hs*H2A.J, and *Hs*TsH2B simultaneously replaced replicative H3, H2A, and H2B, respectively (Figure S2A). These data suggest that *Hs*H2A.J, *Hs*TsH2B, and *Hs*H3.5, the latter two of which are testis specific^33, 34^, all retain the essential functions of yeast replicative histones.

The above data show that histone variant *Hs*H3.5 humanized less frequently than replicative *Hs*H3.1 whereas variant *Hs*H3.4 (*Hs*H3T) failed to humanize all together (Figure 1D). These two variants form unstable nucleosomes *in vitro* and in both cases this instability is attributable to single amino acid substitutions^33, 35^. We swapped in the replicative *Hs*H3.1 residues known to act as nucleosome stabilizing mutations into *Hs*H3.5 (L103F) and *Hs*H3.4 (V111A) and tested for complementation. The stabilizing mutations improved humanization of both *Hs*H3.5 (>100-fold) and *Hs*H3.4 (>400-fold), thus fully complementing for *Hs*H3.1 (Figure S2C). Additionally, *Hs*H3.5 lacks two conserved lysine residues K36 and K79 that are modified by lysine methyltransferases Set2 and Dot1, respectively. Introduction of the two lysine residues into *Hs*H3.5 improved humanization over ∼27-fold (in the absence of the stabilizing L103F mutation), suggesting that, in addition to improving nucleosome stability, restoring the two modifiable lysine residues of histone *Hs*H3.5 is critical for proper histone H3 function in yeast.

### Human macroH2A1 is a dominant negative histone variant in *S. cerevisiae*

We next sought to understand why macroH2A1 failed to replace replicative *Hs*H2A. We confirmed that macroH2A1 is expressed in wildtype yeast by immunoblot of a GFP-tagged macroH2A1 and observed that it is correctly localized to the nucleus (Figure 2A–B). Inducible expression of macroH2A1 resulted in a growth defect in wild-type yeasts (Figure 2C). Using a genome-wide deletion screen of the non-essential yeast genes, we explored genetic interactions (GIs) with macroH2A1 overexpression (Figure S3A). We identified numerous synthetic sick GIs (z-score normalized > 2; Table S4) that were enriched in GO cellular components such as the COMA complex (*MCM21* and *CTF19*), Kinetochore (*IML3*, *PAT1*, *MCM21*, *SLX8*, *MCM22*, *CTF3*, *CTF19*), Nucleosome (*HTB2*, *HHF1*, *HTZ1*), and Organellar large ribosomal subunit (*MRPL36*, *MHR1*, *MRP49*, *MRPL20*, *MRPL10*). Positive genetic interactions or “suppressors of macroH2A1 expression” were enriched for genes within molecular complexes such as the NELF complex (*DST1*, *SPT4*), Bfa1-Bub2 complex (*BFA1*, *BUB2*), Ribosome (*RPS11A*, *RPS6A*, *IMG1*, *MRP10*, *CBS2*, *RPS27B*, *RPS29A*, *RPS10A*, *MRP7*, *MRPL10*, *RPS19B*), and Mitochondrial ATP synthase complex (*ATP1*, *ATP2*; Table S4). These data suggest that macroH2A1 interferes with a broad variety of processes such as centromere-kinetochore function and the metabolism of mitochondria and ribosomes (Figure S3B). Some of the top synthetic sick hits corresponded to the genes encoding yeast histones (Figure 2D, S3C), suggesting that their reduced dosage exacerbates the fitness defect of macroH2A1 in wildtype yeast. Moreover, the toxicity of macroH2A1 was not rescued by the deletion of yeast’s native H2A.Z remodeler, Swr1, but was rescued by introduction of two mutations (I100T and S102P) in the C-terminal region of macroH2A1, predicted to disrupt H2A’s chromatin association^37^ (Figure S4).

**Figure 2.**
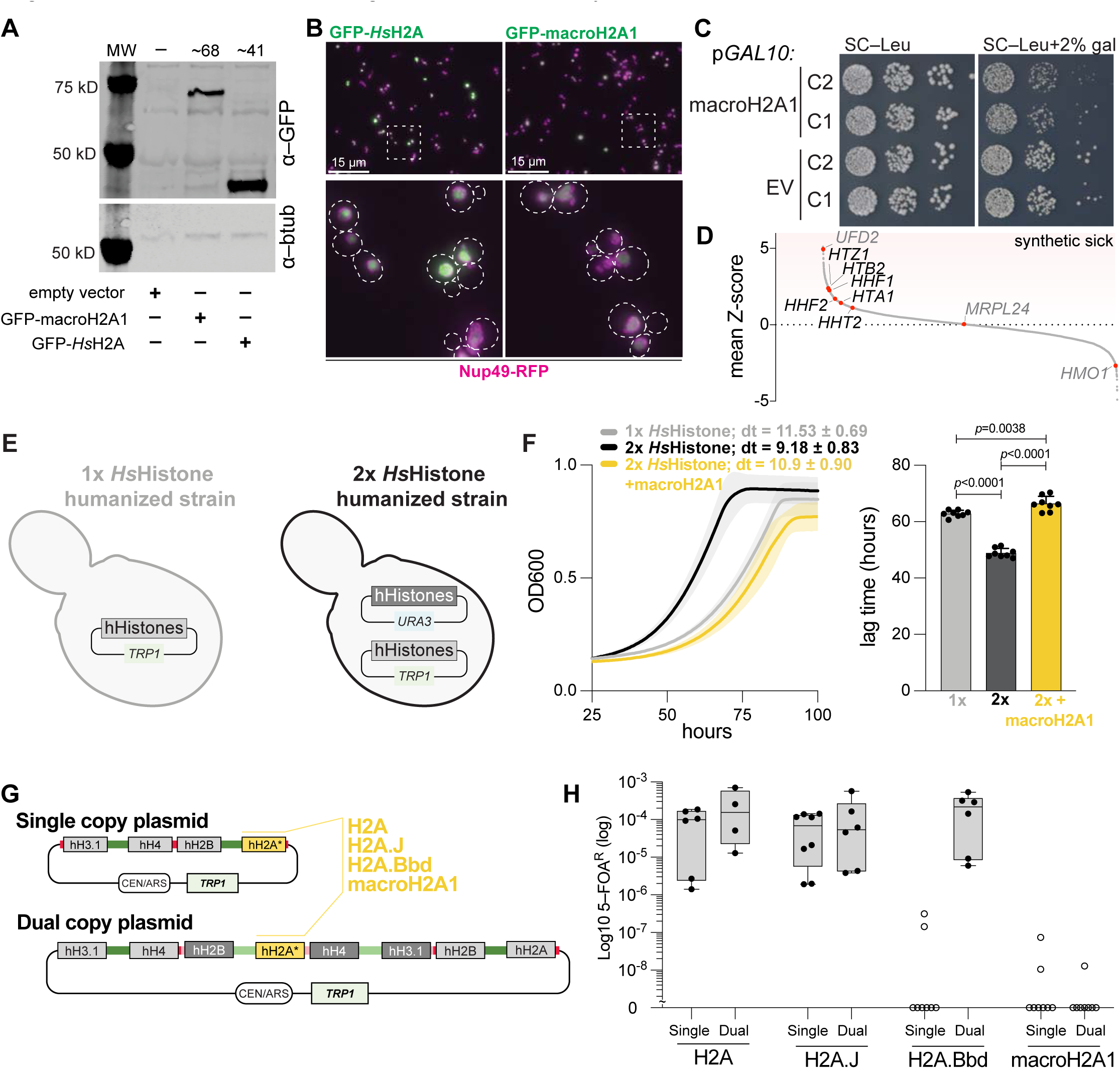
macroH2A1 is a dominant negative histone variant in yeast. **(A)** Western blot analysis of histone expression in wild-type cells. Blotting was done using a dual-color secondary antibody approach and each channel is shown separately. **(B)** GFP-macroH2A1 correctly localizes to the nucleus in wild type cells. Cells with an RFP-tagged nuclear envelope protein (Nup49-RFP) were transformed with GFP-H2A fusions as labeled and then imaged from mid-log phase cultures. **(C)** Overexpression of macroH2A1 in wildtype cells is toxic. Cells with the indicated plasmid were grown in the presence of glucose (no expression) or galactose (over expression). **(D)** Genetic interactions screen of non-essential gene deletions with macroH2A1 overexpression. Histone genes are highlighted as are three example genes which showed negative, no, and positive GIs with macroH2A1 overexpression. **(E)** Schematic of strains used in growth assays. 1x histone humanized strain has a single copy of each human replicative histone on a *TRP1* CEN/ARS plasmid (pDT109). 2x histone humanized strain has two sets of human histones genes (wither all replicative (pDT109 + pMAH022) or with one non-replicate variant (See Table S2)) on two CEN/ARS plasmids. **(F)** Expression of macroH2A1 is toxic in histone humanized cells. Growth assays of 1x (gray) and 2x histone humanized yeast (with replicative *Hs*H2A only (black; pDT109 + pMAH022) or replicative *Hs*H2A with macroH2A1(yellow; pDT109 + pMAH87)). Left; is A_600_ growth curves. Right; calculated lag times. **(G)** Schematic of single-copy (pDT109) and dual-copy (pMAH342) human histone expression vectors. H2A variants can be cloned into the site colored yellow (pMAH345). Promoters are from the native histone cluster loci, dark green *HTA1B1* and *HHF2T2*; light green *HTA2B2* and *HHF1T1*. The yeast histones are derived from the histone loci of *S. eubayanus* and are encoded on the Superloser plasmid (Figure S4 and S5; pMAH316). **(H)** Humanization rates for the various human variant histones in either the absence (single) or presence (dual) of a second set of replicative human histones.

To test whether macroH2A1 is also toxic in the human chromatin background, we co-expressed macroH2A1 in an already histone humanized strain (Figure 2E). Briefly, we transformed a previously humanized strain (with all four replicative human histones encoded on a *TRP1* CEN/ARS plasmid) with a *URA3* CEN/ARS plasmid encoding a second set of human histone variants (either all replicative histones or 3 replicative histones + 1 variant histone). We assayed growth using a high-throughput plate reader and found that the strain with two plasmids encoding replicative human histones (2x hHistones) grew significantly better than the parental strain with a single plasmid (1x hHistones; Figure 2F). We extrapolated a doubling time of 9.18 ± 0.83 hours and a lag time of 48 ± 1.6 hours in the 2x hHistone strain, compared to a doubling time of 11.53 ± 0.69 hours and a lag time of 62 ± 1.2 hours in the 1x hHistone strain. In contrast, the co-expression of macroH2A1 in the 2x hHistone strain significantly slowed its doubling time to 10.9 ± 0.90 hours and increased the lag time to 66.4 ± 2.6 hours (Figure 2F). Critically, this was not due to a gene dosage effect of *Hs*H2A, as the 2x hHistone strain with a single *Hs*H2A gene grew as well as normal 2x hHistone (Figure S4E).

We next tested whether we could isolate histone humanize yeasts with replicative and non-replicative variant histones simultaneously present on the same plasmid using an improved host strain/plasmid configuration (Dual copy plasmid shuffle; Figure 2G). The dual–copy histone shuffle strain has two advantages: 1) after shuffling, it is healthier with two sets of human histone genes than the original system which had a single set and 2) It allows the incorporation of variant histones either in the presence or absence of the corresponding human core histone gene. To minimize recombination between the two plasmids, the yeast histone plasmid encodes a single set of each histone gene cluster from the related species *S. eubayanus* (see methods; Figure 2G & S5). We tested all histone variants in this system (Figure S6B), but for simplicity describe only the results for three H2A variants, H2A.J, H2A.Bbd, and macroH2A1; the latter two being inviable in the 1:1 replacement of replicative H2A (Figure 1D). When we humanized H2A.Bbd in the presence of replicative *Hs*H2A we observed robust isolation of humanized colonies (Figure 2H and Figure S6B), suggesting that H2A.Bbd is either not incorporated into chromatin or lacks essential nucleosome functions in yeast. However, when the same was done with macroH2A1 we failed to observe any histone humanized colonies (Figure 2H and Figure S6B). Collectively our genetic interaction data, co-expression experiments, and histone humanizations suggest that macroH2A1 is incorporated into the chromatin of *S. cerevisiae,* where it may disrupt the structure and function of the chromatin to such a point that viability is lost.

### The histone fold domain of macroH2A1 negatively affects yeast viability

We next set out to map the regions of macroH2A1 that contribute to *S. cerevisiae* growth arrest. All following experiments were performed using the single copy plasmid system with replicative *Hs*H2A replaced with macroH2A1 mutants or chimeric constructs. We first generated a chimeric construct with the macro domain of macroH2A1 grafted to replicative *Hs*H2A and tested for viability after histone shuffling. This chimeric *Hs*H2A–macro-domain led to histone humanized yeast (albeit at a significantly reduced frequency from replicative *Hs*H2A). We denote this as “H2Amacro1” (color-coded in yellow in Figure 3A). Based on these results, we reasoned that the inviability of macroH2A1 maps to its histone fold domain (HFD).

**Figure 3.**
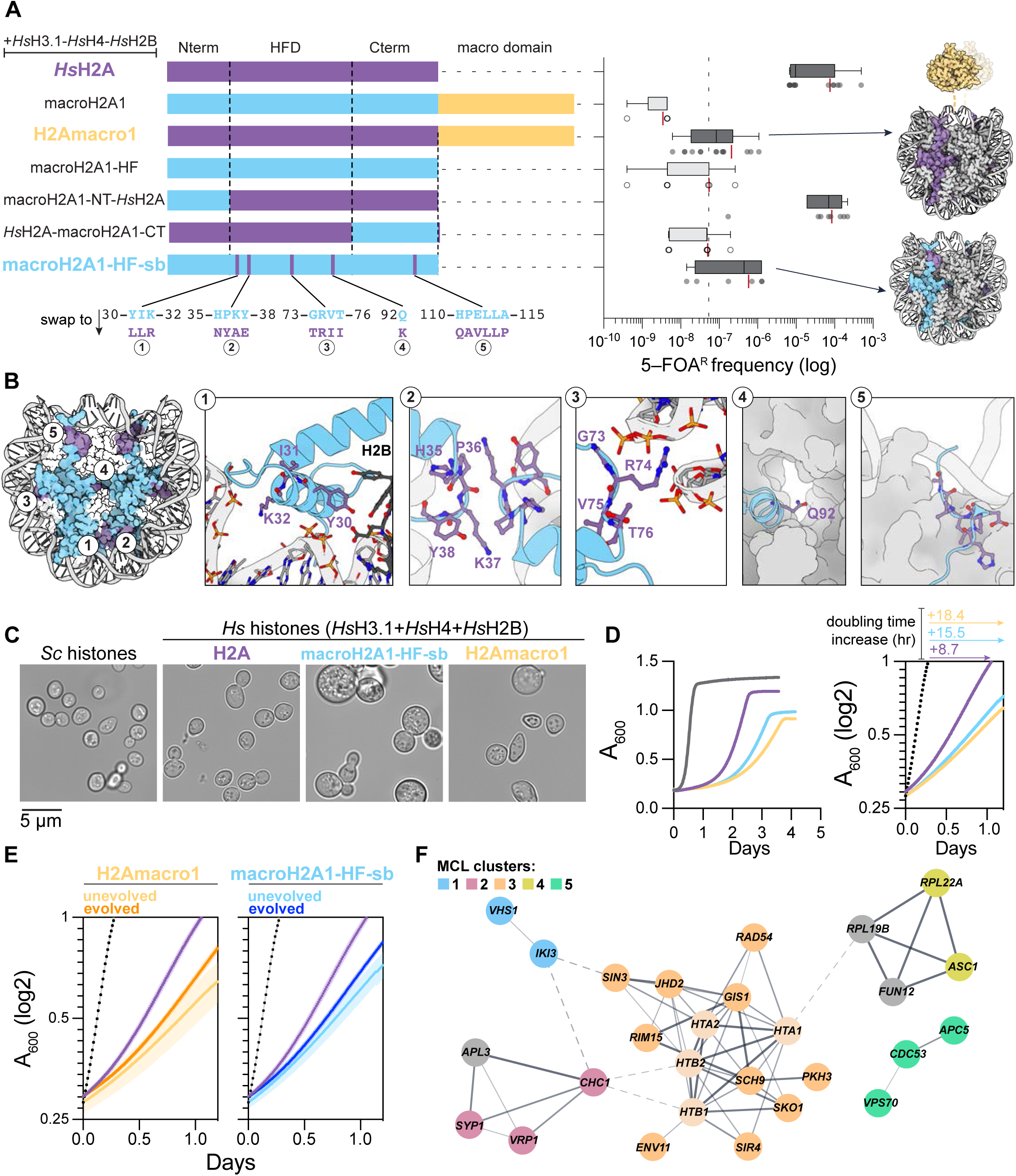
The histone fold of macroH2A1, and not the macro domain, causes yeast inviability. **(A)** Humanization of yeast with macroH2A1 chromatin. Right; schematic of replicative *HS*H2A-macroH2A1 chimeras. Details of macroH2A1 histone fold swap-back experiments are found in Supplemental Figure 7. Left; humanization assay of replicative *Hs*H2A, macroH2A1, and the chimeras. The swap-back details are displayed below the macroH2A1-HF-sb construct. Open circles indicate that the 5-FOA^R^ colonies isolated retained the yeast histones. Boxes represent the median with 25^th^ to 75^th^ percentiles, with whiskers extending to the 5^th^ to 95^th^ percentiles. Dots underneath represent each replicate with red lines representing the mean 5-FOA^R^ frequency. Dashed line at ∼10^-7^ represents the average background frequency of isolating spontaneous *ura3* mutants in our shuffle assay (based on shuffling out the yeast histone plasmid for an incoming plasmid missing a complement for either H3, H2A, or H2B). To the right are schematic views of the H2Amacro1 (macro domain is fused to *Hs*H2A) and the macroH2A1-HF-sb constructs. **(B)** View of the human macroH2A1 nucleosome (PDB: 1U35) with the residues which were swapped back to the corresponding *Hs*H2A residue highlighted in purple. Five zoomed in views showing the details of the swap-back residues, note that the native macroH2A1 residues are shown; numbering corresponds to macroH2A1 (See Methods for details). **(C)** Exemplar images of WT and humanized cells as indicated. Scale bar 5 µm. Images were acquired from log-phase cultures. **(D)** Growth assay of yeast strains. Right; strains (colored as in panel A; gray is the strain with *Sc* histones) were grown in YPD in a 96-well clear bottom plate format and absorbance (A_600_) was measured every 15 minutes for up to 5 days. Line represents the average of at least six biological replicates. Left; absorbance (A_600_) plotted to a log2 scale during the logarithmic growth phase of each strain. Time was set to zero at the calculated end of the lag phase for all growth curves. **(E)** Logarithmic growth curves for the unevolved and evolved H2Amacro1 (left) and macroH2A1-HF-sb (right) histone humanized strains. Growth curves of the *Sc* histone strain (gray) and the *Hs* histone strain (purple; with *Hs*H2A) are shown for clarity. Shaded areas represent the standard error mean of at least three biological replicates. **(F)** Protein-protein interaction (PPI) network of mutant genes isolated in the evolved H2Amacro1 and macroH2A1-HF-sb histone humanized strains (excluding synonymous mutations). Graph was constructed using the STRING database, PPI-enrichment *p* value = 8.41e^-5^. Node colors represent MCL clusters (MCL inflation parameter set to 2). Black colored nodes are interacting genes inferred from the network, the histone genes are colored a lighter color to draw attention to the fact we isolated mutants in human H2A and H2B, not necessarily the yeast histone genes, but the yeast gene names were used to construct the network. Any unconnected nodes and those with fewer than three linked nodes were removed.

To fine-map the inviable residues, we humanized (i) the HFD of macroH2A1 (macroH2A1-HF) and (ii) chimeric fusions of macroH2A1-HF with *Hs*H2A (replacing the N-or C-terminal tails of replicative *Hs*H2A with corresponding regions of macroH2A1-HF; Figure 3A, S7A). We observed that the C-terminal region of macroH2A1-HF (replacing *Hs*H2A C-termini) was sufficient to disrupt humanization (Figure 3A, S7A). In contrast, the N-terminal tail of macroH2A1-HF functionally replaced the N-terminal tail of *Hs*H2A (Figure 3A, S7A). We performed extensive mutagenesis experiments to map the inviable residues of macroH2A1-HF (Methods; Figure S7B–J). We identified a minimal set of 18 residues in the HFD of macroH2A1 that when swapped to the corresponding *Hs*H2A residues led to isolation of *bona fide* macroH2A1 HFD humanized yeast completely lacking replicative *Hs*H2A; we refer to these strains as “macroH2A1-HF-sb” (“sb”: swap-back, color-coded light-blue in Figure 3A–B, S7J). These colonies appeared after ∼3 months of incubation at 30°C on 5-FOA plates, suggesting a massive fitness defect in this “swap-back” mutant. Fitness rapidly improved when clones were patched on YPD plates, a dense mat of cells appeared within two weeks of incubation at 30°C.

### macroH2A1-HF-sb and H2Amacro1 humanized yeasts have reduced fitness

MacroH2A1-HF-sb and H2Amacro1 humanized yeasts formed large cells with a cell cross sectional area on average 4.2 times larger than WT yeast (*Sc* histones; Figure 3C and Figure S8A; cross sectional area of 13 µm^2^ versus 6.3 µm^2^). Remarkably, some cells reached truly enormous sizes ranging all the way up to a cell cross sectional area of 45 µm^2^ (∼50 times larger than WT). Associated with this was a severe increase in doubling time, to over ∼19 hours (Figure 3D, Figure S8B). Relative to WT yeast, the doubling time of macroH2A1-HF-sb and H2Amacro1 humanized yeasts increased 4.3-fold and 5.1-fold, respectively. In comparison, histone humanized yeasts with replicative *Hs*H2A displayed an increased doubling time of 2.4-fold relative to WT yeast. Moreover, macroH2A1-HF-sb and H2Amacro1 humanized yeasts spend a considerably longer time in the lag phase, on average ∼60 hours versus 8.2 hours and 39.5 hours for WT yeasts and histone humanized yeasts with replicative *Hs*H2A, respectively (Figure S8C).

To assess whether growth could improve in macroH2A1-HF-sb and H2Amacro1 humanized yeasts, we continually passaged each in rich medium for up to 60 generations over the course of 4 months. In these evolved strains, we observed only modest improvement to growth (Figure 3E), as their doubling times improved only by ∼0.18-fold and ∼0.33-fold for macroH2A1-HF-sb and H2Amacro1 humanized yeasts, respectively, although the variance in doubling times were reduced (Figure S8B). Moreover, the time spent in lag phase was likewise marginally improved by ∼0.24-fold and ∼0.15-fold for macroH2A1-HF-sb and H2Amacro1 humanized yeasts, respectively. Therefore, continuous culturing led to small, but significant improvements to growth in both macroH2A1-HF-sb and H2Amacro1 humanized yeasts.

We performed whole genome sequencing on ancestral and evolved clones at 30 and 60 generations to identify mutations associated with the fitness increase (Table S5). In the H2Amacro1 histone humanized yeast we observed that all clones lost their mitochondrial genome (retaining a highly amplified mitochondrial origin of replication region), consistent with the overall worse growth of H2Amacro1 humanized yeast compared to macroH2A1-HF-sb strain (which didn’t lose mitochondrial DNA). Surprisingly, we detected a notable mutation in the HFD of the *Hs*H2A domain (R35I). This residue interacts with the DNA phosphate backbone and it is, coincidentally, the orthologous residue of macroH2A1-HF that, when swapped to the replicative *Hs*H2A residue was found to improved humanization (K32R; Figure S7G, see methods). Additionally, we identified a large deletion of the nonessential histone H2B amino-tail (H2BdelG13-K24) in macroH2A1-HF-sb humanized yeast. Intriguingly, deletion of histone H2B amino-tail *in vitro* led to nucleosome destabilization in a thermal stability assay^38^, suggesting, alongside the observed *Hs*H2A-R35I mutation, that one route by which yeast adapt to macroH2A1 HFD or macro domain is though nucleosome destabilizing mutations.

Overall, we identified 52 mutations, of which 42 were nonsynonymous mutations and most of which were not in the histone genes (Table S5). We next constructed an interaction network from these 42 mutations using the String algorithm (Figure 3F). The core of this interaction network (cluster 3; orange) was enriched in chromatin-based biological processes such as Histone lysine demethylation (false discovery rate (FDR) =0.0033) and Chromatin assembly or disassembly (FDR =4.37e^-05^). Additionally, we saw an enrichment for cellular components such as Cytosolic ribosome (cluster 4; yellow, FDR =0.00032) and biological processes such as Endocytosis (cluster 2; red, FDR = 0.00036) and Ubiquitin mediated proteolysis (cluster 5; green, FDR =0.0200). These analyses indicate that both macroH2A1-HF-sb and H2Amacro1 evolved through selection of mutants from a non-random set of genes, which are likely to be adaptive. However, as all clones were isolated in the background of the *DAD1*^E50D^ mutation, a mutant that we have shown to be a potent suppressor of histone humanization^5, 31^, we cannot rule out the possibility of pleiotropy or dependencies of these mutations on *DAD1*^E50D^, therefore we proceeded by studying the ancestral strains which exhibited the fewest mutations (Table S5).

### The histone fold and macro domain of macroH2A1 increase nucleosome repeat length

To assess changes to the structure of chromatin in macroH2A1 humanized yeast we performed Micrococcal Nuclease (MNase) digestions on cross-linked chromatin isolated from strains with yeast histones (*Sc*; WT), histone humanized yeast with replicative *Hs*H2A and the ancestral histone humanized yeast with either macroH2A1-HF-sb or H2Amacro1. We assessed the quality of the digest on an agarose gel and observed that replicative histones, regardless of species, formed correctly phased nucleosomes (Figure S9A, *Sc and Hs*H2A panels) as previously reported^5^. Remarkably, we observed that the nucleosome repeat length (NRL) was slightly increased for humanized yeast with either macroH2A1-HF-sb or H2Amacro1, indicating that both histone fold domain and macro domain of macroH2A1 independently increase the NRL (Figure 4B–C and Figure S9A). To confirm this observation, we sequenced the MNase digested DNA (MNase-seq). Fragment length analysis of the sequenced digested DNA, binned near transcription start sites (TSS), from humanized yeast with either macroH2A1-HF-sb or H2Amacro1 showed a characteristic increase in the NRL genome-wide, when compared to WT yeast or humanized yeast with *Hs*H2A (Figure 4A and Figure S9B,D).

**Figure 4.**
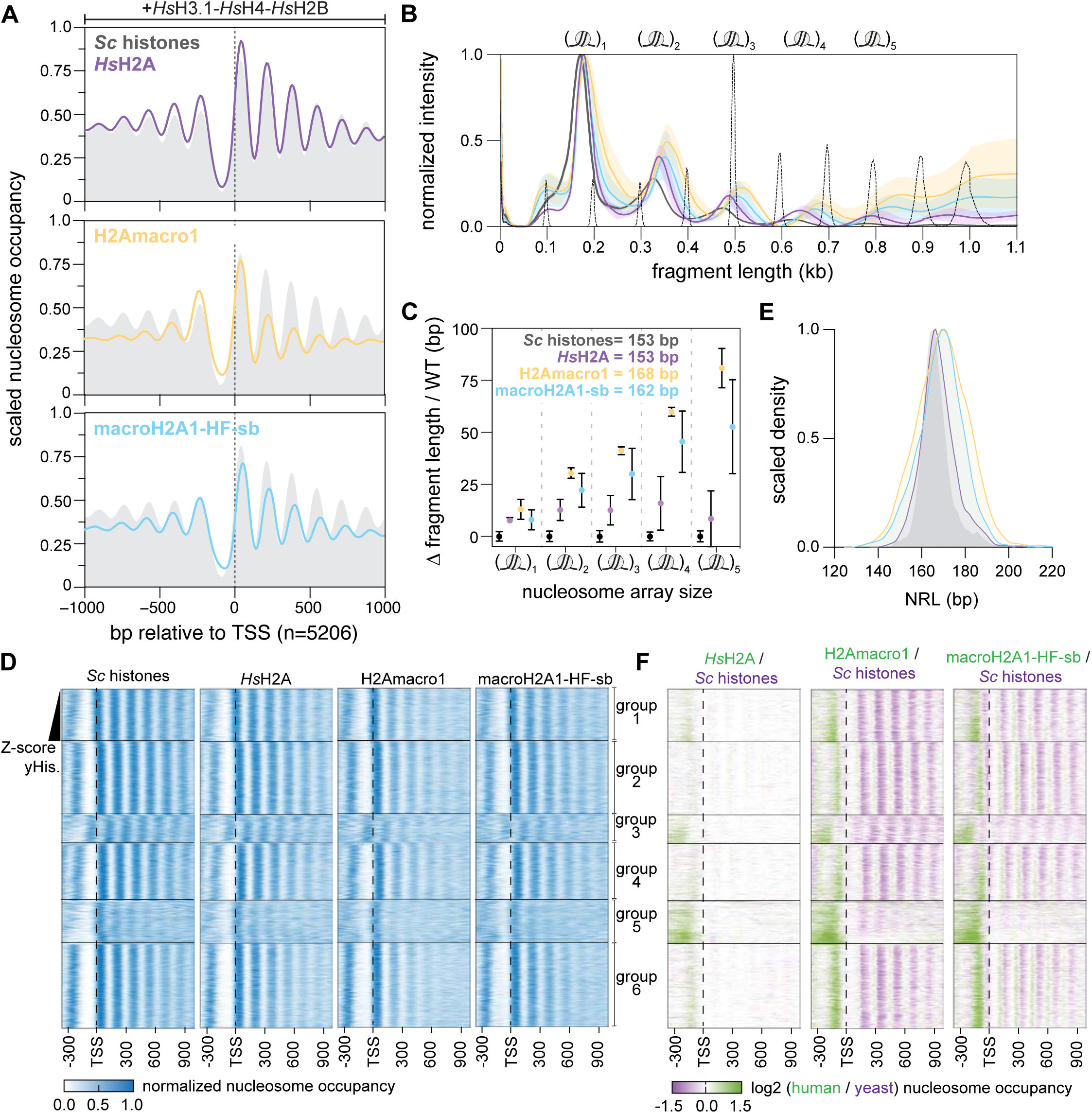
macroH2A1 comprised chromatin has increased nucleosome repeat length. **(A)** Composite plot of nucleosome occupancy relative to the transcription start site (TSS) of 5,206 genes. Gray background occupancy is the mean occupancy of WT yeast and colored lines the mean occupancies of the indicated strain. **(B)** Fragment size analysis of MNase digested DNA using capillary electrophoresis. Mean normalized intensities are shown with the standard deviation of three biological replicates shown as the filled in colored area. Molecular weight maker is shown as the dotted line. **(C)** Difference in fragment length compared to WT digested chromatin. **(D)** Sorted heatmap of nucleosome occupancy near each TSS (n = 5,206 genes). Nucleosome occupancies –350 to +950 bp relative to each gene’s TSS were first sorted through k-means clustering into six groupings. Next, within each grouping, genes were sorted by their z-score ranked transcript abundance in WT yeasts (with increasing abundance). **(E)** Scaled kernel density plot of nucleosome repeat lengths across 4,109 genes relative to the +1 nucleosome. Gray background distribution is observed NRLs of Yeast with *Sc* histones; purple distribution, *Hs* histones; yellow distribution, H2Amacro1; blue distribution, macroH2A1-HF-sb. **(F)** Log2 ratio heatmaps of nucleosome occupancies of heatmaps in panel D.

We next assessed the length of the digested DNA using capillary electrophoresis (Figure 4B). Comparison of fragment lengths to the WT control strain showed that mono-nucleosomes from all histone humanized yeast strains were on average ∼10 bp larger than expected (Figure 4C). For the humanized yeast with replicative *Hs*H2A the 10 bp increase was fixed across all oligonucleosome arrays (mono- to penta-nucleosomes), consistent with the idea that replicative human histones more tightly wrap DNA^39–41^, but do not alter the NRL in yeast^5^. These observations are consistent with and strongly support our direct measurements of nucleosome particle sizes from histone humanized with transmission electron microscopy (Lazar-Stefanita *et al.* co-submitted). We observed that in humanized yeast with either macroH2A1-HF-sb or H2Amacro1 the DNA fragments were larger than expected and displayed the characteristic linear increase with oligonucleosome size, indicative of increased NRL (Figure 4C). From the slope of these increments, we estimated a statistically significant increase to the NRL by 10-14 bp in humanized yeast with either macroH2A1-HF-sb or H2Amacro1 (Figure 4C and S9D). These results suggest that the positioning of individual nucleosomes is shifted downstream of their expected positions relative to the +1 nucleosome.

We next inferred the genome-wide positioning and occupancies of nucleosomes (see methods). We observed, on average, a total of ∼70,000 nucleosomes in our samples (Table S6). Composite-gene analysis of nucleosome occupancies and positions supported the increased NRL relative to the TSS in the humanized yeasts in with either macroH2A1-HF-sb or H2Amacro1 histones, and revealed unexpectedly lower nucleosome occupancies across their genomes (Figure 4A, S9B–F). The positioning of +1 nucleosomes (relative to the TSS) were not significantly altered in humanized yeasts with either macroH2A1-HF-sb or H2Amacro1 nucleosomes (Figure 4A, S9E); whereas, the downstream nucleosomes showed significant dephasing in both strains (Figure 4A, S9E). Interestingly, the humanized yeast with macroH2A1-HF-sb showed uniform nucleosome depletion across the entire gene bodies (Figure S9C); whereas, the terminating nucleosome remained strongly occupied in the H2Amacro1 humanized yeast (Figure S9C; that is the –1-nucleosome relative to the terminating sequence (Ter)). This suggests that the macro domain of macroH2A1 does not interfere with the positioning of the +1 nucleosome (TSS) nor the positioning and occupancy of the terminating nucleosome.

We further examined nucleosome occupancy by k-means clustering the nucleosome occupancy maps of each gene relative to their TSS in WT yeast. Within each cluster we sorted the genes by increasing levels of transcript abundance in WT (see methods). The nucleosome occupancies best clustered into six distinct groups, each of which exhibited unique nucleosome phasing profiles (Figure 4D). Groups three and four showed poor phasing in WT and humanized yeast with *Hs*H2A, and were even less well phased in humanized yeast with either macroH2A1-HF-sb or H2Amacro1 (Figure 4D). From these maps, we estimated the global NRL by measuring the spacing from the +1 to +5 nucleosomes from genes that displayed well phase nucleosomes, observing a net increase to the NRLs in humanized yeast with either macroH2A1-HF-sb or H2Amacro1 (Figure 4D–E; groups 1, 2, 4, and 6). In sum, both H2Amacro1 and macroH2A-sb chromatin showed a significant increase to the NRL, in addition to overall less nucleosome occupancy across gene bodies.

### macroH2A1 chromatin is associated with transcriptional dysfunction

One feature of the nucleosome positioning maps was the accumulation of nucleosomes in the NDR in humanized yeast (Figure 4F). To assess whether nucleosome accumulation was correlated with transcriptional changes of those genes we performed RNA sequencing (see methods, Figure S10A). We observed numerous transcriptional changes to the humanized yeasts with either macroH2A1-HF-sb or H2Amacro1 chromatin. In total we observed 248 genes that were significantly down-regulated and 295 genes that were up-regulated in both macroH2A1-HF-sb and H2Amacro1 humanized yeasts compared to WT yeasts (Figure S10B; Table S7). The down-regulated genes were enriched in KEGG pathways such as Ribosome and Glycolysis, while the up-regulated genes were enriched in biological processes such as Flocculation and Cell adhesion (Figure S10C–D). Additionally, the up-regulated genes showed enrichment to the subtelomeric regions of chromosomes, consistent with loss of telomere silencing (Figure S10E–F).

Accumulation of nucleosomes in the NDR was greater for genes with highly abundant transcript levels in WT yeast. For example, genes in group 6, within the upper 15% of transcript abundance in WT yeast, in both macroH2A1-HF-sb and H2Amacro1 humanized yeast we observed a 93% increase to nucleosome occupancy near the NDR relative to WT (Figure 4F and Figure S9F). In contrast, for the genes in the bottom 15% of transcript abundance in WT, nucleosome occupancy increased only 41% in the NDR relative to WT in both macroH2A1-HF-sb and H2Amacro1 humanized yeast (Figure S9F). The increased occupancy in the NDR was associated with transcriptional repression in both macroH2A1-HF-sb and H2Amacro1 humanized yeasts for the genes in top 15% but not those in the bottom 15% (Figure S11C). Gene set enrichment analysis of the genes in the top 15% group revealed their strong enrichment in processes related to translation, such as small and large ribosomal subunit biogenesis (Figure S11D). We additionally examined group 5, which showed some of the highest levels of accumulation of nucleosomes in the NDR, a 97% increased occupancy, in both macroH2A1-HF-sb and H2Amacro1 humanized yeasts (Figure 4F, S9F). Likewise, the genes in the top 15% of transcript abundance in WT were significantly down-regulated in macroH2A1-HF-sb and H2Amacro1 humanized yeasts (Figure S11E-G), and were enriched with ribosomal proteins and glycolysis related genes (Figure S11H). These results are consistent with our findings that ribosomal RNA levels are reduced in histone humanized yeast (Lazar-Stefanita *et al.* co-submitted) and RNA-sequencing showing the down regulation of genes enriched in ribosomal function in both macroH2A1-HF-sb and H2Amacro1 humanized yeasts (Figure S10C). From these data we suggest that nucleosome accumulation in the NDR is associated with the transcriptional down-turn of highly expressed genes that are critical for central metabolic pathways such as protein translation. Since rRNA levels in histone humanized compared to wild-type yeast are decreased (by ∼2.5-fold, Lazar-Stefanita *et al.* co-submitted), here we suggest that the reduction of ribosomal protein expression is a consequence of overall lower ribosome abundance.

### Transcriptional up-regulation and DNA shape are correlated with nucleosome phasing

We next asked whether the increased NRL we observed was correlated with transcriptional changes. We explored the nucleosome occupancy and positioning data of 114 up-regulated and 154 down-regulated genes shared between humanized yeasts (macroH2A1-HF-sb and H2Amacro1, versus WT yeast; Figure 5A, S10A). We considered the change in position for five nucleosomes downstream the TSS (Figure 5B–C). As expected, we observed no difference in positioning of these nucleosomes in humanized yeast with only replicative histones (Figure 5C, *p* = 0.57). However, for strains with either macroH2A1-HF-sb or H2Amacro1 chromatin we observed a consistent average shift to the right of +10 bp for nucleosome in both up- and down-regulated genes (Figure 5C, *p* < 1e^-4^). When examining the change in nucleosome positioning between differentially expressed genes in yeast with either macroH2A1-HF-sb or H2Amacro1 chromatin, we observed that nucleosomes from the up-regulated genes were significantly less shifted downstream of the TSS (Figure 5C, S12). As for up-regulated genes we observed no linear increase to the change in nucleosome position from the +1 to +5 nucleosomes (Figure S12), suggesting that highly expressed genes retain better nucleosome positioning.

**Figure 5.**
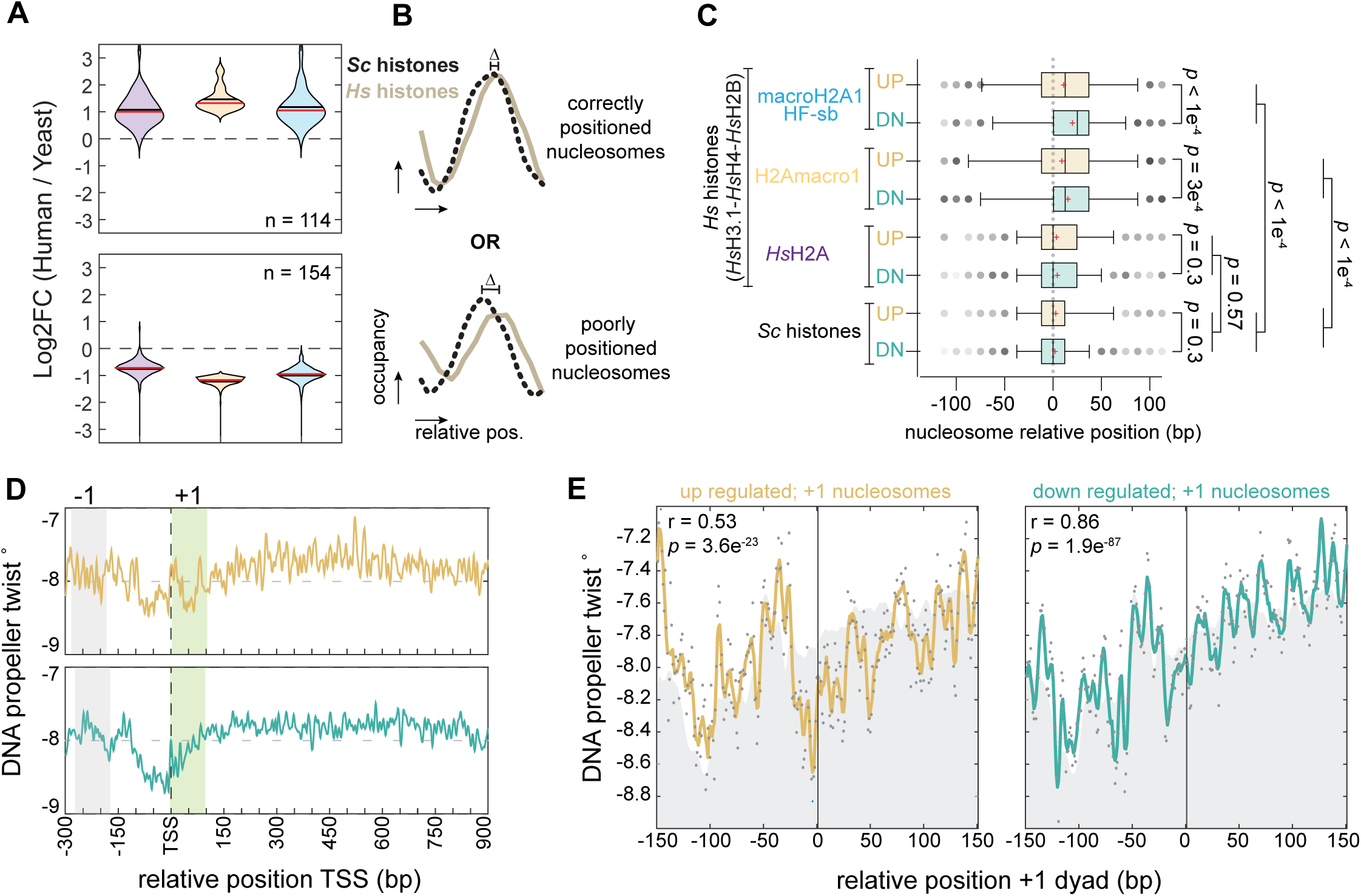
Up-regulated genes display better nucleosome positioning and exhibit distinct predicted DNA shape. **(A)** Log2 fold change expression genes shared between macroH2A1 humanized yeasts that are up- or down-regulated in comparison to WT yeast. **(B)** Examples of nucleosome comparisons that are well positioned between WT and humanized yeasts and those that are poorly position in humanized yeasts. Briefly, we compared the relative position of five nucleosomes downstream of the TSS in each humanized lineage versus the composite position of WT nucleosomes. Each comparison was made to each specific cluster of nucleosomes (see Figure S8F). **(C)** Comparison of the relative positioning of nucleosomes of up- and down-regulated genes in WT and humanized yeasts. Comparisons of the mean change in relative nucleosome positioning between up- and down-regulated genes, for macroH2A1-HF-sb, -7 bp, *p* < 1e^-4^; and for H2Amacro1, -5 bp, *p* = 3e^-4^ (FDR corrected ordinary one-way ANOVA tests). **(D)** Composite plot of the DNA shape feature propeller twist near the TSS of either up- or down-regulated genes. The –1 and +1 nucleosome positions are indicated in shaded gray and shaded green regions, respectively. **(E)** Composite plot of the DNA shape features centered on the dyad of the +1 nucleosomes of either up- or down-regulated genes. Gray back ground in the average composite plot of the +1 nucleosomes for 4,109 genes.

DNA shape features, such as propeller twist (the angle between the plane of the two bases), impart important information shaping the organization of nucleosomes^25^. We therefore examined whether we could detect unique signatures of DNA shape at the dysregulated genes. To examine the DNA shape, we determined the propeller twist near TSSs (-300 bp to +900 bp; Figure 5D)^25^. Examination of composite DNA shape plots in up- and down-regulated genes revealed striking differences in the DNA shape near the +1 nucleosomes (Figure 5D). Up-regulated genes showed a near-symmetrical “U” shape pattern, whereas down-regulated genes showed an asymmetrical shape relative to the +1-nucleosome dyad (Figure 5E), the latter being more similar to the DNA shape of the complete composite set of genes (Pearson r = 0.5169 and r = 0.8598, respectively). As we showed above up–regulated genes were enriched at subtelomeric regions (Figure S10E–F). We therefore examined the DNA shape of the subtelomeric genes (< 30 kb from telomere) in comparison to non-subtelomeric (> 30 kb from telomere) and revealed distinct DNA shapes relative to the TSS and dyad of the +1 nucleosome (Figure S13A–C). Taken together, our data show that transcriptionally up–regulated genes with better phased nucleosomes in both macroH2A1-HF-sb and H2Amacro1 humanized yeasts exhibit distinct chromosomal locations and unique DNA shape near their +1-nucleosome.

### macroH2A1 histone fold and macro domains alter 3D genome organization and drive genome instability

We were curious what effect chromatinization with either macroH2A1-HF-sb or H2Amacro1 have on genome structure and stability. We first explore the consequences on chromatin folding by performing Hi-C. In agreement with the companion paper (Lazar-Stefanita *et al.* co-submitted), in histone humanized yeast, we observed a loss of inter-pericentromeric contacts (Figure 6A–B, S14B). These changes were all observed to be similar, or greater, in both macroH2A1-HF-sb and H2Amacro1 humanized yeasts compared to humanized yeast (Figure 6B–C, S14A–B). The typical “cruciform” arrangement of intra-chromosomal contacts near the pericentromere was largely lost in both macroH2A1-HF-sb and H2Amacro1 humanized yeasts (Figure 6A), noticeable by the increase to interactions between the chromosomal arms with the pericentromeric regions (See ratio maps, observing the two dark red axis emanating ±45° perpendicular of the centromeric center; Figure 6B).Quantification of inter-pericentromeric contacts showed that two clones of humanized macroH2A1-HF-sb had dramatically decreased inter-pericentromeric contacts compared to WT and to histone humanized yeast (Figure S14B). These results indicate that the structure of the pericentromeres is affected, leading to strong centromere de-clustering in histone humanized yeasts. Consistent with this, we observed a significant increase in centromeric RNA in all humanized strains, with the highest expression of *CEN* RNAs in those with either macroH2A1-HF-sb or H2Amacro1 histones (Figure S14C). Both reduced inter-pericentromeric clustering and elevated levels of *CEN* transcription suggest defects in chromosome segregation. Indeed, we observed high rates of chromosome instability (CIN) in both macroH2A1-HF-sb and H2Amacro1 humanized yeasts (Figure 6D, S15C–G). All humanized yeast were generated in the mutant *DAD1*^E50D^ background, which we have previously shown to purge aneuploidies in histone humanized yeast^31^. Remarkably, all macroH2A1-HF-sb and H2Amacro1 humanized yeasts had at least one or more aneuploid chromosomes despite the *DAD1*^E50D^ mutation (Figure 6D, S15C–G Table S8), suggesting that both the histone fold and macro domain of macroH2A1 interfere with its adaptive benefit. Overall, the increased rate of CIN is consistent with the negative GIs we observed between macroH2A1 overexpression and kinetochore genes, suggesting macroH2A1 interferes with centromeric chromatin in yeasts (Figure S3B).

**Figure 6.**
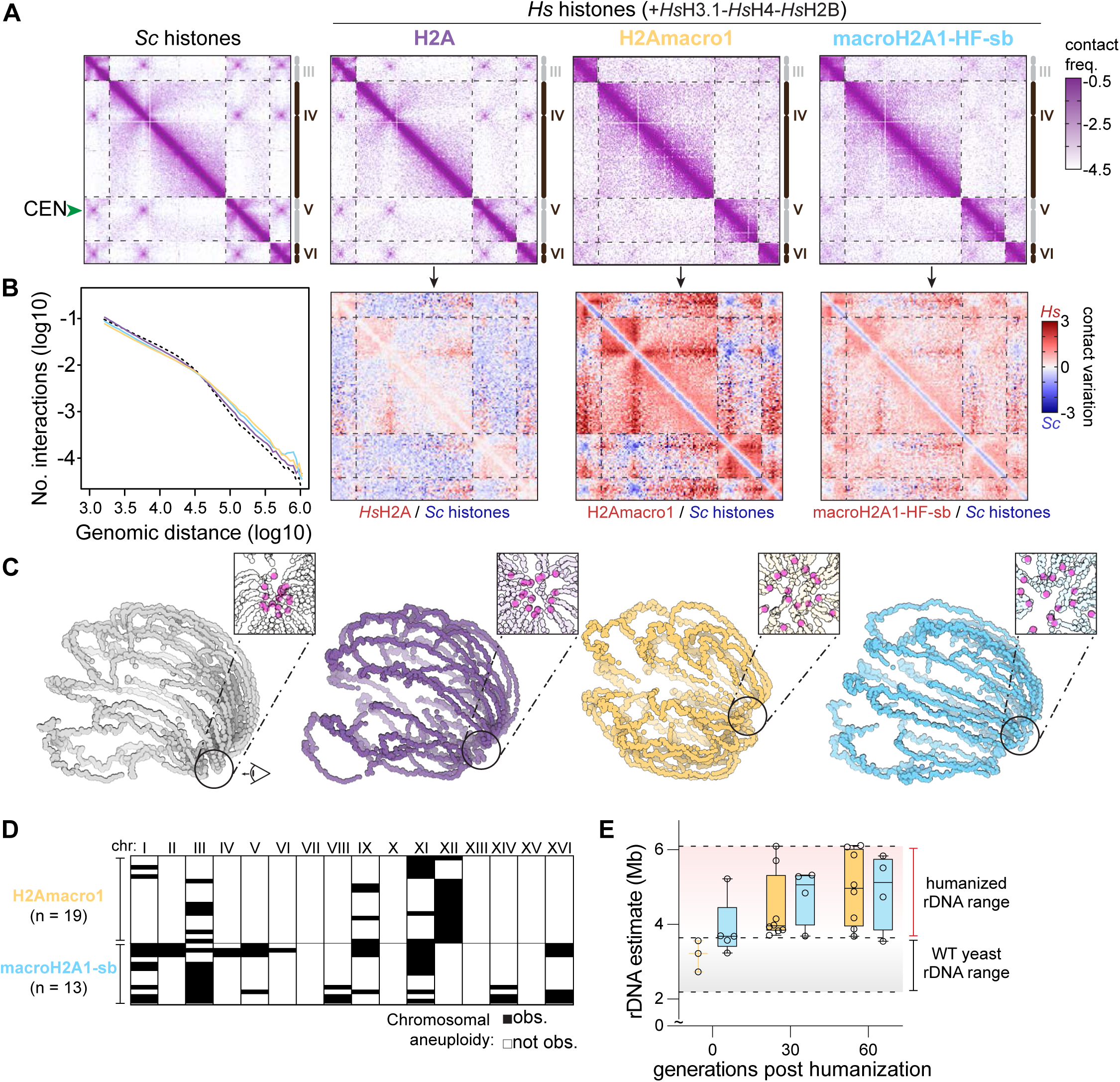
Decreased short-range chromatin interactions and chromosome instability in macroH2A1 humanized yeast. **(A)** Subset of Hi-C heatmaps showing chromosomes *III* to *VI*. An example inter-centromeric contact is indicated with a green arrow. Normalized contact frequencies were binned at 5 kb resolution. Purple to white color scale indicates increase in contact frequency (log10). **(B)** Left: contact probability decay as a function of the genomic distance plot represents the average decay of intra-chromosomal contact frequency with the increment in their genomic distances. Right: log2-ratio maps of human to yeast contact maps in panel A. **(C)** 3D average representations of the complete Hi-C maps in panel A. **(D)** Observed chromosomal aneuploidies in macroH2A1 histone humanized yeast. Aneuploidies were inferred from chromosome sequencing coverage (green, observed; white, not observed). Number of isolates examined is showed where each row represents one isolate. Chromosome coverage plots are displayed in Figure S15. **(E)** Estimation plot of rDNA array size in macroH2A1 humanized yeasts after humanization and after growth for 30 and 60 generations in rich medium. The normal ranges observed for WT and humanized yeasts are provided as colored ranged (Lazar-Stefanita *et al.* co-submitted).

Furthermore, the Hi-C maps revealed that chromatinization with either macroH2A1-HF-sb or H2Amacro1 promoted an overall decompaction of the yeast chromatin, as indicated by the decrease of short-range contacts (<20 Kb; Figure S14A). Loss of short-range contacts may be a consequence of the reduced nucleosome occupancy in both macroH2A1-HF-sb and H2Amacro1 humanized yeasts (Figure 4A). Moreover, the increase to NRL may facilitate chromatin fiber flexibility potentially leading to further decompaction^42, 43^. Correspondingly, the loss of short-range interactions was accompanied by an increase in long-range contacts (>20 Kb) in both macroH2A1-HF-sb and H2Amacro1 humanized yeasts (Figure 6B ratio maps, S14A). Increased distal interactions are partly attributable to the loss of strong inter-pericentromeric interactions, which typically constrain chromosomes, thus substantial declustering of the pericentromeres may promote increased intrachromosomal contacts as the chromatin fiber is overall less constrained in spatially. Moreover, the increased distal interactions may be a proportional response to chromatin decompaction at the shorth length scale (i.e., if the total number of short-range contacts decrease, long-range contacts increase proportionally to the sum total of contacts). We propose that the combination of decreased nucleosome occupancy and increased nucleosome linker length drive an overall decompaction of chromatin. In addition, our data are consistent with longer nucleosome linkers creating more open chromatin structure^42, 43^.

In Lazar-Stefanita *et al.* replicative human histones were shown to cause loss of rDNA silencing and consequently, rDNA array instability leads to the rapid expansion of the array to over 5 Mb in size (>1/3 of the yeast genome) (Lazar-Stefanita *et al.* co-submitted). Here we estimated the size of the rDNA array for strains with macroH2A chromatin and observed a similar increase in the rDNA size (Figure 6E). We observed a similar trend of rapid expansion following the initial humanization event, suggesting that the mechanism is similar among all histone humanized yeast strains. We propose that rDNA expansion is likely quenched by the upper chromosomal arm length limit of ∼7 Mb, which equates to a chromosomal arm nearly equal in length to half the distance of the spindle-pole axis^44^.

### macroH2A1 histone fold and macro domains promote ectopic chromosomal rearrangements

Examination of our whole genome sequencing data of the ancestral and evolved clones revealed how genomes evolved in the presence of macroH2A1-HF-sb and H2Amacro1 chromatin. In all clones we observed the appearance of multiple chromosomal rearrangements (Figure 7 and S15). These were less frequent in the H2Amacro1 humanized yeasts (Figure S15) i.e., in this humanized yeast we observed aneuploid chromosome *XII* in which one copy displaying an internal deletion of ∼160 kb, with the breakpoints mapping near two Ty1 long terminal repeats (LTRs; Figure S14D, S15A–B). Intriguingly, the aneuploidy and the deletion were stable across the 60 generations that we tracked, as such we were curious how the size of the rDNA array on either copy of chromosome *XII* compared (Figure S15D–E). Contact quantifications of chr *XII* Hi-C map suggested that intra-chromosomal contacts across the rDNA array were increased (contacts between the right arm of chromosome *XII* across the rDNA locus in histone humanized cells; Figure S14D). The observed increase could be trivial, owing to fact that the strain has two copies of chromosome *XII*, or potentially due to one rDNA array being a smaller barrier^45^. To tease out either scenario we noticed that the increased intra-chromosomal contacts across the rDNA array were restricted in the flanking regions of the deletion, suggesting that the increased contacts arise solely from the chromosome with the internal deletion (Figure S14D–E). Moreover, these intra-chromosomal contacts were more frequent than expected if only due to the copy number increase of having two copies of chromosome *XII* (2.7x increase vs. 1.3 x increase, respectively), suggesting that the rDNA array on the chromosome with the ∼160 kb deletion is potentially reduced in size^46^.

**Figure 7.**
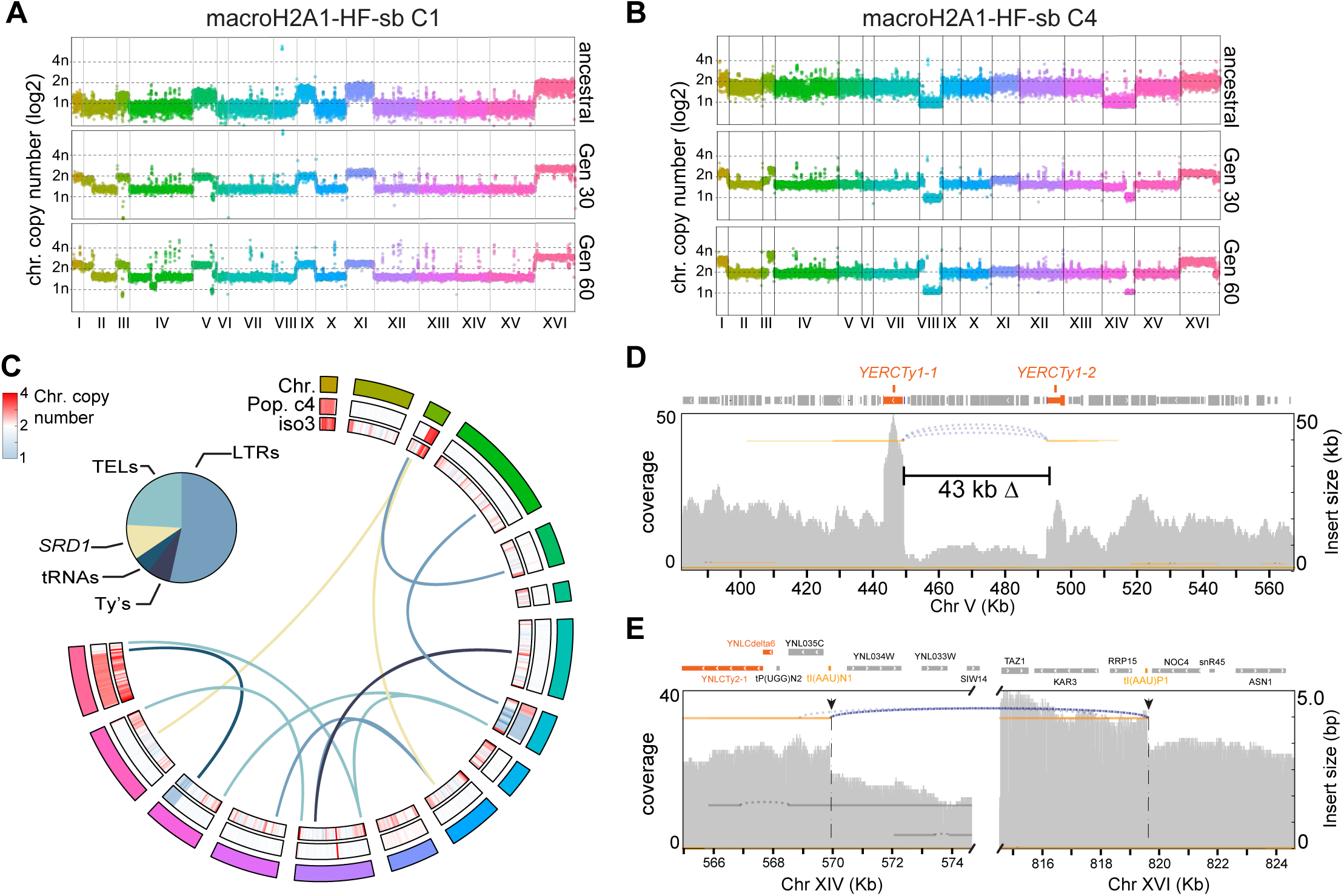
The histone fold of macroH2A1 promotes ectopic recombination events between repetitive elements. **(A–B)** Chromosome coverage plots from whole genome sequencing of macroH2A1-HF-sb clones 1 and 4. Three times points are shown; ancestral, after 30 generations and after 60 generations in rich medium. Ploidy estimates were normalized to the median coverage of the lowest covered chromosome (e.g., for clone 1, this is a ∼50 Kb region on chromosome V showing a deletion from ∼440 Kb to 492 Kb, containing the essential genes SCC4, SPT15, COG3, thus is likely to be present at least at a single copy). Ploidy is drawn on a log2 scale. **(C)** Circos plot of chromosomal rearrangements inferred from Nanopore sequencing of macroH2A1-HF-sb humanized yeast clone 1. Chromosomes are presented in clockwise fashion from Chr. I to Chr. XVI. For each chromosome the sequencing coverage (log2 normalized to the median, binned at 24 Kb) is plotted both from the Illumina data (Pop. C4 track) and nanopore data (iso3 track). Translocations are plotted as connecting links between chromosomes and are colored by the type of sequences which recombined. Inset pie chart depicts the relative proportions of each class of repetitive elements for which we observed translocation event between (both clone 1 and clone 4; Table S9). **(D)** Evidence of a 43 kb deletion in chromosome V between two Ty1 elements, YERCTy1-1 and YERCTy1-2 (Orange boxes). Mapping reads are plotted in orange and gaps in the reads are plotted as dashed arches. **(E)** Evidence of a translocation between chromosome XIV and XVI between two isoleucine tRNAs. Gene track is shown above with tRNAs colored in yellow–orange and Ty/LTRs element in dark-orange. Below is a samplot of the nanopore sequencing data, both the coverage is plotted and the insert size of reads (those reads which mapped non-contiguously). Regions on chromosome XIV and XVI with noncontiguous mapping nanopore reads are shown. Mapping reads are plotted in orange and gaps in the reads are plotted as dashed arches.

Examination of whole genome sequencing coverage plots from clones 1 and 4 of the macroH2A1-HF-sb humanized yeast revealed numerous chromosome breakpoints, as indicate by abrupt changes in coverage (Figure 7A–B). Moreover, both clones were clearly polyploid, with the majority of chromosomes at a copy number of two (normalized to regions of deletions, which contain essential genes). Many break points mapped to repetitive elements such as Ty elements, Ty LTRs, tRNAs, and Sub-telomeres. Therefore, we were unable to conclusively map these putative chromosomal rearrangements using short paired end Illumina sequencing data. Moreover, as we performed our Hi-C experiments in the ancestral strains that did not exhibit some of these putative chromosomal rearrangements (Figure 7A–B) we could not leverage the contact maps to map them.

We generated nanopore reads from three isolates from both clones 1 and 4 of macroH2A1-HF-sb humanized yeasts. As suggested by the Illumina sequencing, we observed numerous translocations between Ty’s, LTRs, tRNAs, and sub-telomeric regions (Figure 7C; Table S9). For example, we observed a large ∼43 Kb internal deletion between the Ty1 elements, YERCTy1-1 and YERCTy1-2, on chromosome *V* (Figure 7D). Additionally, we observed a well-supported translocation between chromosome *XIV* and *XVI*, which we mapped to a translocation event between two isoleucine tRNAs (Figure 7E). In conclusion, long read sequencing aided in revealing the complex nature of chromosomal structural variants in macroH2A1-HF-sb humanized yeast.

## Discussion

The complete exchange of replicative histones for variant histones in yeast led to significant consequences, particularly for the H2A variant, macroH2A1. Moreover, mutational swapback analysis of the macroH2A1 HFD and spontaneously isolated mutations (H2A-R35I and H2BdelG13-K24), together suggest that the primary toxicity of macroH2A1 in yeast is the over-stability of macroH2A1 containing nucleosomes. We hypothesize this may lead to severe and pleiotropic phenotypic consequences, such as nucleosome dephasing. Globally we observed a downstream shift of nucleosomes across the genome relative to the TSS. Intriguingly, the exceptions were up-regulated genes in macroH2A1 humanized yeast, which showed less nucleosome dephasing – suggesting that transcription-coupled nucleosome remodeling may improve nucleosome positioning. These data support the model suggesting that non-replicative histone variants, or at least macroH2A, can alter the basic organization of nucleosomes *in vivo*.

We also observed a significant accumulation of nucleosomes in the NDR in histone humanized clones, an effect which was more pronounced in macroH2A1 humanized yeast. Whether or not tied to transcription-coupled nucleosome turnover, accumulation in the NDR was generally positively correlated to the abundance of a gene’s transcript in WT yeast. These data suggest that levels of transcription may inform the deposition of macroH2A1. Reduced transcription and increased accumulation of macroH2A in the NDRs was most apparent when examining highly expressed genes (in WT yeasts) such as ribosomal protein or glycolytic genes. Thus, the accumulation of macroH2A in the NDRs of transcriptionally down-regulated genes may be due to reduced nucleosome eviction by chromatin remodelers such as the RSC complex^47–49^. Intriguingly, in our companion paper we observed a global down-turn of the total levels of ribosomal RNA (∼2.5-fold) likely due to aberrant rDNA array regulation (Lazar-Stefanita *et al.* co-submitted). Thus, we propose a feed-back mechanism, whereby reduced levels of rDNA cascades to a reduction in transcription of ribosomal proteins, thereby leading to accumulation of macroH2A1 nucleosomes by way of reduced nucleosome eviction near the NDR of ribosomal protein genes. We cannot directly rule out the possibility that macroH2A1 accumulation at these genes drives reduced gene transcription, however these models are not mutually exclusive. Put together, we observed two phenomena related to transcription-coupled nucleosome occupancy. First, nucleosome arrays were better phased with increased transcription, and second, nucleosome accumulation in the NDR was inversely correlated with decreased transcription. Reduced nucleosome eviction may explain the latter, whereas transcription-coupled nucleosome remodeling may explain the former^50^.

Biochemical reconstitutions have established that ATP-dependent chromatin remodelers set the phasing of nucleosomes^22, 24–27^. Our observations that replicative histones, regardless of species, result in normal phasing of nucleosomes in yeast, support the idea that replicative nucleosomes’ interactions with chromatin remodelers are deeply conserved^51^. In line with this, *in vitro* reconstitutions have also shown that purified yeast chromatin remodelers properly phase replicative histones, regardless of the species’ histones examined^25^. However, as our data suggest, certain histone variants may lack (or have new) interactions essential to maintaining correct phasing in yeast. We propose that histone variants may modulate locally distinct nucleosome organization through exclusionary interactions with chromatin remodelers. We do not address this hypothesis directly using our *in vivo* system, as we do not precisely modulate the levels of specific chromatin remodelers. However, biochemical work has shown that macroH2A1 nucleosomes display reduced recruitment of chromatin remodelers^18^ and the efficient deposition of macroH2A1 in mammals requires the ATPase–dependent action of LSH/HELLS, a SNF2-like chromatin remodeler^52^. While yeast does encode a homolog of mammalian LSH, Irc5, it likely lacks the specific protein-protein interactions required to interact with macroH2A1^53^. Future efforts should address the effects of histone type in combination with chromatin remodelers on basic nucleosome organization.

The total absence of replicative *Hs*H2A histone resulted in dramatic genome instability. First, we observed an almost complete loss of inter-pericentric interactions, a signature of Rabl chromosome organization^54^, consistent with a severe defect to chromosome segregation, manifested as increased rates of aneuploidy (Figure 6D). In agreement, we observed numerous negative GIs between macroH2A1 and yeast deletion of genes encoding kinetochore proteins – suggesting that macroH2A1 may further disrupt centromeric chromatin. All humanized yeasts were generated in the *DAD1*^E50D^ mutant background, which we have shown to rescue kinetochore dysfunction and reduce chromosomal aneuploid levels^31^. Surprisingly, humanized yeast with macroH2A1 displayed numerous aneuploids, suggesting the adaptive benefit of *DAD1*^E50D^ is reduced consistent with macroH2A1 directly interfering with kinetochore function.

The phasing and occupancy of nucleosomes is thought to be critical for genome integrity^58^. We observed increased rates of genome instability brought on by chromatinization with the two macroH2A1 derivatives studied here. For clones with H2Amacro1, we observed only one large deletion event that was stable over many generations. However, for the macroH2A1-HF-sb clones we observed continuing accumulation of deletion and rearrangements, suggesting that the histone fold domain of macroH2A1 contributes the most to genome instability. There is substantial evidence of macroH2A1’s role in maintenance of genome *stability* in metazoans^52, 55^. MacroH2A1 histone promotes the resolution of DNA double strand breaks through homologous recombination (HR), through the formation of protective domains of chromatin^56, 57^. Our data clearly shows that both the histone fold and macro domains of macroH2A1 alone are insufficient to ensure genome stability, and when comprising the entirety of the chromatin, each can drive genome instability. Interestingly, certain clones of the histone humanized yeasts with macroH2A1-HF-sb, which lacked chromosomal rearrangements, carried mutations in genes involved in HR-directed repair of DNA damage (*rad54*-S121R; Table S5), perhaps tempering the effects of macroH2A1 histone fold domain. Breakpoints of the chromosomal rearrangements in macroH2A1 humanized yeasts mapped to Ty elements, LTRs, sub-telomeres, and tRNAs (Table S9), suggesting that these repetitive regions of the genome become fragile when chromatinized with macroH2A1-HF-sb. Moreover, the chromatin decompaction we observed in histone humanized yeast with either macroH2A1-HF-sb or H2Amacro1, may facilitate increased interactions between distantly located repetitive elements. Lastly, non-conserved protein-protein interactions between replicative H2A and macroH2A1 histone fold domain may further drive genome instability, particularly ectopic recombination events.

The *in vivo* manipulations of human variant histones in yeast set the stage for reconstitution of more complex complements of histones. We generated strains that lack entirely replicative H3, H2A, and H2B, replaced by the non-replicative human variant histones H3T, H3.4, TsH2B. This result was surprising, given that yeast has no corresponding ortholog of these variant histones and that H3.4 and TsH2B primarily package DNA during spermatogenesis^59^. Whether or not the underlying chromatin structure is perturbed in these strains remains to be investigated. Further effort in this system should be coupled to manipulations of chromatin remodelers and precise transcriptional changes to determine the factors which regulate chromatin structure and function *in vivo*.

## Author Contributions

M.A.B.H., D.M.T., J.D.B. designed the research; M.A.B.H., L.L.S., G.O., A.W., D.M.T., M.J.S. carried out experimental work; M.A.B.H., G.O., L.L.S. performed data analysis; M.A.B.H. wrote the original manuscript draft and prepared figures; M.A.B.H, L.L.S, D.M.T, J.D.B. edited the manuscript.

## Methods

### RESOURCE AVAILABILITY

#### Lead contact

Further information and requests for resources and reagents should be directed to and will be fulfilled by the lead contact, Jef D. Boeke.

#### Materials availability

All yeast strains and plasmids generated in this study are available from the lead contact upon request. This study did not generate new code.

#### Data and code availability

All sequencing data generated in this study (whole genome sequencing, HiC, RNA sequencing, and MNase sequencing) have been deposited to the sequence read archive under the BioProject PRJNA950985.

### EXPERIMENTAL MODEL AND SUBJECT DETAILS

#### Strains, plasmids, and oligos used

All strains and plasmids used in this study are listed in Tables S1 and Table S2, respectively, and are available upon request. Sequences of oligonucleotides used are provided in Table S3.

### METHOD DETAILS

#### Histone Humanization Assay

Histone humanizations were performed in the *DAD1*^E50D^ dual-histone plasmid shuffle strain (yMAH700), unless otherwise indicated. The *DAD1*^E50D^ mutation improves humanization rates by a factor ∼10^4^ by weakening kinetochore-microtubule interactions^31^. The shuffle strain, where a single set of yeast core histone genes is maintained on a counter-selectable plasmid (*URA3*; Superloser plasmid, pDT139), is transformed with the appropriate human histone plasmid (containing the *TRP1* marker). This “Superloser” plasmid can be destabilized following addition of galactose, using a *GAL10* promoter adjacent to the *CEN* sequence, and then swapped for an orthogonal plasmid containing a full complement of human histones by using the 5-FOA negative selection^30^. This forces yeast to subsist solely on the incoming human histone plasmid. Once transformants were visible, three clones were inoculated into 5 mL of SC–TRP+GAL/RAF liquid medium and grown until saturation (typically 2 days). Culture absorbance (A_600_) was measured and then 1µL, 10µL, 100µL, and 1mL of the saturated culture was plated to SC– TRP+5FOA agar plates. Agar plates were then incubated at 30°C for up to three months, within a sealed container with a damp paper towels to maintain moisture. Only colonies appearing after 2 weeks of incubation were counted and PCR genotyped to verify loss of yeast histones as previously described^30, 31^. Humanization frequencies were then determined by taking the ratio of colony forming units divided by the total number of cells plated. In some cases, where indicated, the humanization frequencies were normalized to the value of humanization for replicative human histones.

#### Protein extraction and western blotting

Immunoblotting of macroH2A-GFP (plasmid pMAH276) and human H2A-GFP (plasmid pMAH282) was performed in the wild-type shuffle strain (yDT67). Briefly, strains were first transformed with a *URA3* plasmid encoding four human histones (with either macroH2A-GFP or human H2A-GFP, in addition to human H3.1, H4, and H2B). Transformants were then grown at 30°C overnight in SC–Ura medium and the following morning diluted in fresh medium and grown until mid-log phase (A_600_ ∼ 0.8 – 1.0). Cultures were then collected with centrifugation, washed once with water, and resuspended in lysis buffer (40mM HEPES-NaOH, pH 7.5, 350 mM NaCl, 0.1% Tween 20, 10% glycerol) + protease inhibitors (cOmplete)). Resuspensions were transferred to tubes with a pre-aliquoted amount of 0.5 mm diameter yttria-stabilized zirconium oxide beads and cells disrupted at 4°C using the MP-Bio FastPrep-24™ lysis system. Lysate was centrifuged at maximum speed for 25 minutes, and clarified lysate was used for western blotting.

Approximately ∼10 µg of protein was loaded on a 12% Bis-Tris NuPAGE® gel in MES buffer. Protein was then transferred to 45 µm LF PVDF membranes using the Bio-Rad Trans-Blot Turbo system, following the manufactures specification and using the mixed molecular weight preset. Transferred membranes were then blocked for 1 hour at room temperature with a 1:1 solution of TBS buffer and LiCor blocking buffer. Next, membranes were incubated overnight at 4°C with primary antibodies in a 1:1 solution of TBST (TBS + 0.05% Tween20) and LiCor blocking buffer (Rabbit anti-GFP, Torrey Pines Scientific TP401; and Mouse anti-alpha-tubulin, Sigma T5168). Membranes were then washed 5x times with TBST, with incubations of 10 minutes between washes at room temperature. Then membranes were incubated with fluorescent secondary antibodies (IRDye® 800CW Goat anti-Mouse IgG and IRDye® 680RD Goat anti-Rabbit IgG) in a 1:1 solution of TBST and LiCor blocking buffer with 0.01% SDS for 1.5 hours at room temperature. Finally, membranes were washed 5x times with TBST, with incubations of 10 minutes between washes at room temperature, and imaged using an Odyssey® imaging system.

#### Histone fluorescence protein tag and imaging

Fluorescence imaging of macroH2A-GFP and human H2A-GFP was performed in the wild-type shuffle strain with a nuclear envelope RFP tag (Nup49-RFP; strain yMAH1279). Briefly, strains were first transformed with a *URA3* plasmid encoding four human histones (with either macroH2A-GFP or human H2A-GFP, in addition to human H3.1, H4, and H2B). Transformants were then grown at 30°C overnight in SC–Ura medium and the following morning diluted in fresh medium and grown until mid-log phase (A_600_ ∼ 0.6 – 0.8). Cells were then adhered to the surface of an ibidi µ-slide VI with Concanavalin A from *Canavalia ensiformis* (10 mg/mL in water) and imaged using an EVOS M7000.

#### macroH2A1 overexpression and growth assay

macroH2A1 was cloned into a galactose inducible CEN/ARS plasmid (pMAH692) and transformed into BY4741. Transformants were grown at 30°C overnight in SC–Leu and the following morning normalized to A_600_ ∼1.0 and dotted out onto either SC–Leu or SC–Leu+Gal agar plates. Plates were incubated at 30°C for two days and then imaged.

#### High-throughput genetic interactions screen

The genetic interactions screen was performed as previously described^60, 61^. We used a conditional overexpression plasmid containing a *LEU2* selectable marker and macroH2A1 driven by the *GAL1* promoter (pMAH692). Using high-throughput, mating-based method, selective ploidy ablation (SPA)^62, 63^ we transferred the plasmid, as well as an empty control plasmid, into an array of the yeast deletion collection of non-essential genes; about 4800 strains in total^64^. The assay was performed using a semi-automatic robotic pinning system, the ROTOR HDA (Singer Instruments, UK) and rectangular agar plates containing the deletion collection previously arrayed as 384 different strains in quadruplicate per plate, i.e., at 1536 colony density. Each incubation step was performed at 30°C. The final SC–Leu 2% galactose 5-FOA agar plates of the assay were incubated for 4 days and imaged using a Scan Maker 9800XL Plus (Mikrotek) plate scanner. The colonies were analyzed using colony quantification

software^65, 66^. Colonies that grew poorly with the empty control plasmid were excluded from the analysis.

#### Histone humanized yeast plate reader growth assays

The histone humanized yeast, yDT180 (derived from the *DAD1*^E50D^ shuffle strain), was transformed with *URA3* CEN/ARS plasmids encoding a full complement of human histones (either all replicative histones (pMAH22) or a single variant with 3 replicative histones (e.g. human macroH2A1, *Hs*H2B, *Hs*H3.1, and *Hs*H4; pMAH87)) or encoding just *Hs*H2B, *Hs*H3.1, and *Hs*H4 (pMAH27). Transformations of histone humanized yeast were modified as follows. A single colony to be transformed was grown until reaching saturation in YPD. The night before transforming, this culture was diluted 3:200 in fresh YPD and grown at 30°C for at least 12 hours or until A_600_ ∼0.6 was reached. From here standard lithium acetate transformation procedures were followed. To ensure isolation of transformants, we transformed at least 1 µg of plasmid DNA. Plates were left to incubate at 30°C for up to two weeks until transformants appeared.

Transformants were then cultured for 5 days in 5 mL of the appropriate liquid medium to maintain selection for both plasmids (SC–Trp–Ura). Once cultures reached saturation, they were diluted to A_600_ ∼1.0 and this suspension was used to inoculate 220 µL of growth medium to a starting A_600_ of 0.1 in a 96-well flat-bottomed UV transparent plate. Growth was then monitored at 30°C for 120 hours, with measurements of the A_600_ every 15 minutes, using EON microplate Spectrophotometer (Biotek). Growth curves were analyzed using manufacture’s supplied software and plotted in Prism. Doubling times were calculated as the ratio of the natural log to the rate of growth during log phase (*ln*2/*r*), where growth rate (*r*) is equal to the natural log of the change in A_600_ over a given time interval, 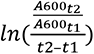.

#### Construction of expanded set of histones expressing plasmids

In order to approach this experiment, we needed an expanded set of orthologous histone promoters available for expressing core histones in *S. cerevisiae* (minimally, we needed six total promoters). This is to reduce sequence similarity between the two plasmids, thereby limiting plasmid recombination events^30^. To this end, we cloned the histone genes and promoters of the closely related species *S. eubayanus* into a counter-selectable *URA3* plasmid (Figure S5A–B). To ensure the histone loci of *S. eubayanus* function in *S. cerevisiae* we first PCR amplified and cloned each pair (SeHTA1B1HHF2T2; pMAH303 and *SeHTA2B2HHF1T1*; pMAH296) into a BssHII linearized *TRP1* CEN/ARS plasmid (pRS414) by yeast gap repair (Figure S4A–B). Plasmids were recovered from yeast, transformed in to bacteria, and verified by digestion. The viability of *S. eubayanus* histone genes and promoters were tested using our dual-plasmid histone shuffle assay (Figure S4C–E). Lastly, the histone clusters *HTA2B2* and *HHF1T1* were subcloned into superloser plasmid (pMAH316) to construct the *S. eubayanus* based histone shuffle strain.

#### *SWR1* CRISPR/Cas9 deletions

We deleted the coding sequence of *SWR1* from the histone shuffle strain using CRISPR/Cas9 genome editing as previously described^31^. A targeting guide RNA plasmid was co-transformed with a donor template into a strain expressing Cas9 (Cas9 plasmid, pNA0519; and sgRNA expressing plasmid, pMAH269). Successful editing is indicated by the reduced killing phenotype of the guide RNA plasmid upon addition of a donor template. We observed successful editing in 100% of the clones examined by PCR genotyping (Figure S6A). *swr1Δ* histone humanized strains were generate as described above.

#### Mapping the inviability of macroH2A1 histone fold

In order to map the residues of macroH2A1-HF that were inviable in yeast we first divided the region corresponding to the core histone fold domain and C-terminal tail of macroH2A1 into seven arbitrary sub-regions (Figure S7B). We then swapped in these sub-regions of macroH2A1-HF into the chimeric fusion construct containing the *Hs*H2A and the N-terminal tail of macroH2A1-HF (pMAH338) and tested if each swapped in region of macroH2A1-HF obstructed the function of the chimeric histone in *S. cerevisiae* (function as measure of the frequency of 5-FOA^R^ colonies following histone plasmid shuffle; sub-region 1, pMAH397; sub-region 2, pMAH399; sub-region 3, pMAH401; sub-region 4, pMAH403; sub-region 5, pMAH405; sub-region 6, pMAH407; sub-region 7, pMAH409). We first performed single sub-region swap experiments and found that sub-region 3 had the strongest negative effect on *Hs*H2A function (Figure S7D). Three additional sub-regions (two, four, and six) had less detrimental effects, but were still significantly less fit than the base construct (Figure S7D). Combining these sub-regions in pairs (i.e., regions 2+3) resulted in total failure to complement (sub-regions 1+2, pMAH411; sub-regions 2+3, pMAH413; sub-regions 3+4, pMAH415; sub-regions 4+5, pMAH417; sub-regions 5+6, pMAH419; sub-regions 6+7, pMAH421;), suggesting that multiple residues underly the inviability of macroH2A1-HF (Figure S7D).

We then performed single residue swap-backs within each of the inviable sub-regions in order to identify the specific residues responsible for the inviability of macroH2A1-HF. These experiments were carried out as “swap to rescue” (See Table S2 for detailed plasmid list), where we swapped each residue within the inviable sub-regions of macroH2A1-HF back to the *Hs*H2A residue (Figure S7E–I). For sub-region 3, we mapped the entirety of the inviability to residue Tyr38, which is part of the L1-loop interaction between H2A-H2B dimers (Figure S7E)^17, 67^. Furthermore, introduction of the Tyr38Glu swapback into the various inviable sub-regions resulted in only a partial rescue to the viability of each (Figure S7F). For example, introduction of Tyr38Glu significantly increased the average 5-FOA^R^ of region two from of 2.63e^-6^ to 3.52e^-5^ (Figure S7F). However, for sub-region four, introduction of Try38Glu did not lead to any significant improvement (from 7.97e^-6^ to 4.05e^-6^; Figure S7F). These data suggested that Try38Glu swap-back alone is necessary but not sufficient to rescue the inviability of macroH2A1-HF. By continuing to map the inviable residues for sub-regions two, four, and six we were able to identify the inviable residues of sub-regions two and four, but could not single out any one residue for region six (Figure S7G–I). Collectively, the inviable residues were either involved in interactions between the H2A-H2B dimers (Tyr38), between H2A and the DNA phosphate backbone (Lys32 and Arg74), the docking domain (Gln92), and near the DNA entry/exit site (residues 110 to 115) – suggesting that mutating these residues to the corresponding H2A residue helps to overcome the increased stability of macroH2A1 nucleosomes(Figure 3B)^68^.

#### RNA extraction and sequencing

RNA was extracted, sequenced, and data was analyzed as previously described^31^. Libraries were sequenced on an Illumina NextSeq 500 with paired end 2 x 150 bp read chemistry. We generated ∼25 million reads per sample. We defined up- and down-regulated genes in histone humanized yeast with macroH2A1 chromatin as genes with log2 fold-change <-1 or >1 compared to WT yeast and a false discovery rate adjusted *p* value <0.01. Gene enrichment analysis was done using the webtool ShinyGO (v 0.77)^69^.

#### MNase digestions and sequencing

Yeast strains were grown overnight at 30°C in YPD to saturation. The following day cultures were diluted to a A_600_ of 0.2 in 100 mL YPD media and grown to a A_600_ 0.8–1.0 at 30°C. Cells were then cross-linked by adding 2.7 mL of Formalin (final concentration of 1%) and incubated at 25°C with shaking for 15 minutes. To quench the formaldehyde, 5 mL of 2.5 M glycine was added and incubated for 5 minutes. Cells were then collected with centrifugation at 3000 x g for 5 minutes at 4°C, washed twice with ice-cold water. Pellets were either immediately processed or snap-frozen with liquid nitrogen and stored at -80°C.

Cells were resuspended in 1 mL of spheroplasting buffer (1.2 M sorbitol, 100 mM potassium phosphate pH 7.5, 1 mM CaCl_2_, with freshly added β-mercaptoethanol (0.5 mM) and 1 mg/mL Zymolyase 100T. Zymolyase digestion were monitored for production of spheroplasts. Spheroplast were collected by centrifugation at 3000 x g for 5 minutes, washed once in spheroplasting buffer and resuspended in 500 µL of MNase digestion buffer (1M sorbitol, 50 mM NaCL, 10 mM TRIS-HCL (pH 7.4), 5 mM MgCL2, 0.5 mM spermidine, 0.075% NP-40, with freshly added β-mercaptoethanol (1 mM) and either 2 units/mL or 0.2 units/mL MNase). Reactions were incubated for 45 minutes at 37°C and stopped by the addition of 16.6 µL of 0.5 M EDTA (30 mM final). Crosslinks were reversed by the addition of 12.5 uL 20% SDS (0.5% final), 12.5 µL proteinase K (20 mg/mL), and incubated for 1 hour at 37°C and for two hours at 65°C. Digested DNA was extracted with two rounds of phenol-chloroform extraction and DNA was precipitated with isopropanol. DNA was resuspended in TE buffer with 1 mg/mL RNAse A and incubated at 37°C for 30 minutes. Finally, DNA was purified with the Zymo DNA clean and concentrator kit according to the manufacture’s specifications.

Digested DNA was used as the input for Illumina library preps using the NEB Ultra II kit following the manufacture’s specification. Libraries were sequenced on an Illumina NextSeq 500 with paired end 2 x 150 bp read chemistry. We generated approximately 21 million reads per sample.

#### Capillary electrophoresis and NRL estimate

Approximately 20 ng of MNase digested DNA was analyzed using the Agilent ZAG DNA analyzer system with the ZAG 135 dsDNA kit (1-1500 bp). The fragment length data was analyzed in MatLab. Oligonucleosome sizes (up to penta-nucleosomes) were estimated using the ‘findpeaks’ function in the signal processing toolbox. Nucleosome repeat length was calculated as the slope of the line passing through the estimated oligonucleosome lengths.

#### MNase sequencing data analysis

Demultiplexed reads were first analyzed with Trimmomatic (v0.39)^70^ to remove sequencing adaptors and then with FastQC (v0.11.4) to assess read quality. Processed reads were then aligned to the Scer3 genome (R64) using the Burrows Wheeler aligner (BWA) mem algorithm (v0.7.7)^71^. For the mononucleosome analysis, we filtered reads with estimated insert sizes in the range of 120–180 bp. Filtered reads were then used as input for mononucleosome analysis using the DANPOS (v2) pipeline^72^. Nucleosome peaks, binned at 10 bp, were called using the ‘Dpos’ algorithm to call positions relative to the WT samples. Next composite plots were made using the ‘Profile’ algorithm relative to the transcription start sites^73^. Mono-nucleosome occupancy relative to the transcription start sites of 5206 genes was analyzed^73^. First, we clustered the data using k-means cluster with (with k = 6), resulting in six classes of genes based on the relative positioning of nucleosomes from the TSS. The value of K was determined using the “elbow” method and using previously defined number of clusters as a guide^27^. Next, within each cluster, we sorted the genes by their z-score normalized RNAseq transcript abundance in wildtype yeast (Figure S9F).

Nucleosome repeat length of each gene, relative to the +1 nucleosomes, was calculated by taking the slope of the line running from the mononucleosome fragment length to the pentanucleosome fragment length. We only consider those genes in groups 1, 2, 4, and 6, as those showed good phasing across all strains. Density plots of these NRL values were generated in MatLab using the ‘ksdensity’ function. To analyze the fragment lengths, we aligned processed reads to the Scer3 genome (R64) using Bowtie2 algorithm. Alignments were down sampled to ∼2 million reads for each sample and then used to generate Vplots and profile plots using the R package VplotR^74, 75^.

#### Nucleosome positioning analysis of differentially expressed genes

Differentially expressed genes in either histone humanized H2Amacro1 or macroH2A1-HF-sb were defined as log2 fold change >1 and <-1 with an adjusted *p*-value <0.01 (a total of 572 genes). The nucleosome occupancy of each gene (binned by 10 bp), relative to its transcription start site, was then sorted into clusters as before with k-means clustering. We excluded genes from clusters 3 and 5 as these genes did not exhibit well-phased nucleosomes, leaving us with a total of 268 genes (114 up-regulated and 154 down-regulated). For each cluster we determined the average relative nucleosome position for six nucleosomes downstream of the TSS in wildtype yeast (yeast with yeast histones). We then defined a window of 200 bp around each mean nucleosome position and then using these coordinates determined the position of the maximum peak for each nucleosome from every gene (totaling 684 nucleosomes for down-regulated genes and 924 nucleosomes for up-regulated genes). These positions were then plotted relative to the mean position for the wild-type nucleosome.

We examined the percent change in nucleosome occupancy in the nucleosome depleted region for all 5206 genes with annotated TSS. The NDR was defined as the region +50 bp from the -1 nucleosome to -50 bp from the +1 nucleosome (Figure S11B). We calculated the relative change in nucleosome occupancy to WT yeast (with *Sc* histones) as a percent change. We then examined NDR occupancy by sorting genes by their z-score normalized expression levels in WT yeast (Figure S11A, E). Lastly, we used the top and bottom 15% most/least abundant genes to compare the relative log2FC expression changes in histone humanized yeast relative to WT (Figure S11C, G). Protein-protein interactions (PPI) were determined by constructing a PPI network for the top 15% genes using the String algorithm (Figure S11D, H).

We examined the DNA shape feature propeller twist, which has been shown to correlate well with both nucleosome positioning of INO80-set nucleosomes and overall DNA rigidity^25^. We calculated the genome-wide propeller twist for the R64-2 genome build of *S. cerevisiae* using the R package DNAshapeR. The resulting DNA-shape was binned with a 5-bp rolling average and composite plots were constructed relative to the TSS or the dyad of the +1 nucleosome.

#### HiC libraries and analysis

Details on methodology for HiC data generation and analysis can be found in Lazar-Stefanita *et al.* (Co-submitted).

#### Whole genome sequencing

Genomic DNA was extracted as previously described and Illumina sequencing libraries were made with the NEB Ultra II FS kit^31^. Libraries were sequenced on an Illumina NextSeq 500 with paired end 2 x 36 bp read chemistry, generating ∼16 million reads per sample. Single nucleotide variant analysis, ploidy levels and chromosome coverage maps were generated as previously described^31^. To construct the String interaction network, we filtered out genes with synonymous mutations and used the remaining list of mutant genes as input queries. The interaction network was constructed using functional and physical protein associations and the resulting network was clustered by MCL clustering with the inflation parameter set to 2. Breakpoint analysis of coverage data was done by thorough inspection in IGV genome browser.

#### Nanopore sequencing and analysis

Overnight yeast cultures of humanized macroH2A1-HF-sb clones 1 and 4 were pelleted (∼5 mL), washed in 1 x PBS and resuspended in 5 mL of spheroplast buffer (1 M sorbitol, 50 mM potassium phosphate, 5 mM EDTA pH 7.5) supplemented with DTT (5mM) and zymolyase (mg/mL) and shaken at 210 rpm for 1 hour at 30°C. Spheroplasts where centrifuged at 2,500 g at 4°C, gently washed with 1M sorbitol and incubated in proteinase K solution (25 mM final EDTA, 0.5% SDS, Proteinase K 0.5 mg/ml) for 2 hours at 65°C with gentle inversion every ∼30 minutes. Lysates were extracted twice with a 1:1 ratio of Phenol:Chloroform:isoamyl alcohol and pooled aqueous layers were treated with ∼10ug of RNase A for 30 mins at 37°C before an additional 1:1 extraction with chloroform:isoamyl alcohol. DNA was precipitated with 1/10 volume 3M sodium acetate (pH 5.2) and 2.5X volume of ice-cold 100% ethanol and inverted until DNA strands visually appeared. High molecular weight DNA was spooled using a pipette tip, transferred to a new tube containing 70% ethanol wash, dried and dissolved overnight in TE buffer (10 mM Tris-HCl pH 8.0, 1 mM EDTA).

High molecular weight gDNA was quantified using Qubit 1x dsDNA HS Assay reagent (Thermo, Q33231) on the Qubit flex Fluorometer. DNA samples were simultaneously tagmented and barcoded using Oxford Nanopore Rapid Barcoding kit (SQK-RBK004) according to the manufacturers protocol. Barcoded samples were pooled, cleaned and concentrated with SERA-MAG beads (Cytiva, 29343052). The library was immediately loaded onto a Minion R9.4.1 flow cell (SKU: FLO-MIN106.001) and sequenced using the Gridion Mk1 device for 46 hr.

Base calls were made with the Guppy high-accuracy model (v6.2.11). We sequenced to a depth of 21.5x for clone 1 and 42.6x for clone 4, with read N50’s of 12,279 bp and 12,999 bp, respectively, allowing us to confidently infer the breakpoints across repetitive Ty elements (typically ∼6 kb). Reads were first trimmed to remove barcode adaptors with Porechop and then aligned to the R64-2 Scer genome assembly using the Minimap2 aligner . Quality of alignments was assessed with Alfred^77^, confirming a high proportion of reads with secondary alignments (15.2% and 23.5% of total for clone 1 and clone 4, respectively). Structural variants were then called using the Sniffles^78^ and CuteSV^79^programs and the resulting vcf files were manually merged. Circos plots displaying chromosome coverage and translocations were made using the TBtools software package^80^. Analysis of the rearrangement regions were done using the Samplot program^81^ to visualize non-contiguously mapping reads.

*Movie S1.* **3D representations of HiC maps from WT and histone humanized strains**

Composite 3D maps are shown counter-clockwise; WT (white; *S*.*c*. histones), histone humanized replicative *Hs*H2A (gray; *H*.s. histones), histone humanized macroH2A1-HF-sb (blue; *H*.s. histones), and histone humanized H2Amacro1 (cyan; *H*.s. histones). Centromeres are marked as yellow-colored spheres, telomeres are black colored spheres, and rDNA as pink colored spheres.

## Acknowledgments

We are thankful to members of the Boeke lab for their helpful critiques and comments on the data presented here, to Dr. Chris Todd Hittinger and Dr. Emilyclare Baker for the *Saccharomyces eubayanus* strain yHEB1515, from which we sourced the histone promoters and genes. This work was supported by NIH (NIGMS) fellowship F32GM116411 to D.M.T., a Genome Integrity Training Program NIH (NIGMS) T32GM115313 to M.A.B.H., and a Rules of Life: Epigenetics 2 grant from the NSF (award number: 1921641) to J.D.B.

## List of Supplementary Tables

*Table S1.* List of Strains used

*Table S2.* List of Plasmids used

*Table S3.* DNA oligos used in this study.

*Table S4.* Top Genetic interactors with macroH2A1 overexpression in yeasts.

*Table S5.* List of variants identified from whole genome sequencing of macroH2A1-HF-sb and H2Amacro1 histone humanized yeasts

*Table S6.* Global comparison of nucleosome positioning from combined biological replicates

*Table S6.* Differential gene expression analysis of histone humanized macroH2A1 (combined macroH2A1-HF-sb and H2Amacro1) vs. WT Sc histones

*Table S8.* Occurrences of aneuploidy in macroH2A1 histone humanized yeasts

*Table S9*. Putative Translocations identified in nanopore sequencing dataset in macroH2A1-HF-sb clone 1 and 4

---

**Figure S1.**
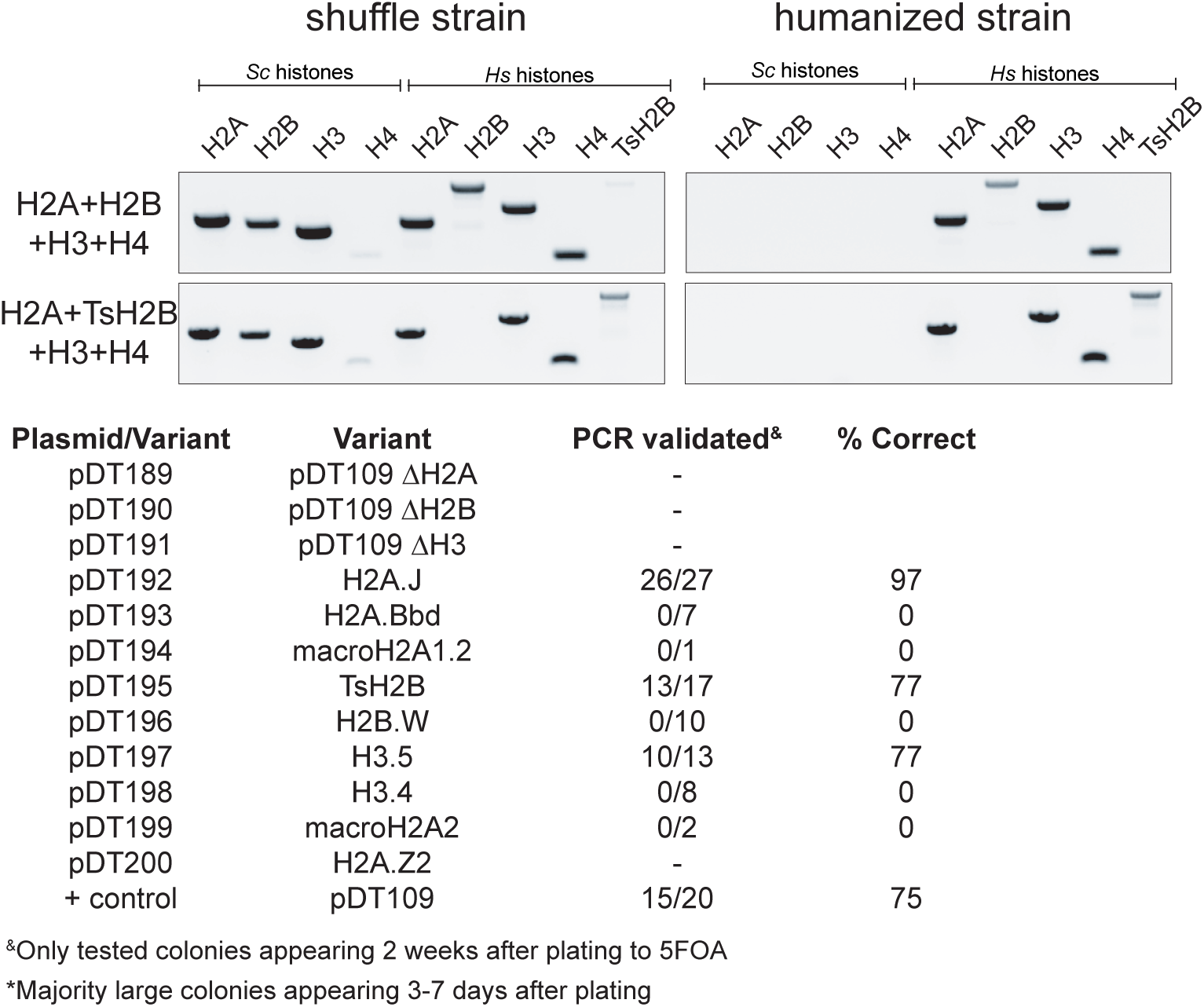
Validation of single gene complementation. Top; example PCR genotyping of humanized yeast with replicative histones and with TsH2B replacing replicative H2B. Below; summary table of PCR genotyping of colonies which emerged after two weeks of growth.

**Figure S2.**
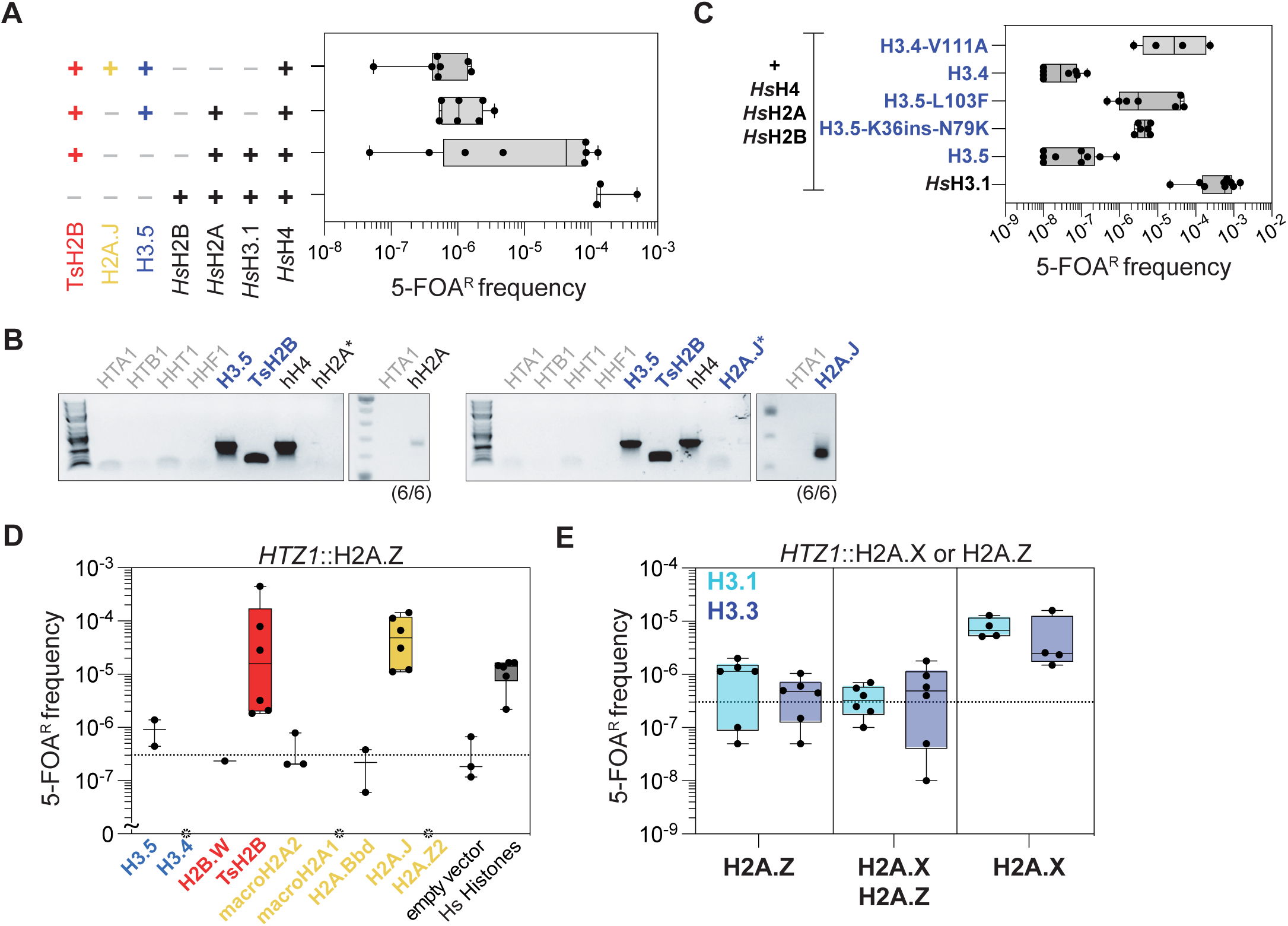
Additional histone humanizations with testis-specific variants. **(A)** Humanization assay of yeast with human testis-specific histones. **(B)** PCR genotyping of testis-specific histone humanized yeasts. Numbers in parentheses indicates number of positive clones out of total tested. **(C)** Humanization assays of H3.4 and H3.5 with nucleosome-stabilizing mutations and H3.5 with lysine residues.

**Figure S3.**
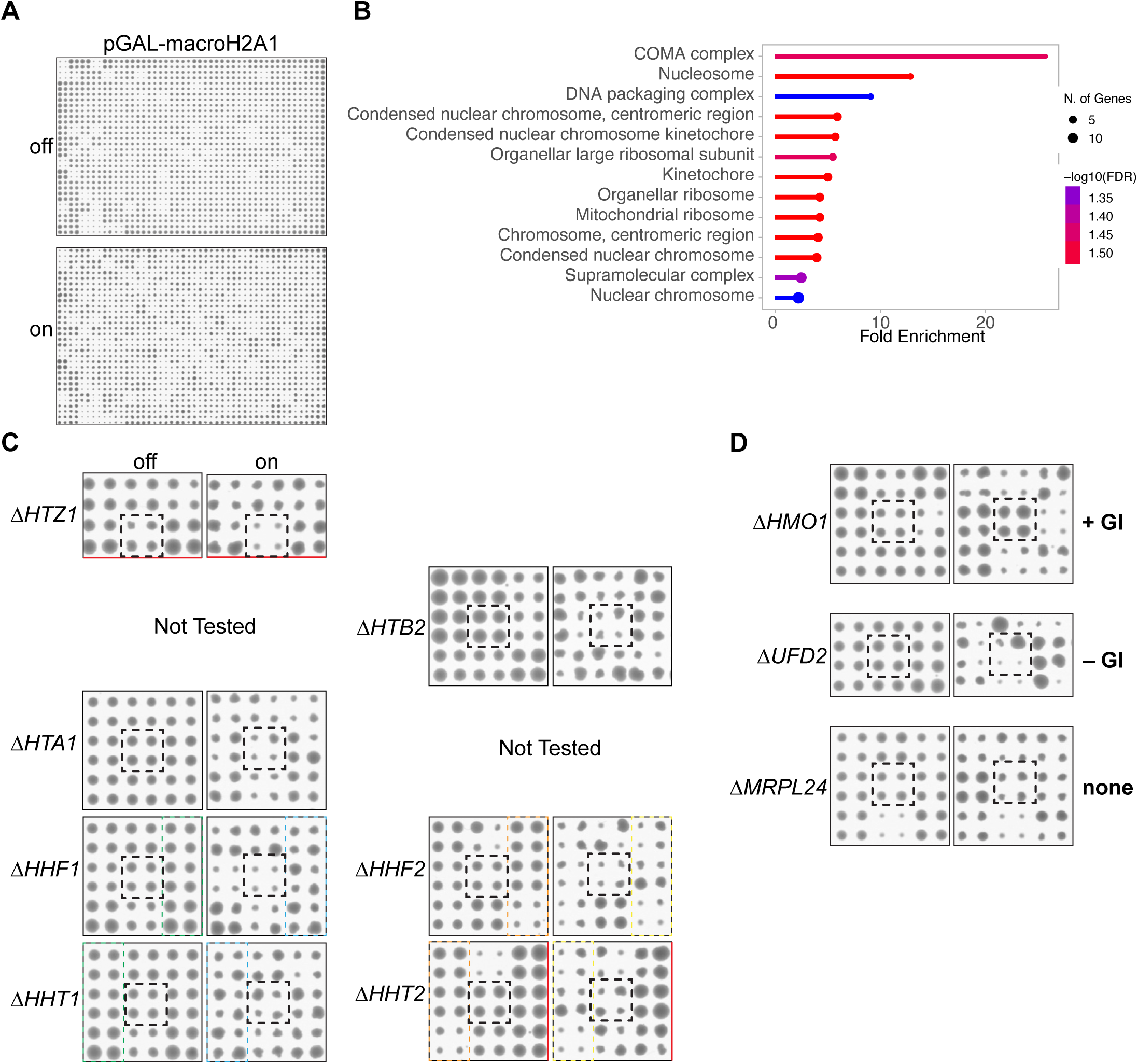
macroH2A1 genetic interaction screen. (A) Example images of screen without macroH2A1 expression (off) and with macroH2A1 expression (on). A universal donor strain (all centromeres tagged with a Ura3-pGAL-CEN) was used to transfer the macroH2A1 expressing plasmid (or empty vector) into the non-essential gene deletion collection through mating, followed by selection on galactose and 5-FOA to remove the universal donor strain’s chromosomes and select for non-essential gene deletions expressing macroH2A1. (B) GO biological processes enrichment of the synthetic sick gene deletions. (C) GIs of native histone genes with macroH2A1 expression. Note, HTB1 and HTA2 are not included in the list of non-essential genes and thus were not tested. Red marks near edge indicate the border of the growth plate. The colonies with the relevant genotype are outlined in a black dashed box. Dashed outlined areas indicated regions of the image that are shown across images. (D) Example GIs of either positive (HMO1), negative (UFD2) or no interaction (MRPL24).

**Figure S4.**
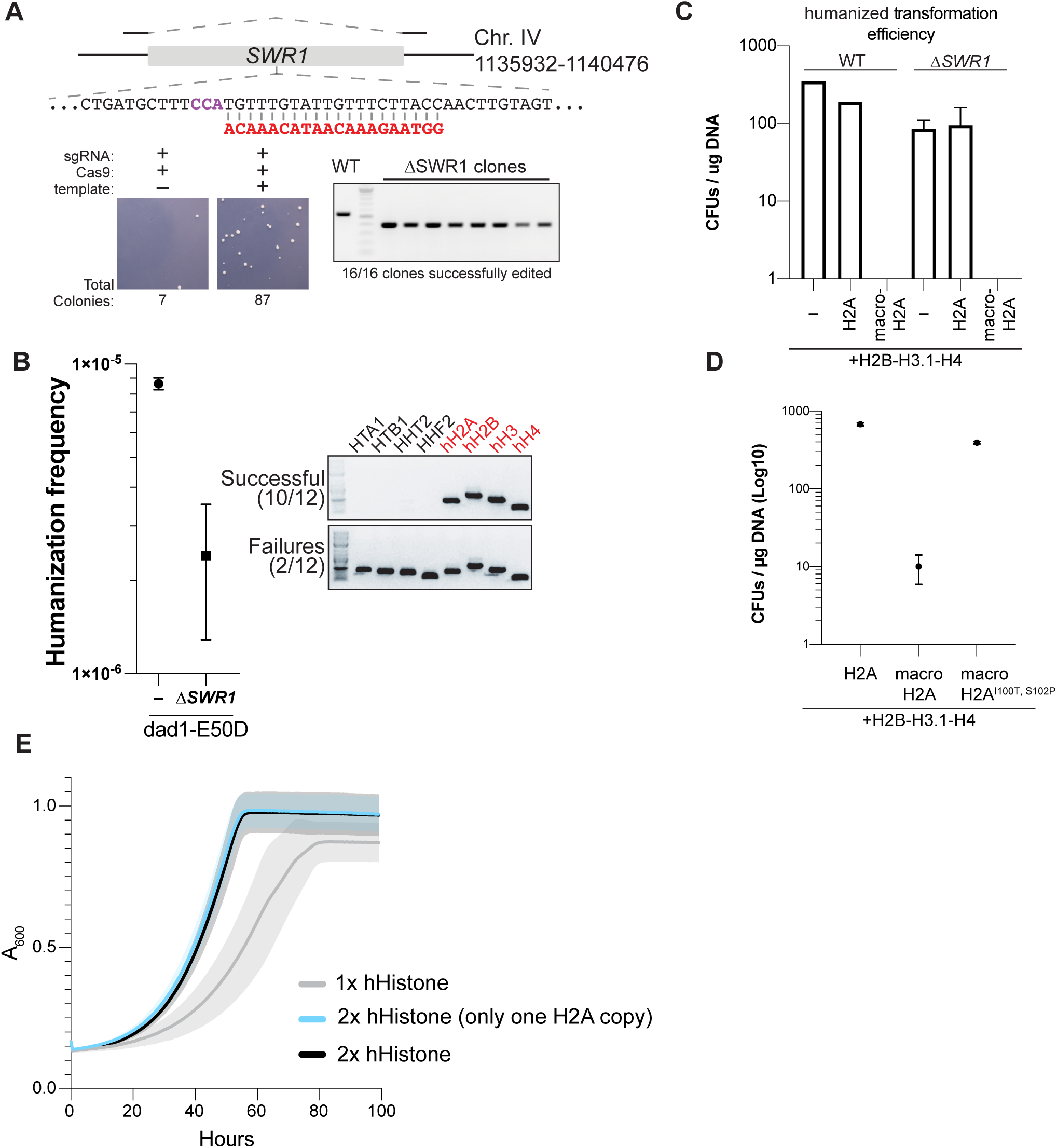
Swr1 complex does not catalyze deposition of macroH2A1 in yeast. **(A)** CRISPR-Cas9 editing strategy to delete *SWR1* in histone shuffle strain. **(B)** Humanization rates for *swr1*Δ histone shuffle strains with replicative human histones. **(C)** Colony forming unit (CFU) transformation assay of WT or *swr1*Δ histone-humanized yeasts. Transformation of plasmids encoding only *Hs*H3.1-*Hs*H4-*Hs*H2B in addition to either replicative *Hs*H2A or macroH2A1. macroH2A1 lowers transformation efficiency in either WT or *swr1Δ* strains, suggesting that Swr1 is not responsible for the toxicity of macroH2A1. **(D)** CFU transformation assay of histone-humanized yeasts with two mutations (I100T and S102P) in the C-terminal region of macroH2A1.

**Figure S5.**
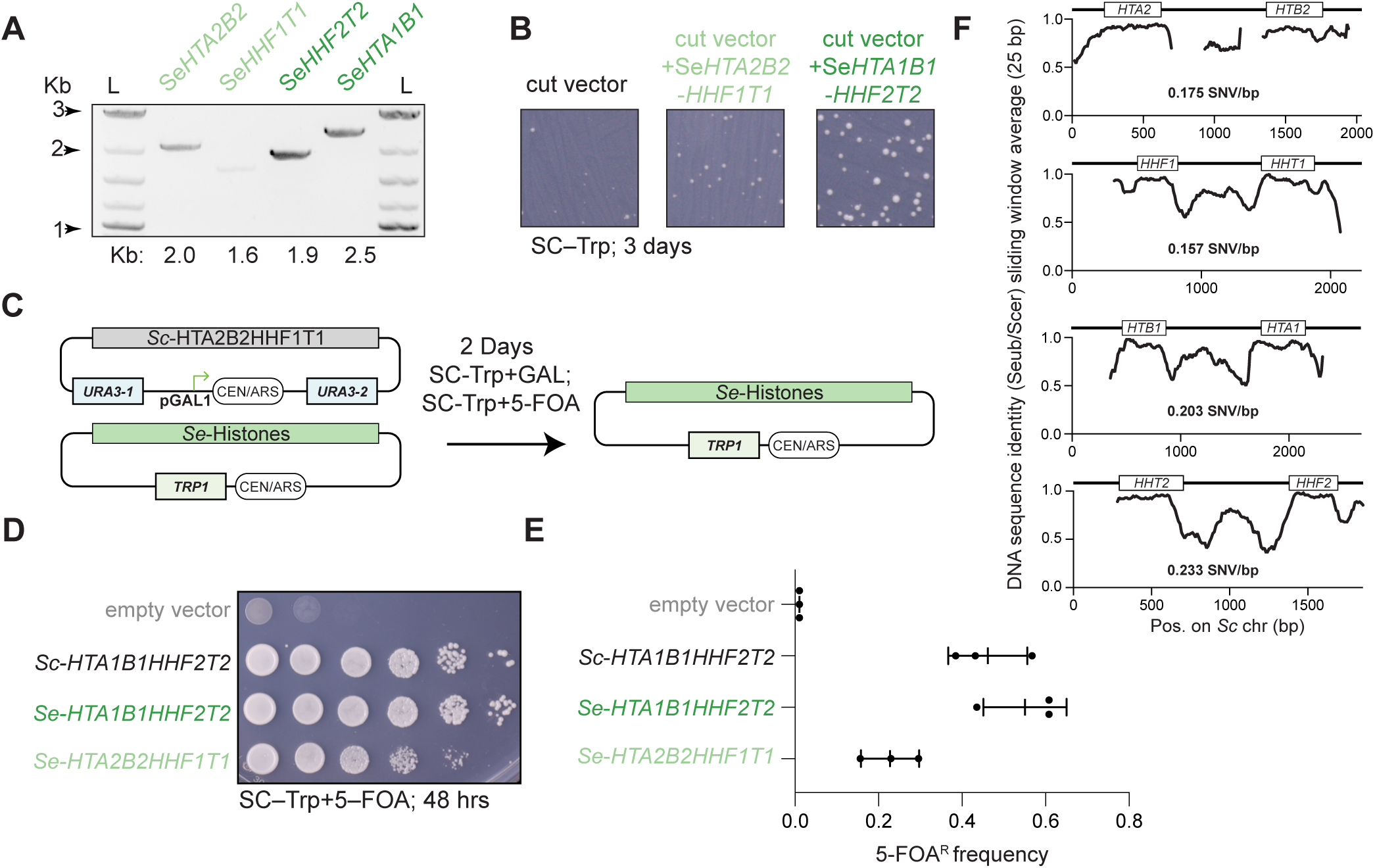
Repurposing of *S. eubayanus* replicative histones for use in *S. cerevisiae*. **(A)** PCR amplification of the native histone loci from *S. eubayanus*. Loci were amplified from the terminating sequences downstream of each histone gene (defined as 150 bp downstream of the stop codon). **(B)** In vivo assembly of expression vectors. **(C)** Overview of plasmid shuffle assay to test viability of *S. eubayanus* histones. **(D)** Spot assay of plasmid shuffle assay plated onto 5-FOA to counterselect the *URA3* (*S. cerevisiae* histone genes) plasmid **(E)** Quantification of biological replicates of shuffle assay **(F)** DNA sequence identity of histone gene clusters between *S. cerevisiae* and *S. eubayanus*.

**Figure S6.**
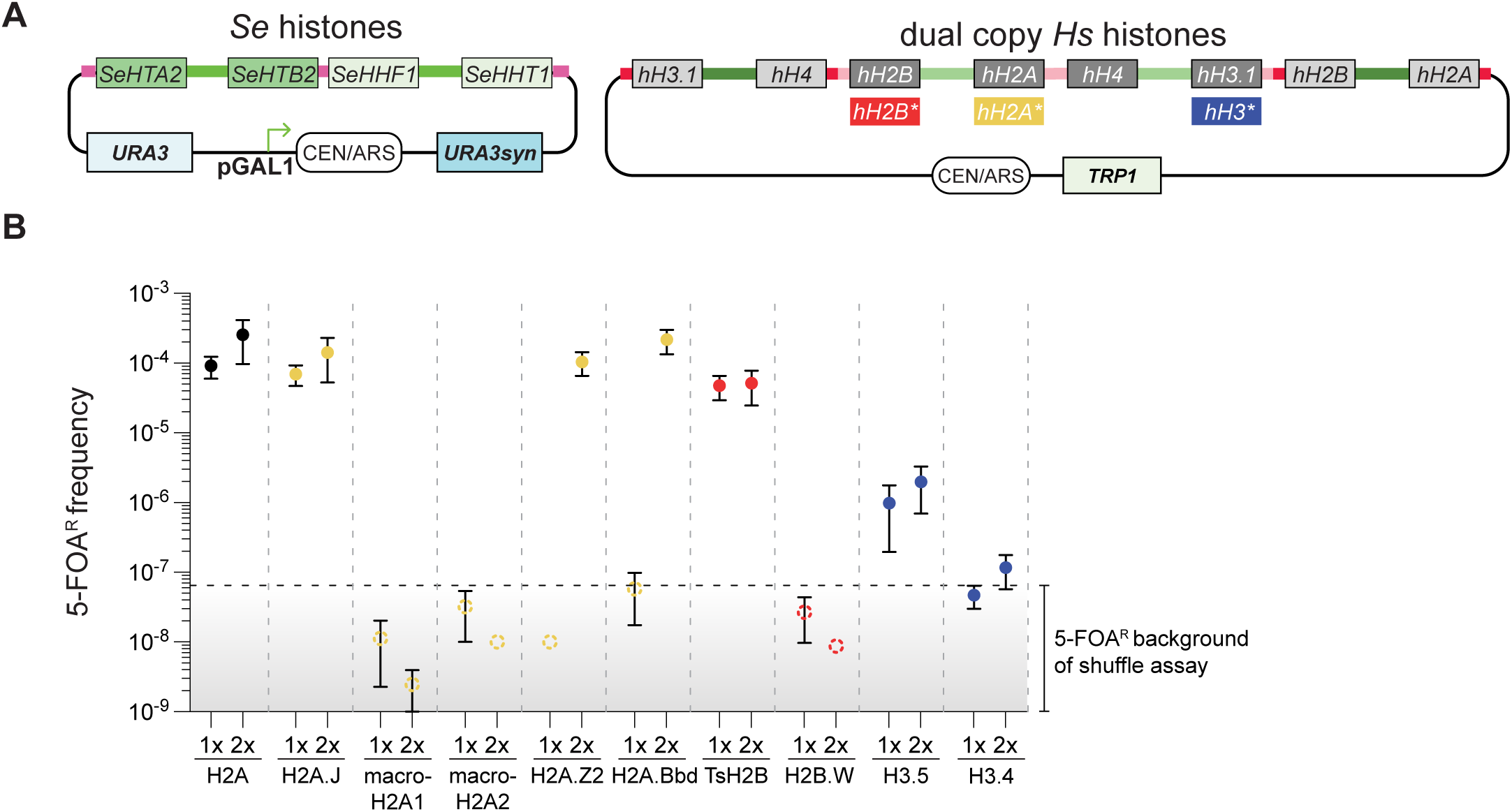
Epistatic interactions between replicative and non-replicative histones. **(A)** Overview of plasmid shuffle strategy with 2x human histone plasmids. The histone shuffle strain used carries a single set of *S. eubayanus* (*Se*) replicative histones (and histone promoters) encoded on a *URA3* counter-selectable plasmid. We then transformed in a plasmid encoding two copies of each human histone gene (each histone type encoded by two differently recoded genes), with some plasmids (as indicated) encoding a single non-replicative histone variant plus its associated replicative histone (e.g., replicative H2A + macroH2A1). **(B)** Humanization assay from 2x shuffle strategy. Note, results from Figure 1D are replotted to improve visual comparison.

**Figure S7.**
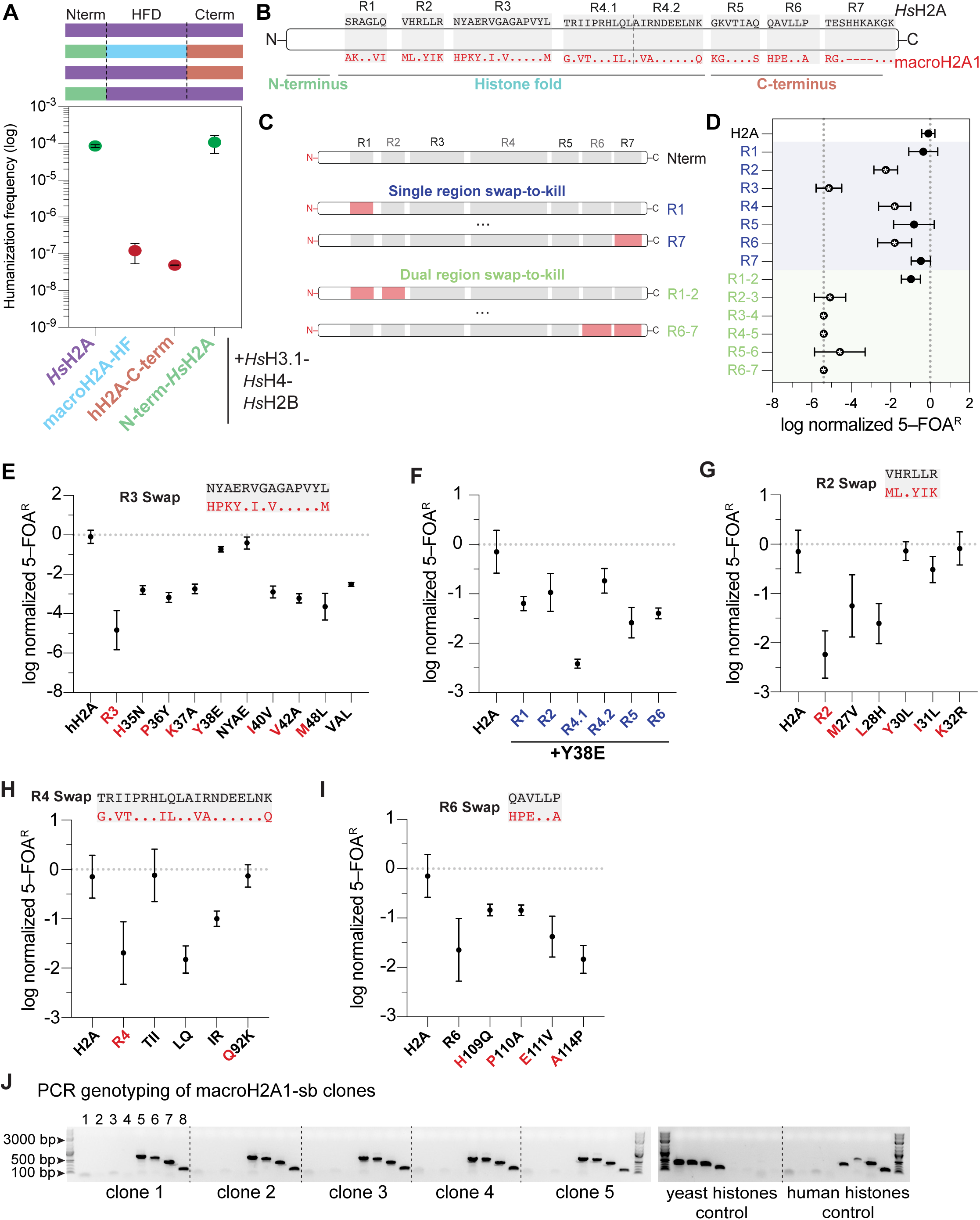
Dissecting inviable residues of macroH2A1 histone fold. **(A)** Humanization assay of chimeric histones of replicative H2A with variant histone macroH2A1. **(B–C)** Overview of regional swaps of the macroH2A1 histone fold domain. Replicative H2A sequence is shown in black above and macroH2A1 in red below. The N-terminus in this experiment was that of macroH2A1. **(D)** Humanization assay of “swap-to-kill” experiments. Regions marked with an asterisk significantly diminished the complementation of replicative H2A. **(E)** Humanization assay of swap-back experiments of inviable region 3 of macroH2A1. Rates of 5-FOA^R^ were log-normalized to the average 5-FOA^R^ frequency of replicative H2A. **(F)** Epistatic interactions of swapped-back region 3 (Y38E) with additional inviable regions of macroH2A1. **(G)** Humanization assay of swap-back experiments of inviable region 2 of macroH2A1. **(H)** Humanization assay of swap-back experiments of inviable region 4 of macroH2A1. **(I)** Humanization assay of swap-back experiments of inviable region 6 of macroH2A1. **(J)** PCR genotyping of humanized macroH2A1-HF-sb strains. Amplicons are as indicated, lanes 1 to 4, yeast H2A, H2B, H3, and H4, respectively; lane 5 macroH2A1/H2A, and lanes 6 to 8, *Hs*H2B, *Hs*H3, and *Hs*H4, respectively.

**Figure S8.**
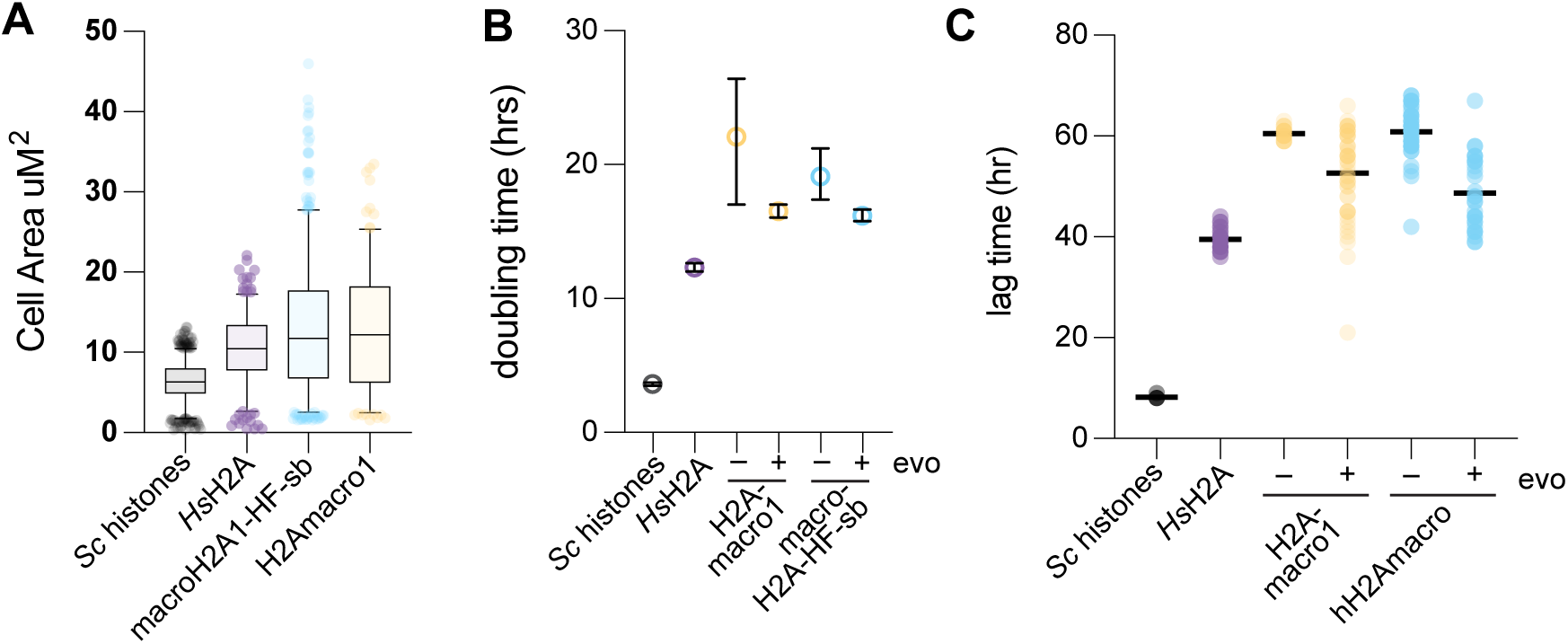
Cell size, doubling time and lag time of macroH2A1 humanized yeast. **(A)** Cross-sectional area quantifications of humanized yeasts and WT control. **(C)** Doubling time calculations from nonlinear regressions of the A_600_ data in log phase of growth, 95% confidence intervals around the mean doubling time are shown. **(C)** Lag time quantification of humanized yeasts and WT control.

**Figure S9.**
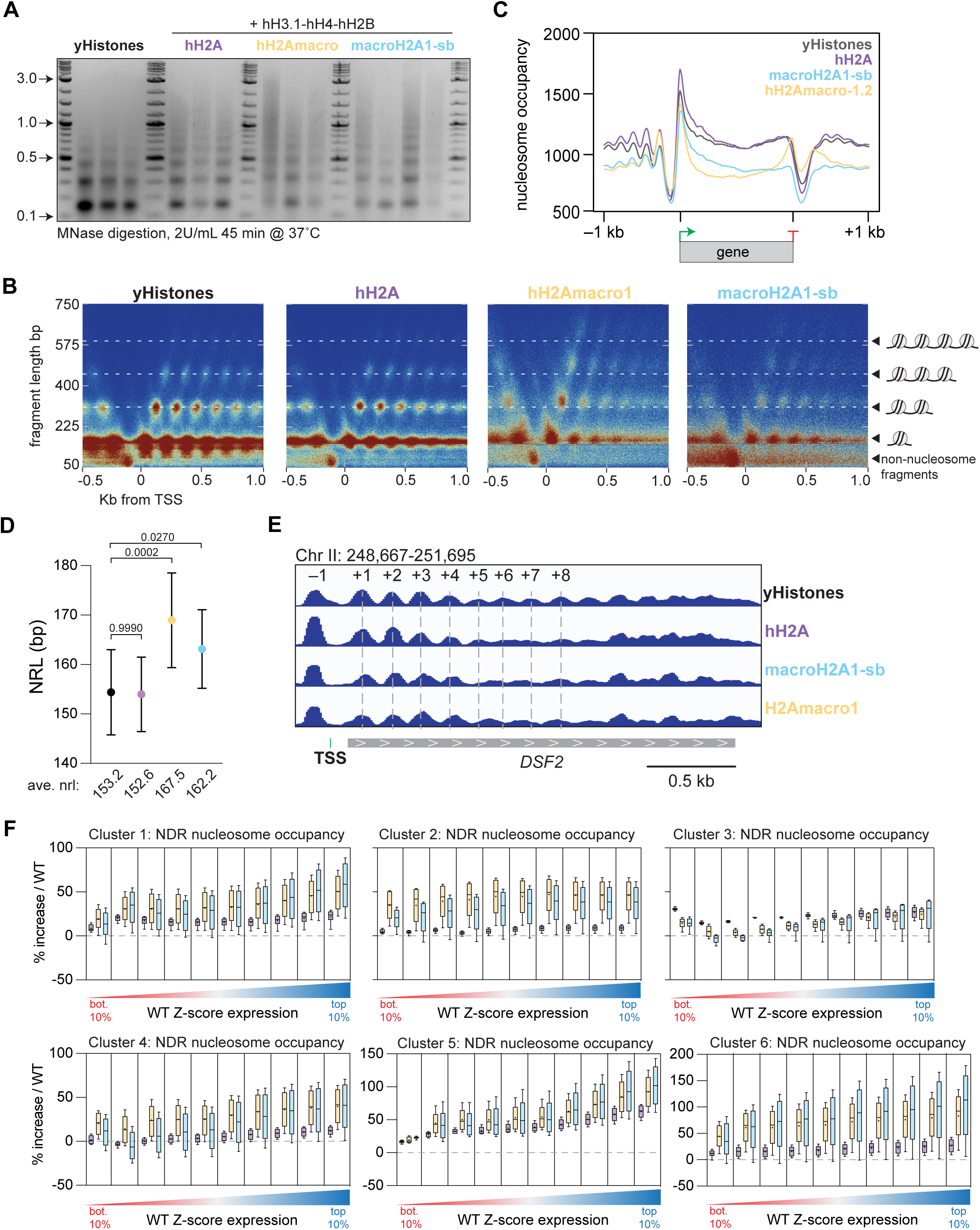
MNase digestions and MNase-seq analysis. **(A)** MNase digested chromatin of WT and humanized strains. Digested DNA was run on a 1% agarose gel and stained with ethidium bromide. **(B)** Composite plot of fragment lengths of sequenced MNase digested DNA binned relative the transcription start sites (TSS). **(C)** Metagene plot of nucleosome occupancy plus and minus 1 kb from the TSS and TTS. **(D)** Inferred nucleosome repeat length from capillary electrophoresis analysis. **(E)** Example gene track of nucleosome occupancy. **(F)** Quantifications of the percent change in nucleosome occupancies in the NDR in *Hs* histone yeasts versus *Sc* histone yeasts. Genes within each cluster were binned into groups corresponding to 10% intervals of the WT z-score expression levels (bottom 10%, genes with the least abundant transcripts; top 10%, genes with most abundant transcripts). Colors of boxes represent type of *Hs* H2A chromatin examined, as in panel A.

**Figure S10.**
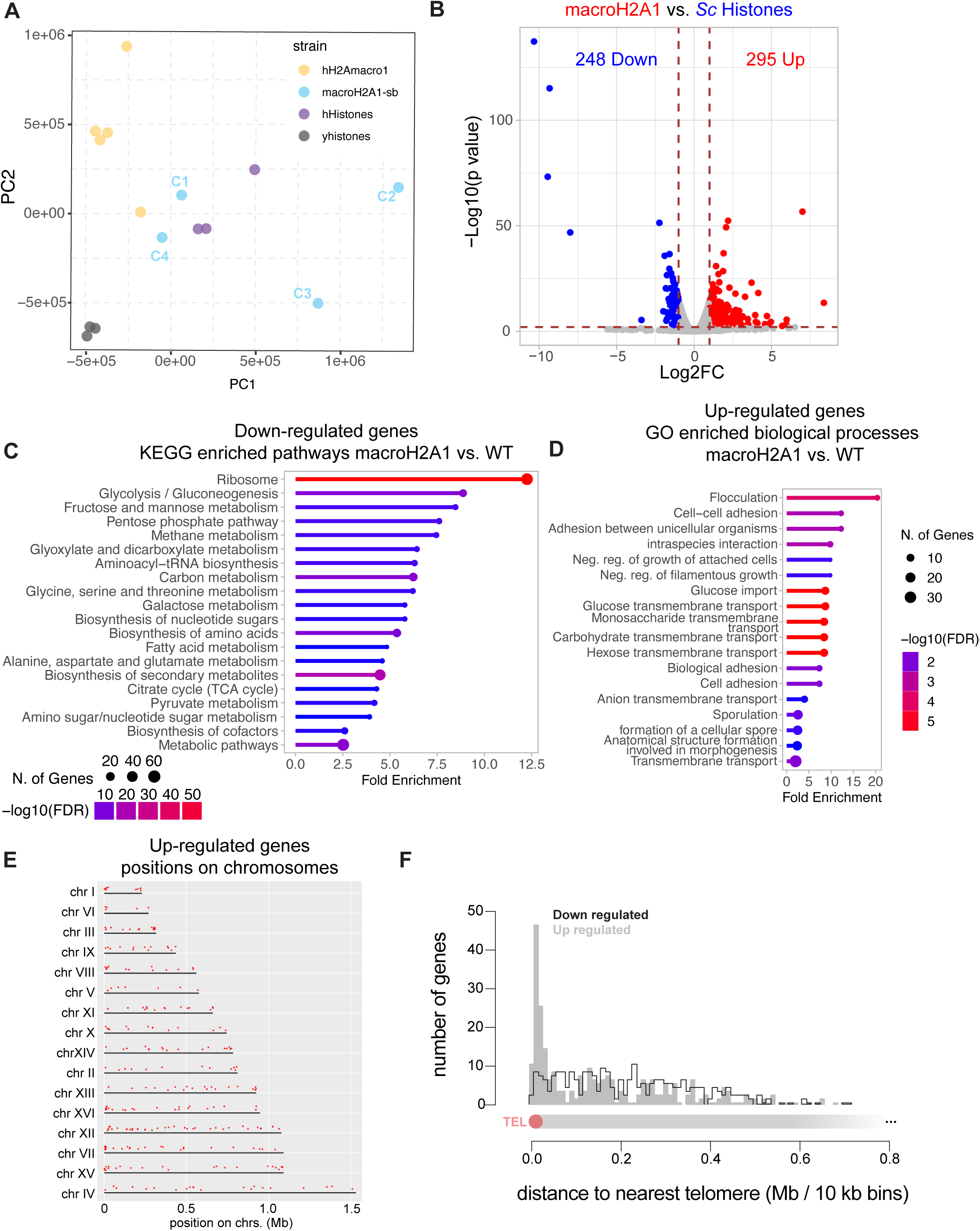
RNA-seq in macroH2A1 humanized cells. **(A)** PCA plot of PC1 and PC2 of transcriptomes of WT and histone humanized yeasts. **(B)** Volcano plot of differently expressed genes comparison between WT and macroH2A1 humanized yeasts. Genes with a log2FC of less than -1 or greater than +1 and adjusted *p*-value < 0.01 were considered significant. Blue genes are down-regulated and red genes up-regulated in macroH2A1 humanized yeast. **(C)** KEGG enrichment analysis of down-regulated genes in macroH2A1 humanized yeast. **(D)** GO biological processes enrichment analysis of up-regulated genes in macroH2A1 humanized yeast. **(E)** Positions of the down-regulated genes on each chromosome. **(F)** Distribution of the distance of the down-(solid gray) and up-(lined black) regulated genes relative the nearest telomere. Note, the general decline in gene density in gene’s >0.4 Mb is an effect due to the fact that most chromosomal arms in *S. cerevisiae* are <0.4 Mb in length.

**Figure S11.**
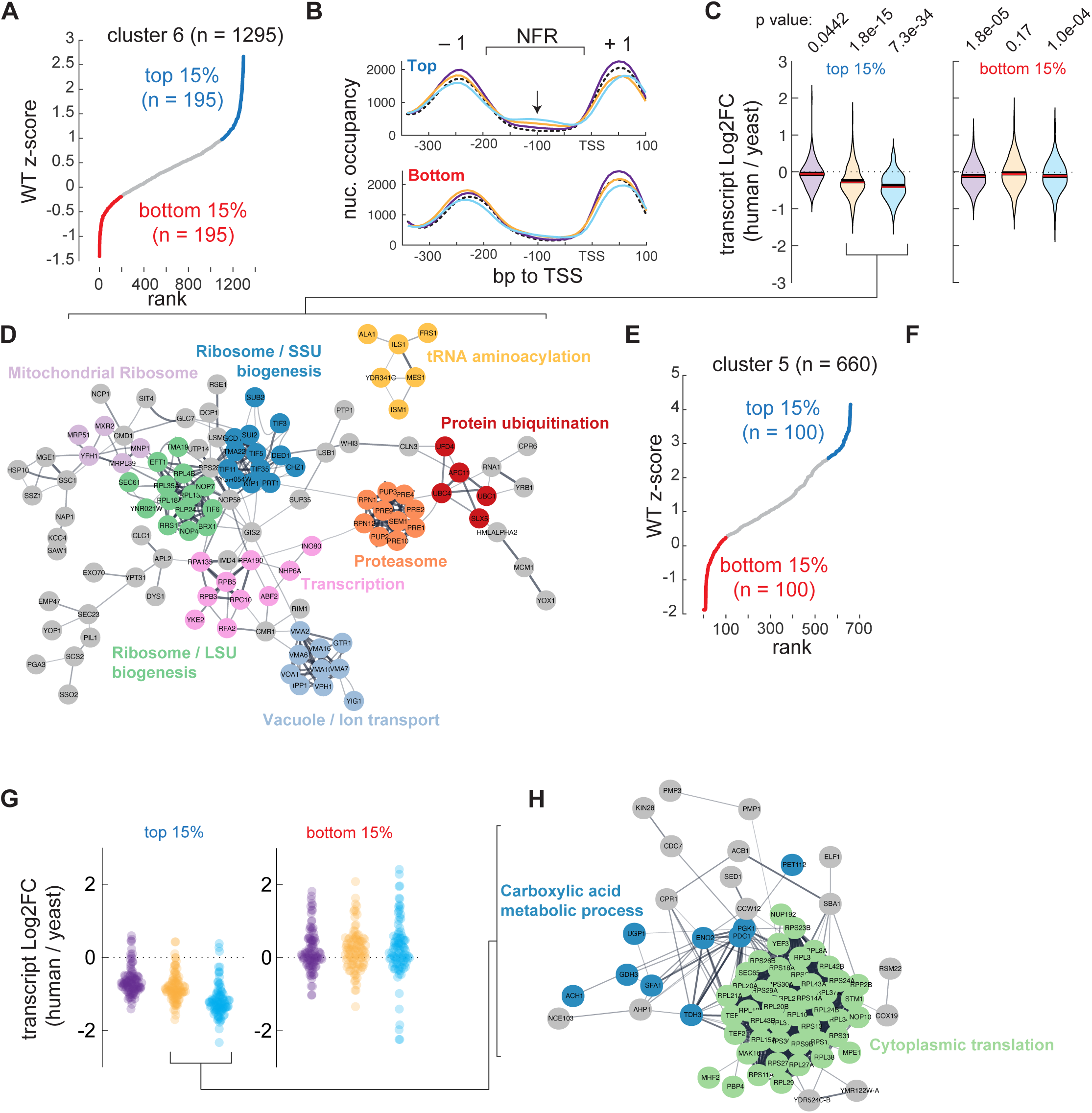
Global decrease of protein translation inferred from MNase-seq and RNAseq in histone humanized yeasts. **(A)** Z-score rank of transcript abundance in WT yeast of genes in cluster 6 of nucleosome occupancy. **(B)** Nucleosome occupancy in the nucleosome-depleted regions of either the top 15% most abundant transcripts or bottom 15% of cluster 6 genes (colored violin plots indicate the strain being compared to WT (*Sc* histones); purple, histone humanized (all replicative histones); yellow, H2Amacro1 histone humanized; and cyan, macroH2A1-HF-sb histone humanized. **(C)** Log2FC of the top 15% most abundant transcripts or bottom 15% of cluster 6 genes. **(D)** Protein-protein interaction network of down-regulated genes in panel C. **(E–H)** Same as before, but shown for cluster 5 genes.

**Figure S12.**
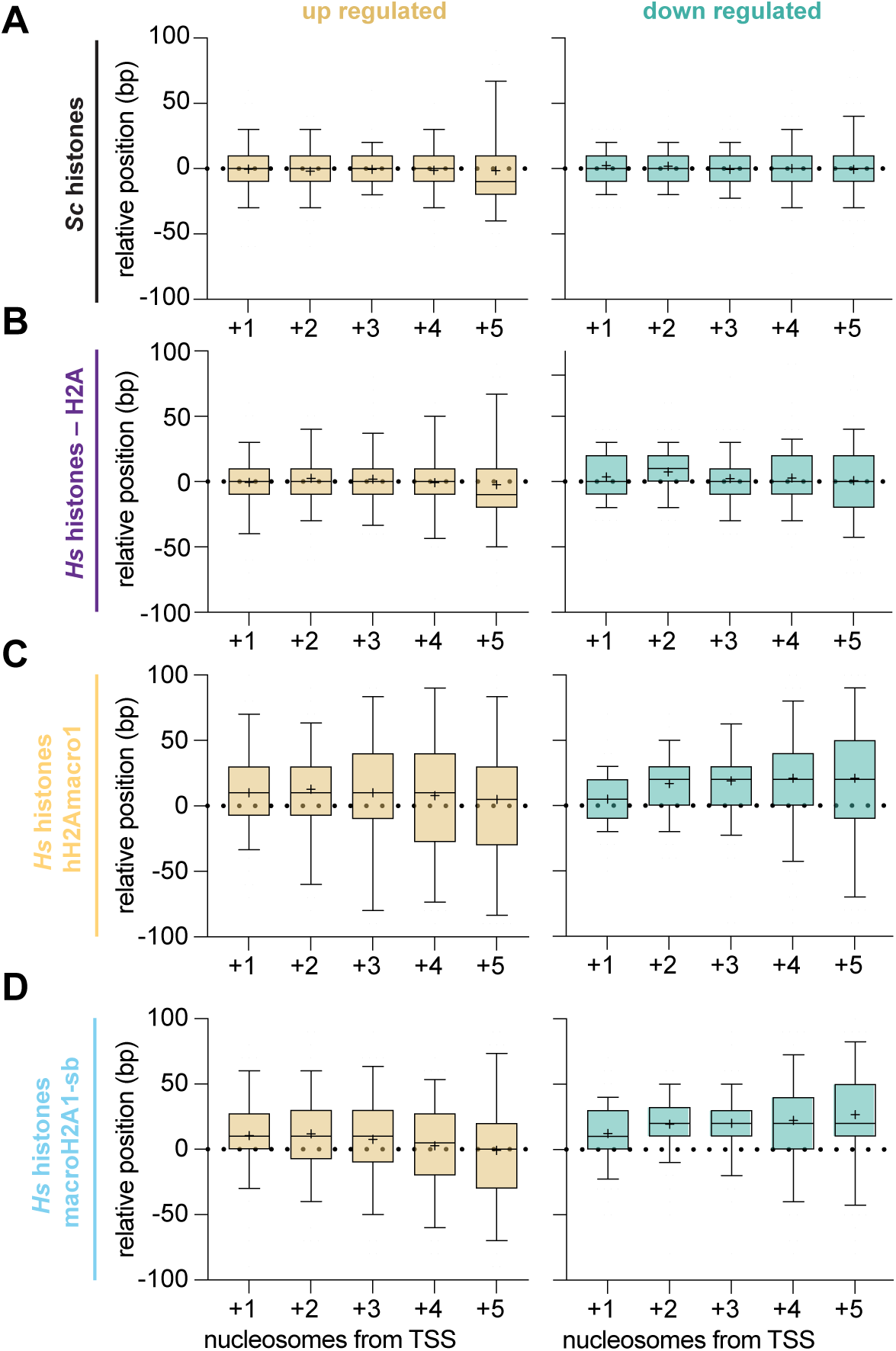
Relative nucleosome positioning downstream of the TSS. **(A-D)** Relative nucleosome positions for five nucleosomes downstream of the TSS for the up- or down-regulated genes in macroH2A1. Strain examined is as shown for each panel.

**Figure S13.**
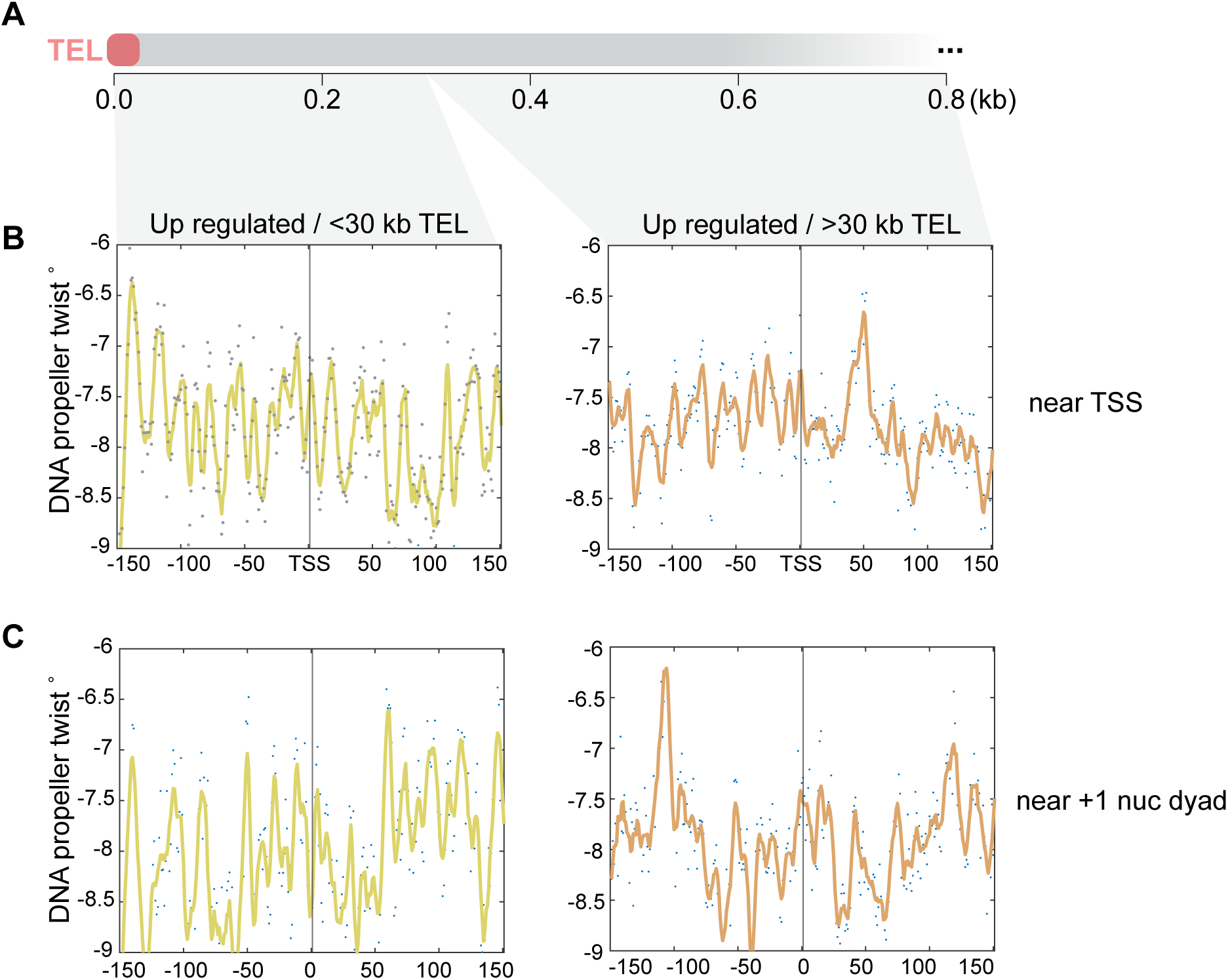
DNA shape features of up-regulated genes near to and far from subtelomeres. **(A)** Diagram of distance to telomeres (metachromosome plot). **(B)** DNA propeller twist near the TSS of up-regulated genes with subtelomeres and up-regulated genes outside of subtelomeric regions. **(C)** DNA propeller twist near the +1-nucleosome dyad of up-regulated genes with subtelomeres and up-regulated genes outside of subtelomeric regions.

**Figure S14.**
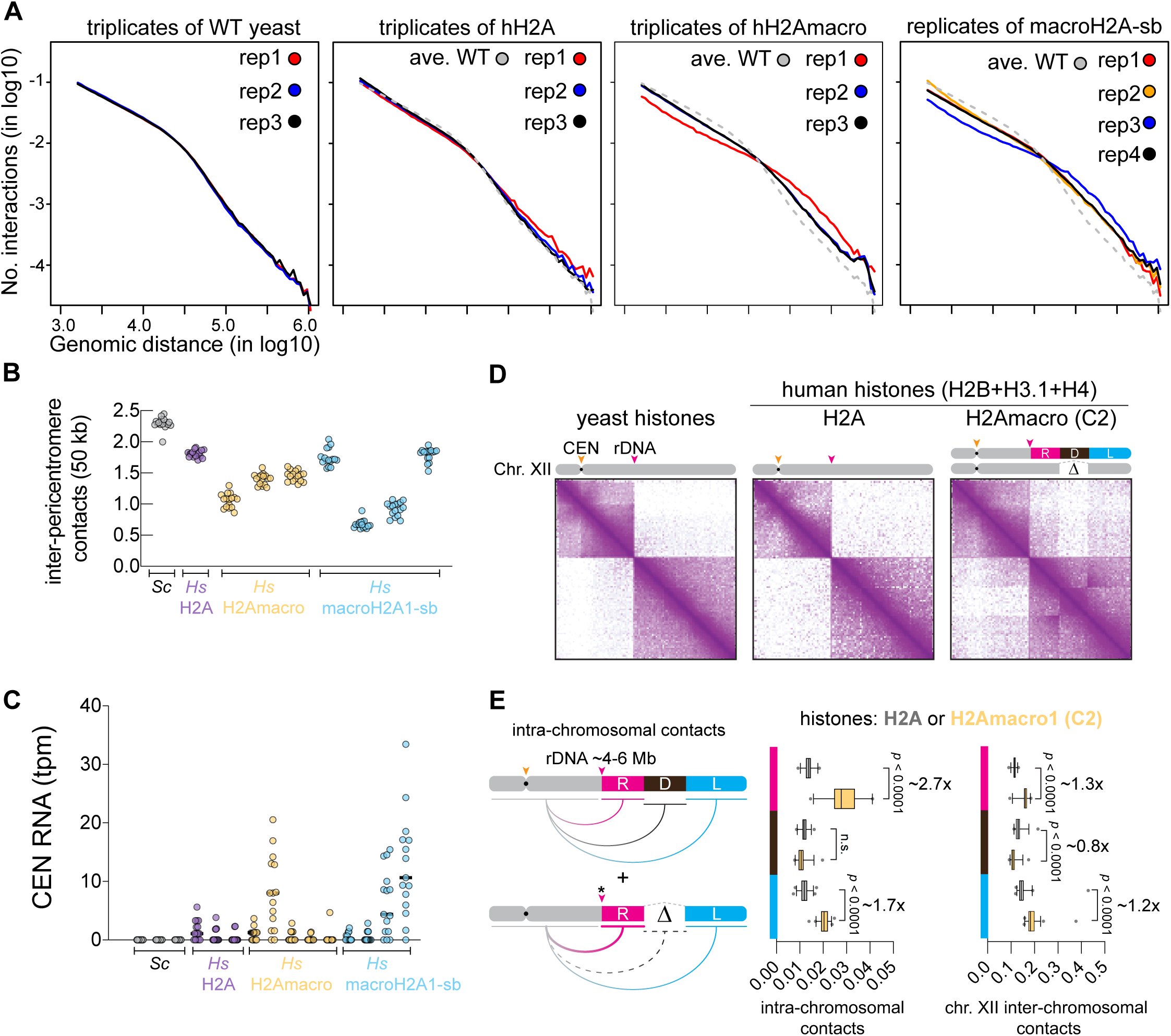
Clonal variation in genome stability of macroH2A1 humanized yeasts. **(A)** Contact probability decay plot (as in Figure 6B) of each replicate. The composite average plot of WT yeast is shown as gray dashed line in each plot of the histone humanized yeast strains. **(B)** Inter-pericentromeric contact quantifications from normalized Hi-C maps, plotted in 50 kb-windows centered on a given centromere. Each dot represents the sum of all contacts a given peri-centromere makes with the remainder 15 peri-centromeres. **(C)** Centromeric RNA quantification from RNA sequencing. **(D)** Chromosome *XII* Hi-C maps of WT, replicative H2A and H2Amacro1 (clone2) humanized cells. Positions of centromere (CEN) and rDNA locus are indicated. In H2Amacro1 (clone2) two copies of chr *XII* are present with one copy housing an internal deletion. Three regions downstream the rDNA array were annotated on chr *XII* schematic relative to the deletion: right region (R; pink), the deletion itself (D; black), and left of the deletion (L; blue). **(E)** Quantification of intra- and inter-chromosomal contacts in function of the internal deletion on chromosome *XII*. Left: shown are the three regions whose intra-chromosomal contacts with the left-rDNA flanking part of chr *XII* (gray) were quantified. Middle: quantification of intra-chromosomal contacts for each of the three regions in replicative H2A and H2Amacro1 humanized cells. The expected level of contacts between left-rDNA flanking region and the deleted region implies that the rDNA array on the wild-type chromosome is of similar size to replicative humanized rDNA array. Right: quantification of contacts between chr *XII* regions and the rest of the genome (inter-chromosomal). Note, the inter-chromosomal contacts increase in region L due to the clustering of telomeres. The inter-chromosomal contacts of chr *XII* account for the increased frequency due to ploidy increase of chr *XII*, however the magnitude of intra-chromosomal contact increase of region R or L are much larger than a ploidy increase would explain.

**Figure S15.**
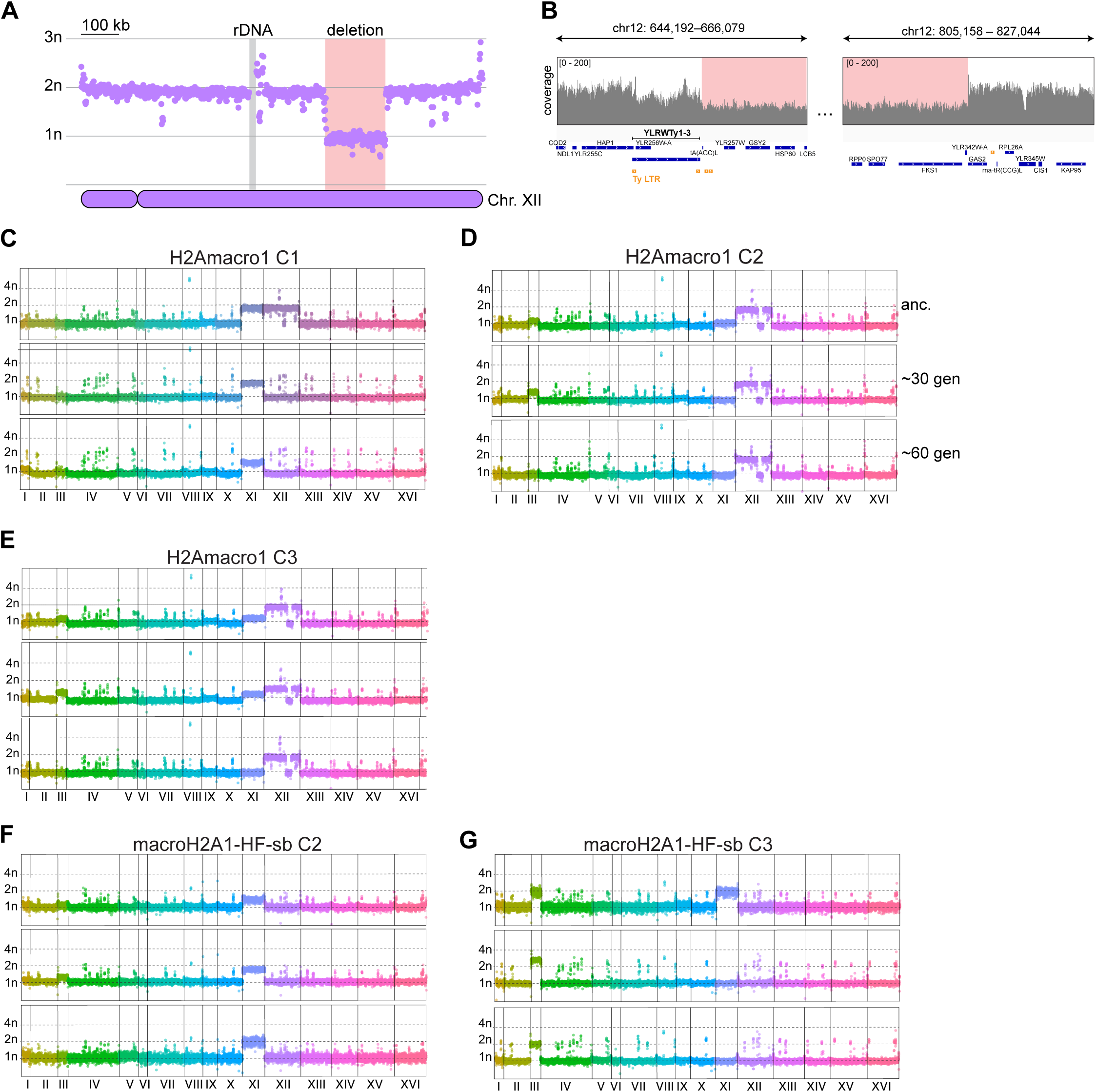
Clonal variation in genome stability of macroH2A1 humanized yeasts, continued. **(A)** Chromosome coverage plot of H2Amacro1 humanized clone 2 showing large internal deletion and aneuploidy of chromosomes XII. Deletion region is highlighted in red. **(B)** Zoomed in WGS coverage tracks of the regions near the break points of the internal chromosome XII deletion. **(C–G)** Example whole genome sequencing coverage plots of H2Amacro1 or macroH2A1-HF-sb humanized yeasts at differing time points.

