## Supplemental Figures for "Human macroH2A1 drives nucleosome dephasing and genome instability in histone-humanized yeast"

Figure S1. Validation of single gene complementation

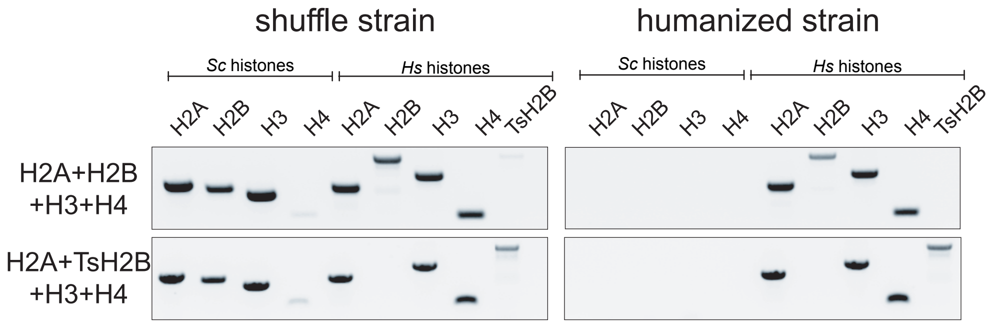


| Plasmid/Variant | Variant | PCR validated[&] | % Correct |
|---|---|---|---|
| pDT189 | pDT109 ΔH2A | - | |
| pDT190 | pDT109 ΔH2B | - | |
| pDT191 | pDT109 ΔH3 | - | |
| pDT192 | H2A.J | 26/27 | 97 |
| pDT193 | H2A.Bbd | 0/7 | 0 |
| pDT194 | macroH2A1.2 | 0/1 | 0 |
| pDT195 | TsH2B | 13/17 | 77 |
| pDT196 | H2B.W | 0/10 | 0 |
| pDT197 | H3.5 | 10/13 | 77 |
| pDT198 | H3.4 | 0/8 | 0 |
| pDT199 | macroH2A2 | 0/2 | 0 |
| pDT200 | H2A.Z2 | - | |
| + control | pDT109 | 15/20 | 75 |

[&]Only tested colonies appearing 2 weeks after plating to 5FOA

*Majority large colonies appearing 3-7 days after plating

Figure S2. Additional histone humanizations with testis-specific variants

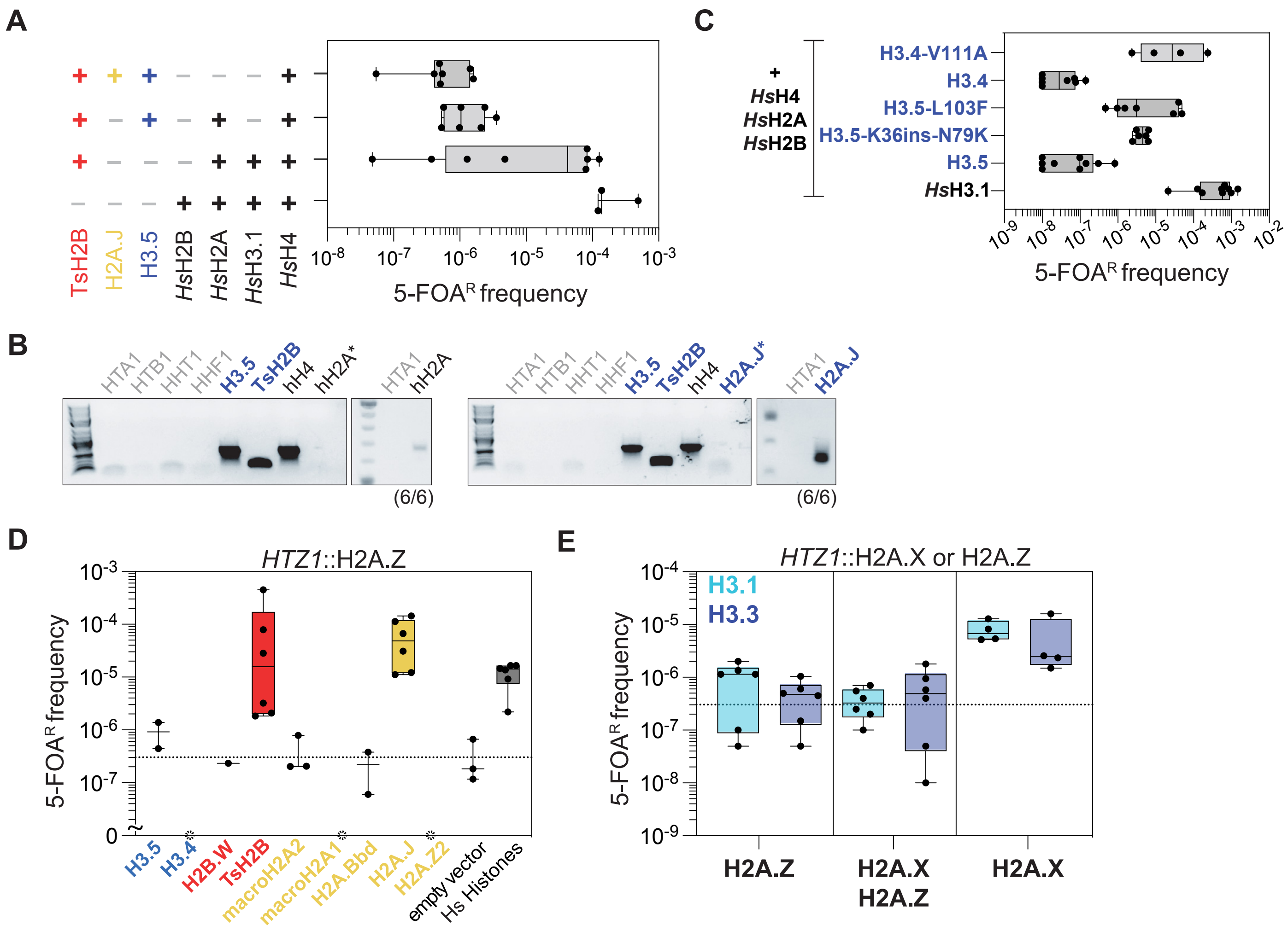

Figure S3. Genome-wide nonessential gene deletion interactions with macroH2A1 expression

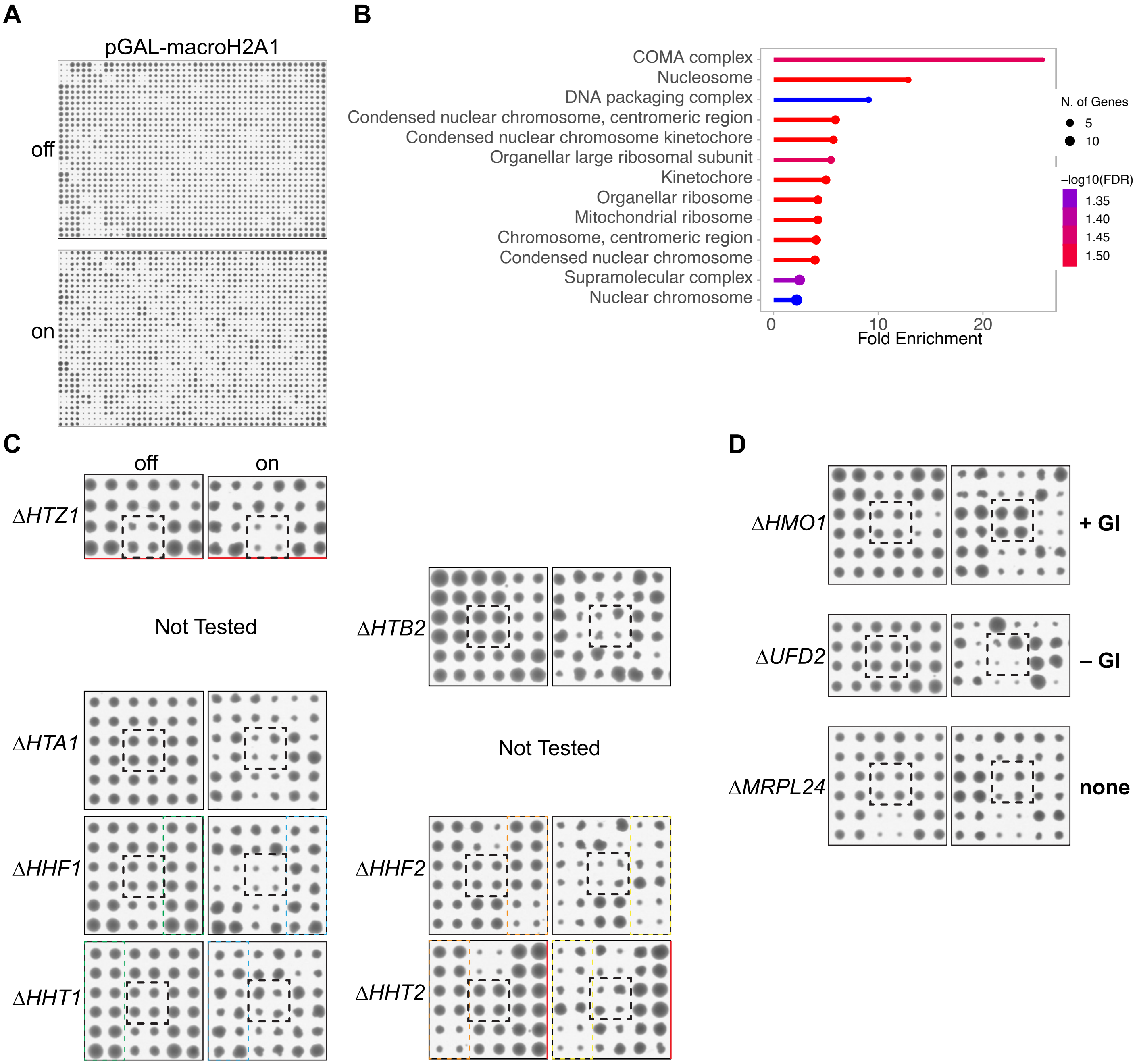

Figure S4. Swr1 complex does not catalyzes the deposition of macroH2A1 in yeast

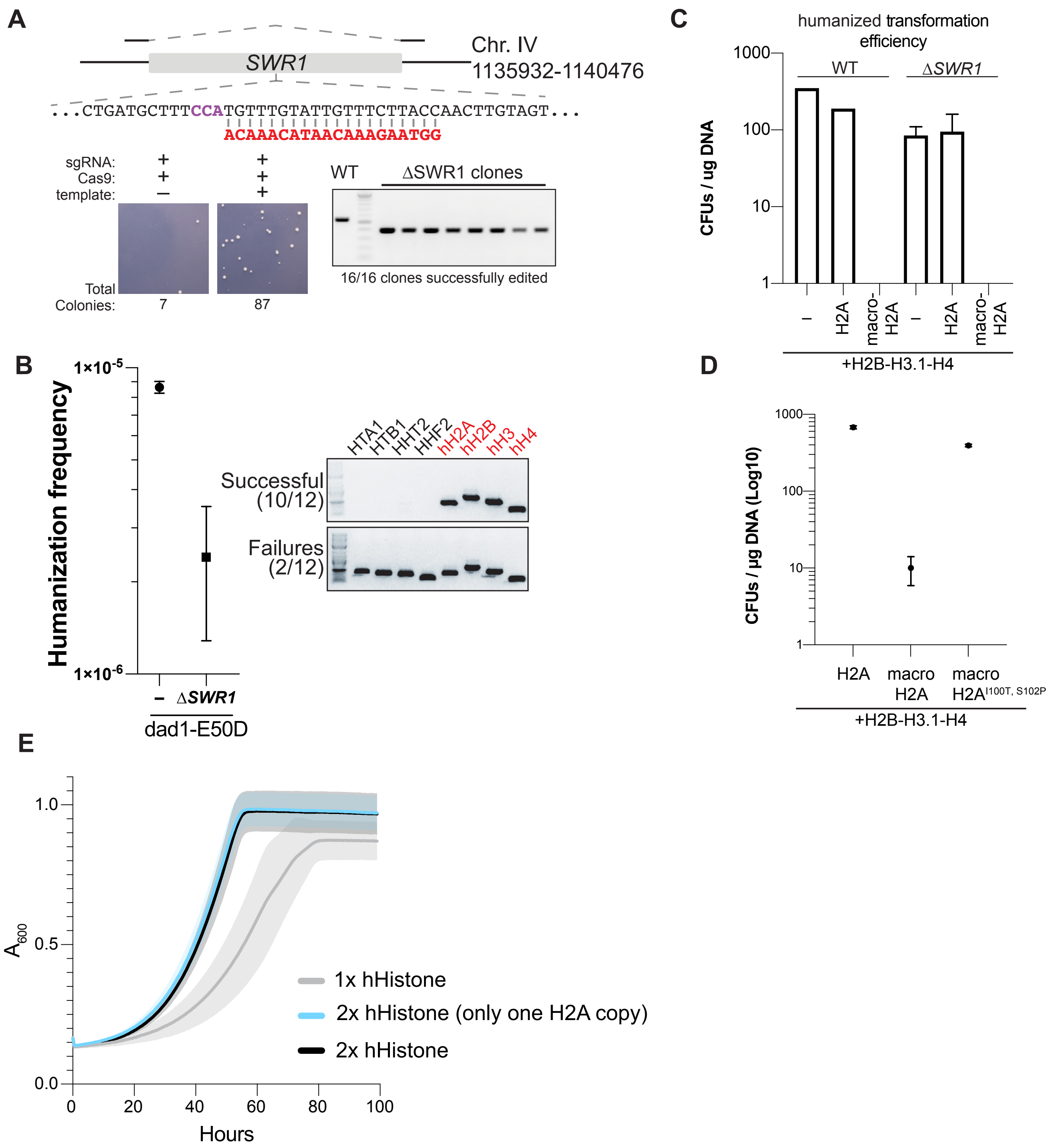

Figure S5. Repurposing of *S. eubayanus* replicative histones for use in *S. cerevisiae*

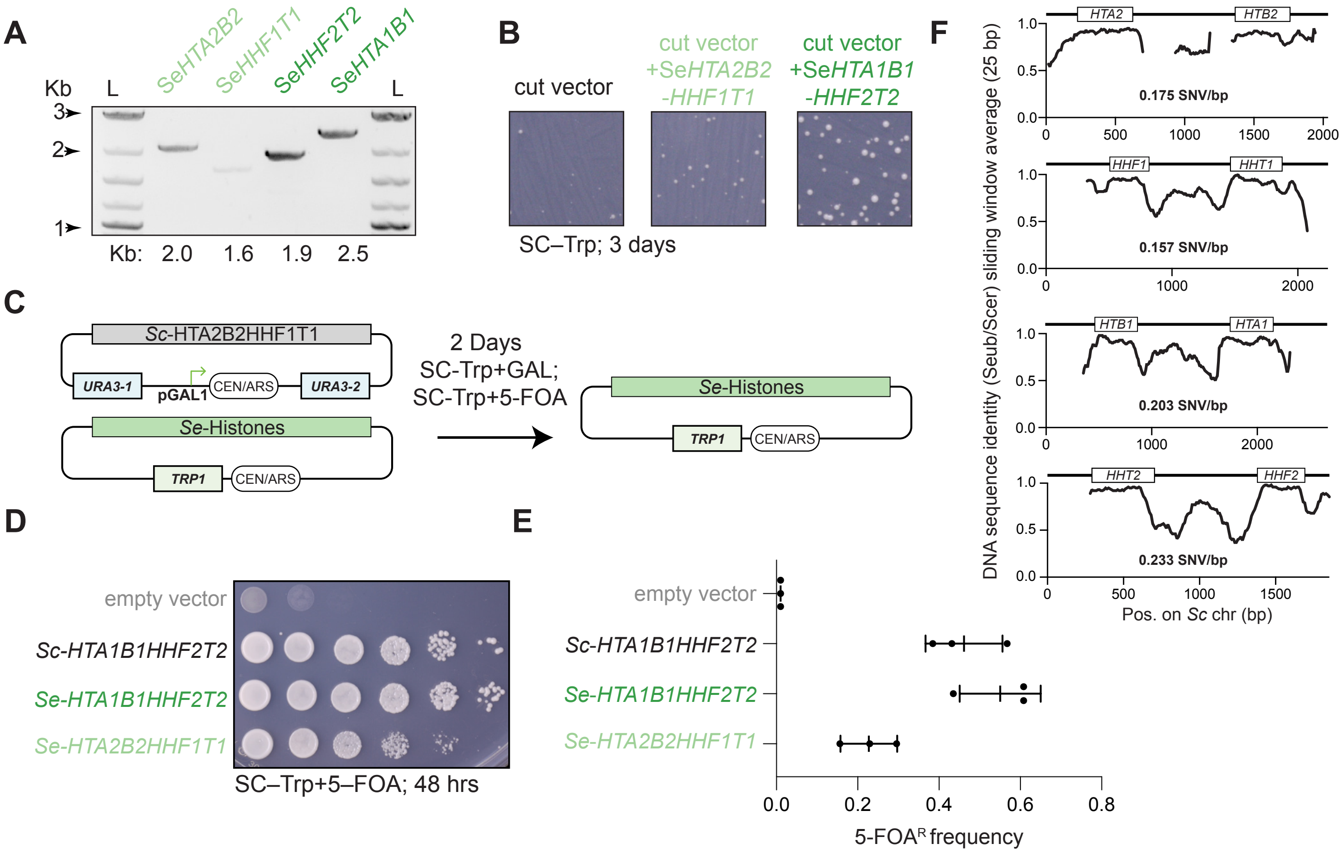

Figure S6. Epistatic interactions between canonical and non-replicative histones

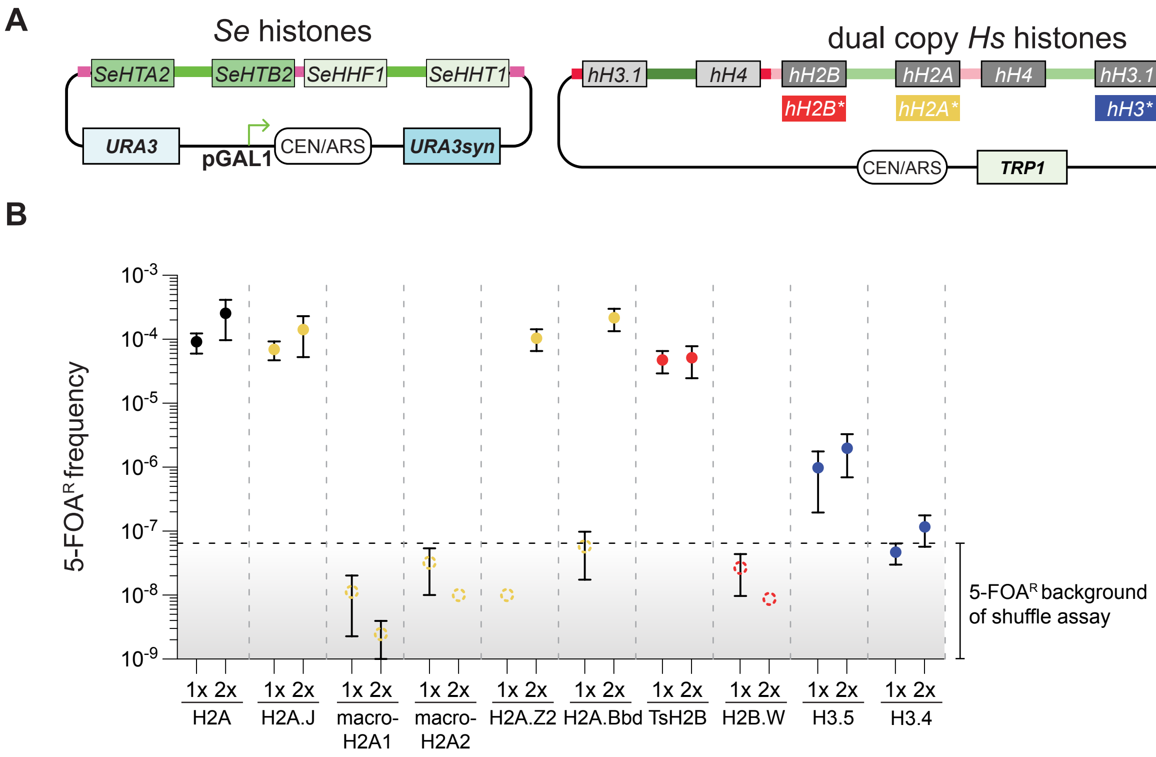

Figure S7. Dissecting the inviable residues of macroH2A1 histone fold

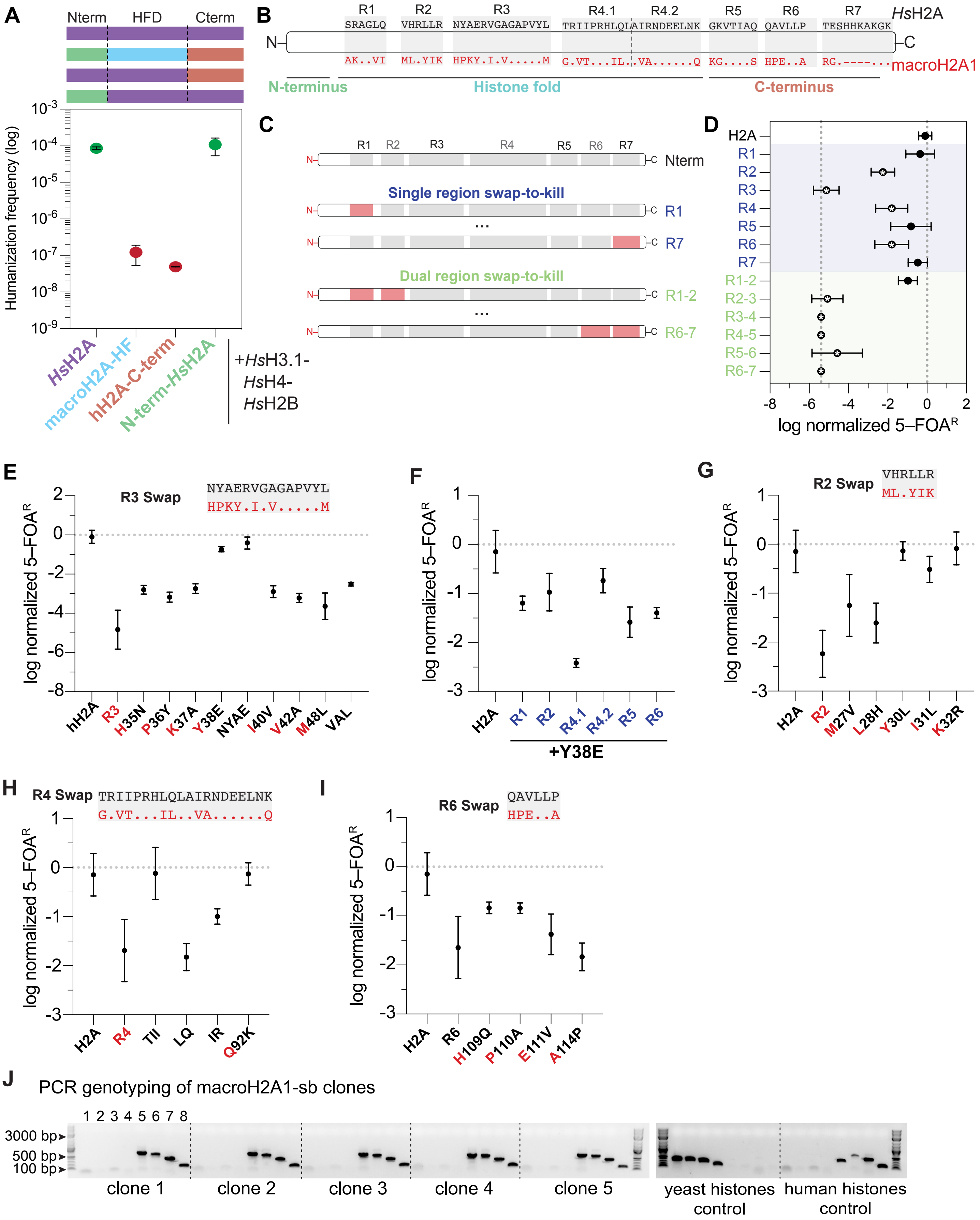

Figure S8. Cell size, doubling time and lag time of macroH2A1 humanized yeast

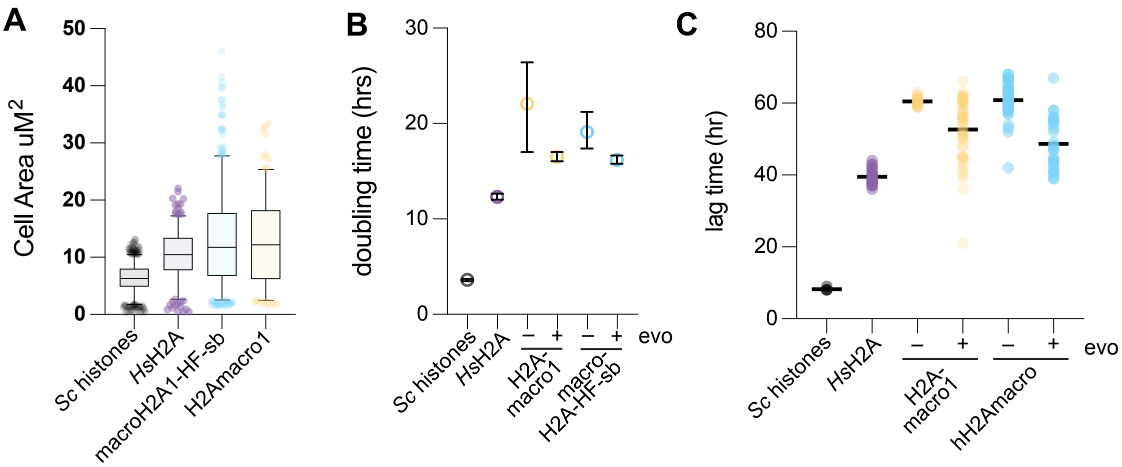

Figure S9. MNase digestions and MNase-sequencing analysis.

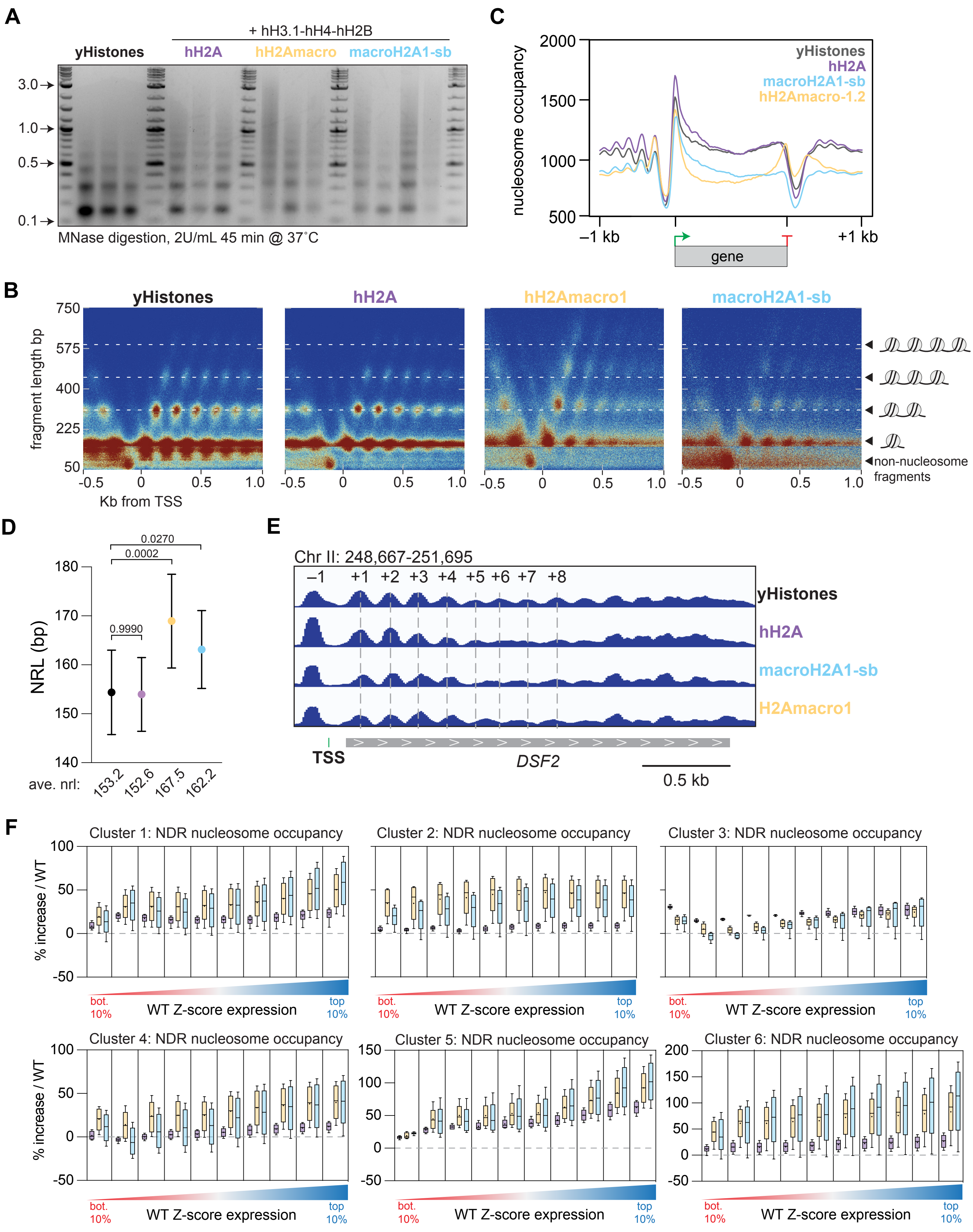

Figure S10. RNA sequencing in macroH2A1 humanized cells.

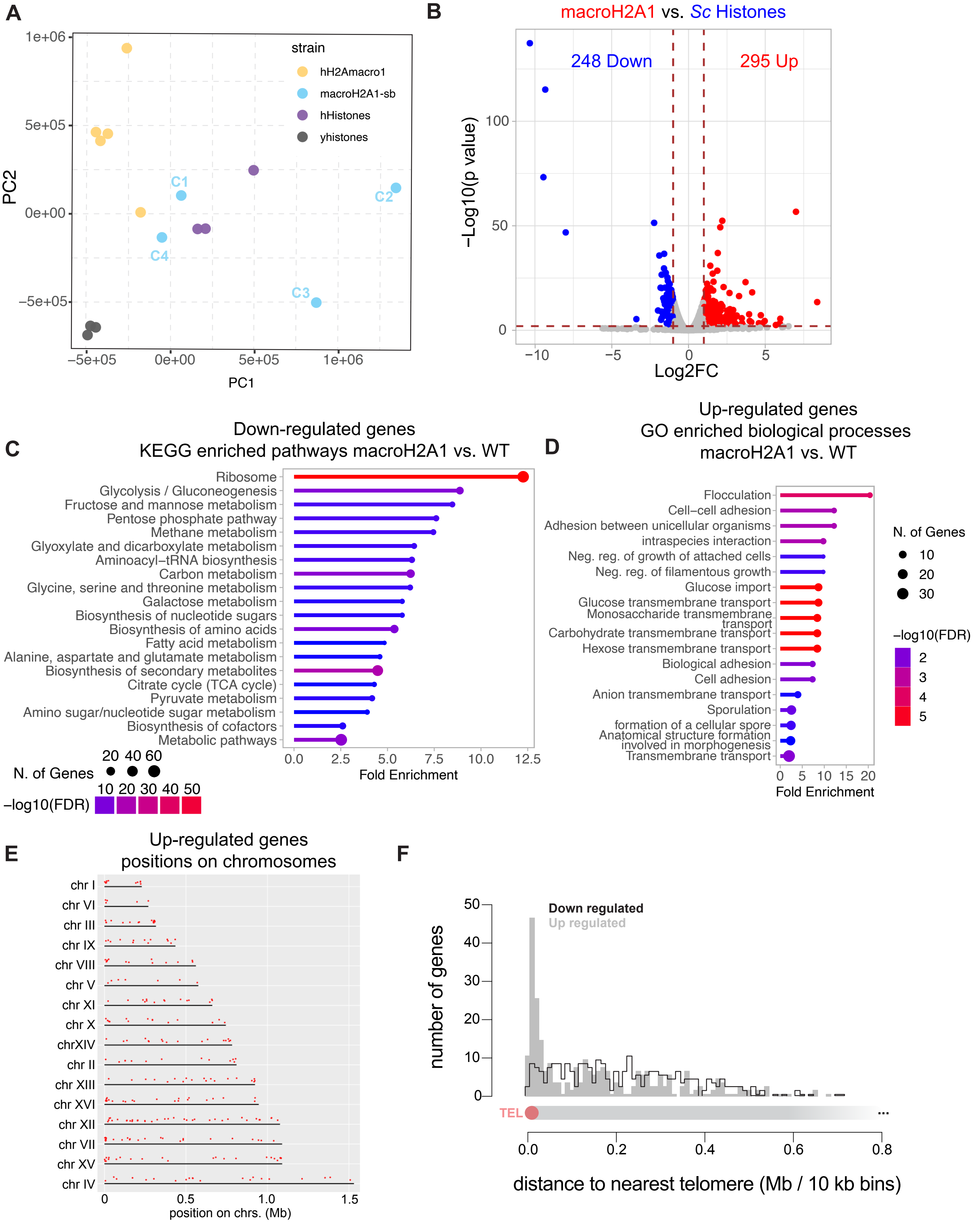

Figure S11. Global turn-down of protein translation inferred from MNase-seq and RNAseq in histone humanized yeasts

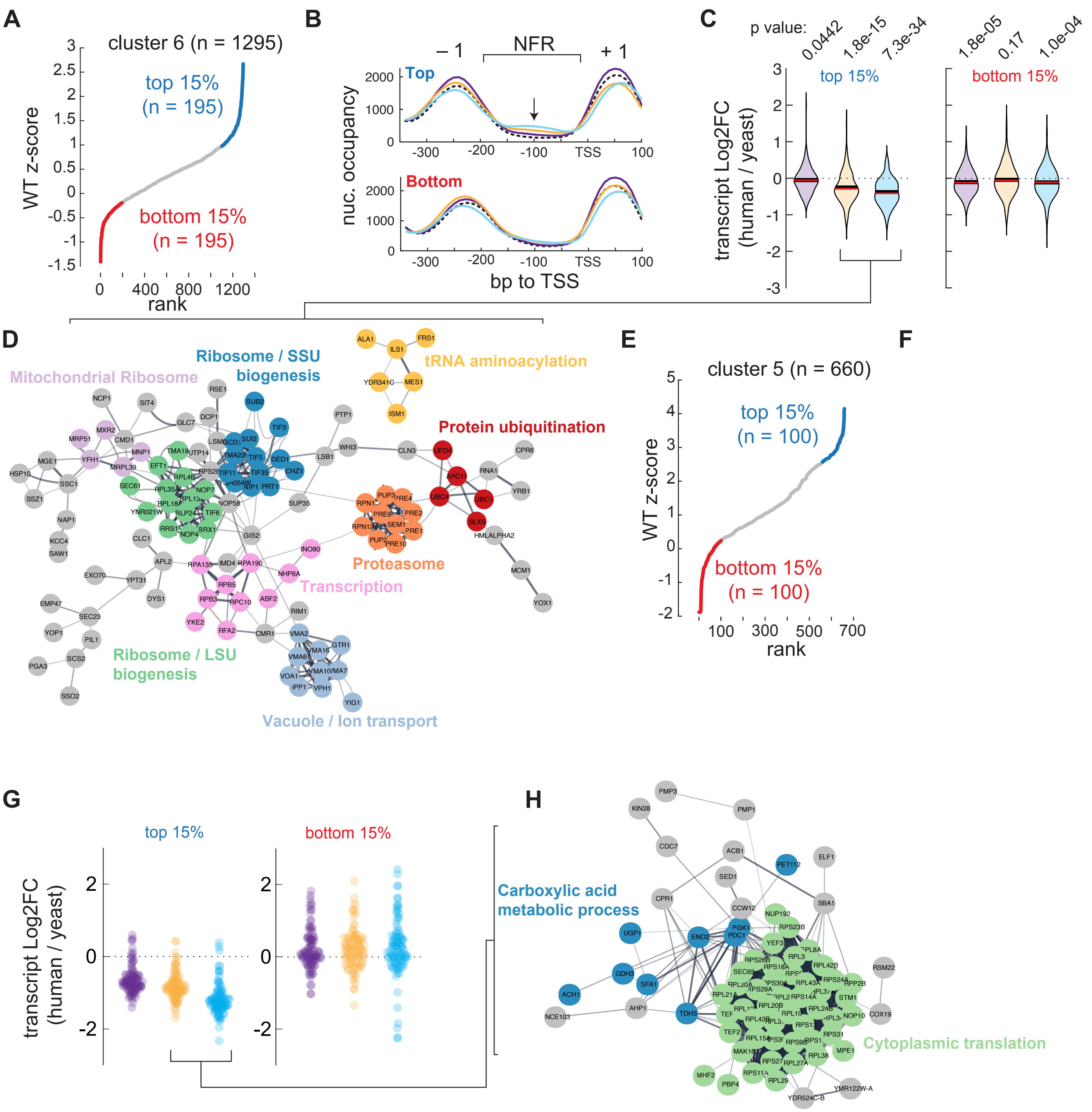

Figure S12. Relative nucleosome positioning downstream of the TSS

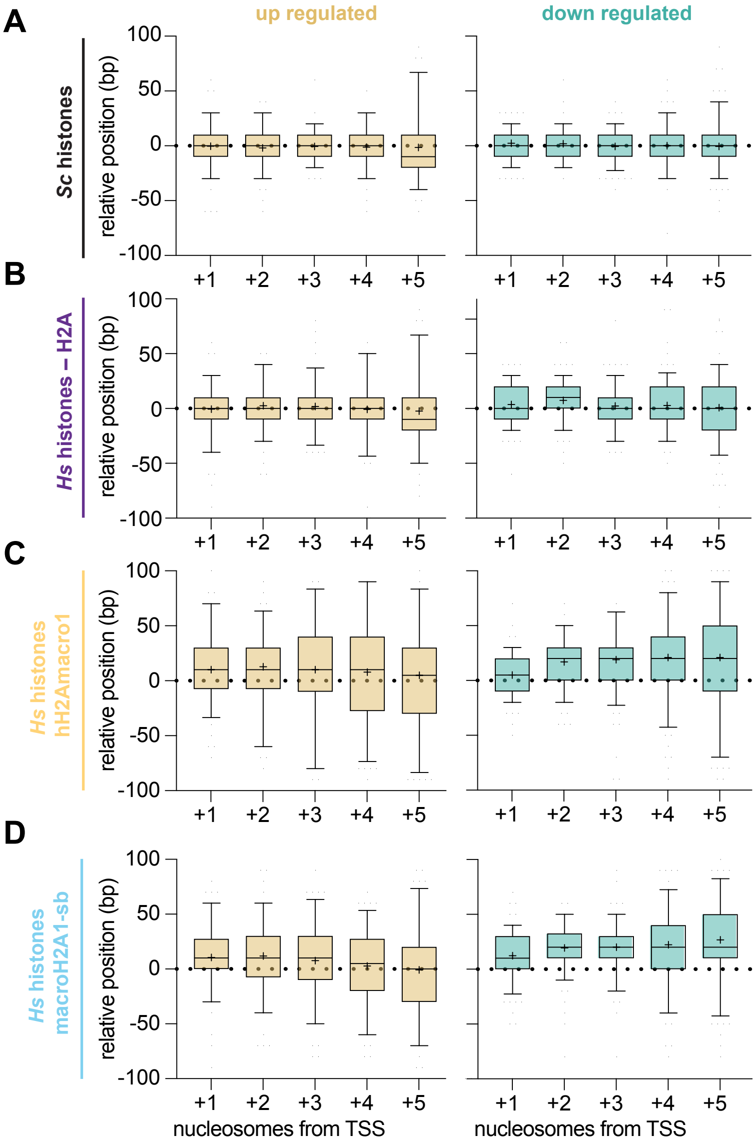

Figure S13. DNA shape features of up regulated genes near and far from telomeres

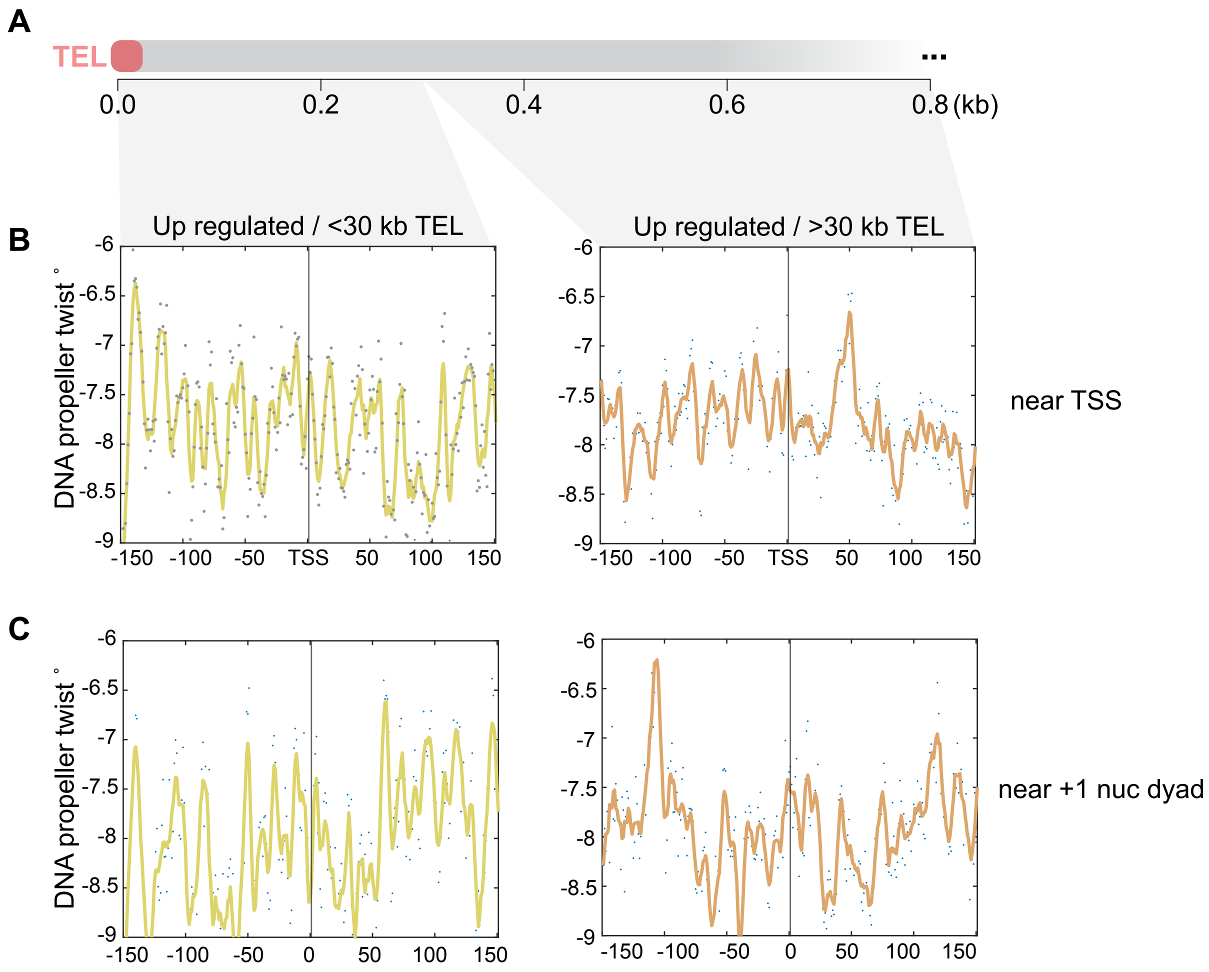

Figure S14. Clonal variation in genome stability of macroH2A1 humanized yeasts

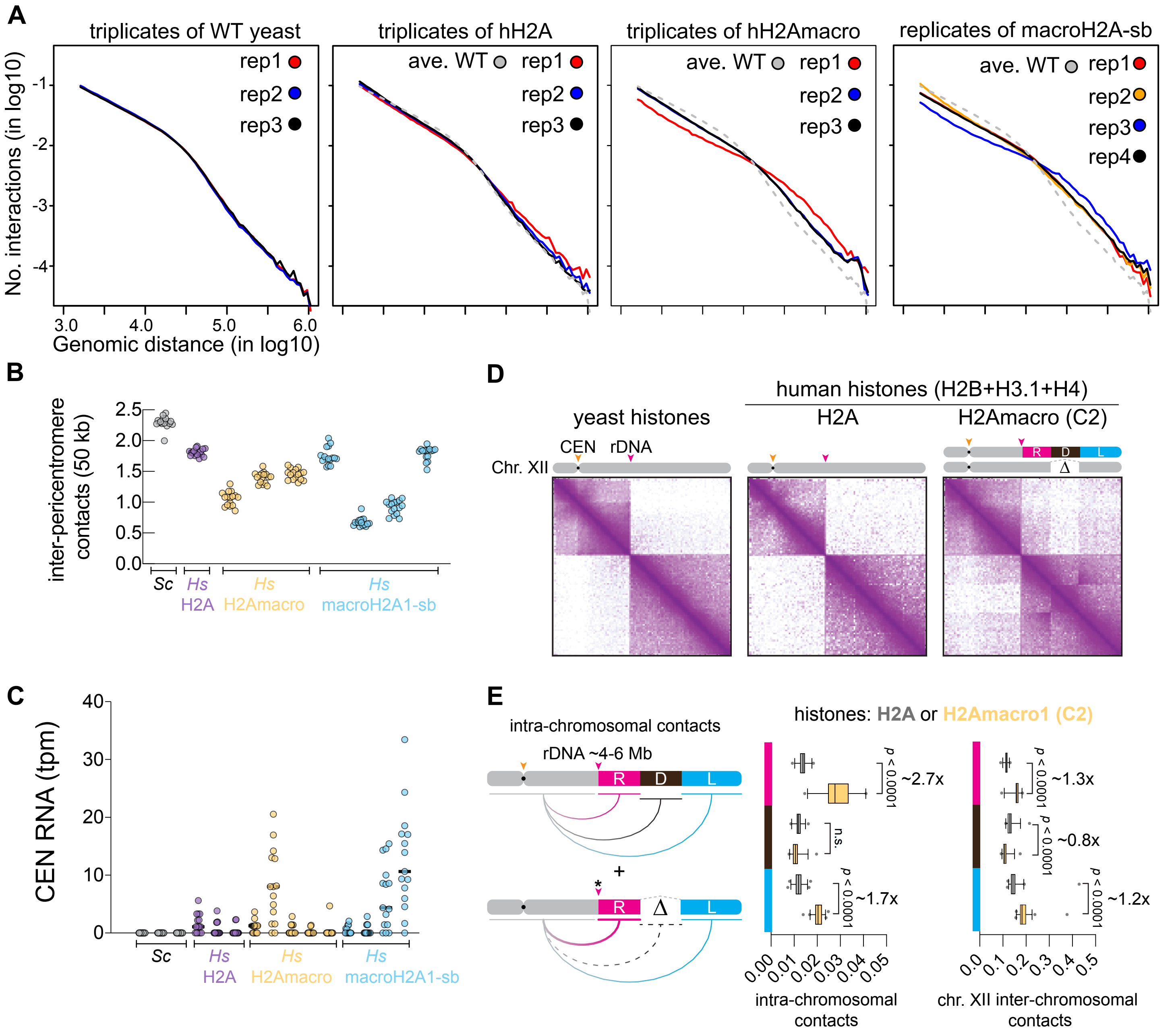

Figure S15. Clonal variation in genome stability of macroH2A1 humanized yeasts, continued

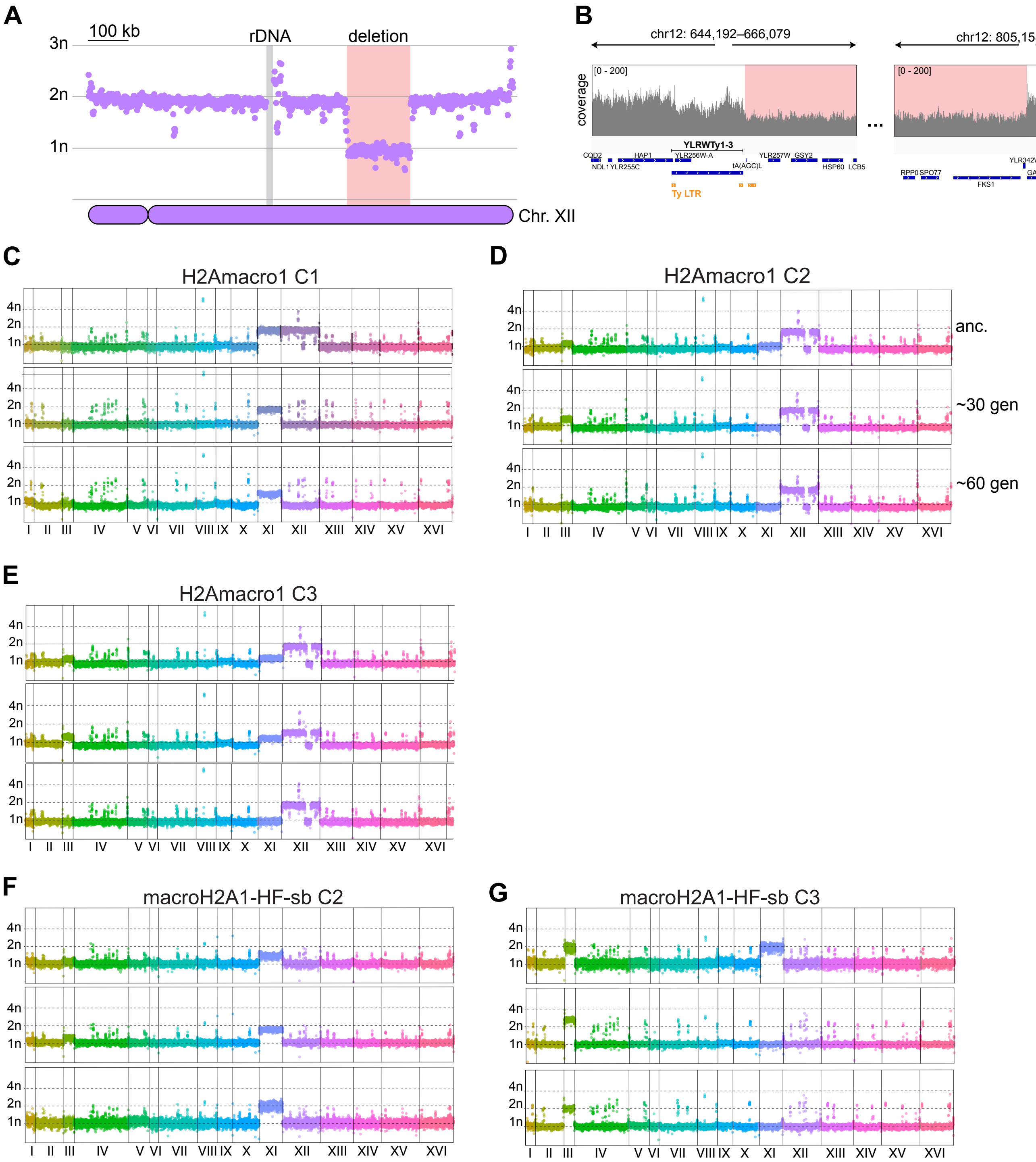
